# Beyond Level-1: Fast Inference of Generic Semi-directed Phylogenetic Networks

**DOI:** 10.64898/2026.04.17.719296

**Authors:** Nathan Kolbow, Joshua Justison, Claudia Solís-Lemus

## Abstract

Hybridization, introgression, and lateral gene transfer shape genomes across the diversity of life, but our ability to reconstruct these histories has been constrained by a methodological bottleneck: nearly all network inference methods are restricted to level-1 topologies, leaving more complex reticulate histories methodologically inaccessible. Here, we extend the widely used SNaQ method to scalably infer arbitrary binary, metric, semi-directed phylogenetic networks. Computational improvements yield substantial speedups in marginal composite likelihood evaluation, enabling genome-scale network inference under this framework for the first time. We systematically evaluate inference accuracy across networks of varying complexity, including cases where the true history falls outside the inferred search space, and find that SNaQ reliably recovers complex reticulate histories under diverse conditions, and still recovers meaningful information about hybridization events even when the full topology is not correctly inferred. Applied to the phylogeny of *Xiphophorus* (Poeciliidae), the method reveals a richer history of hybridization than level-1 approaches could capture, with network models that fit the data significantly better than previously inferred topologies. By enabling scalable inference beyond level-1 networks, our work facilitates the reconstruction of far richer reticulate histories from genomic data, bringing phylogenetic analysis closer to capturing the full network of life.

## 1 Introduction

Hybridization, introgression, and horizontal gene transfer are pervasive forces in evolution, shaping the genomes of plants, fungi, bacteria, and animals alike [1–3]. These processes leave distinct signatures in genomic data, but they cannot be represented in a phylogenetic tree: a model that assumes a strictly bifurcating, non-reticulate history. Phylogenetic networks extend trees by explicitly modeling horizontal evolutionary events, offering a more realistic framework for reconstructing the history of life.

Inferring phylogenetic networks from genomic data is substantially more challenging than tree inference. Under the multispecies network coalescent (MSNC) model [4–6], incomplete lineage sorting and reticulate processes are jointly modeled, enabling rigorous inference of reticulate histories in the presence of gene tree discordance. This framework provides a biologically realistic representation of evolutionary history, but it also introduces substantial computational challenges, particularly for large and complex networks [2, 7, 8]. Phylogenetic network estimation takes significantly longer than tree estimation primarily due to (i) a drastically larger topological search space (infinite without restrictions), and (ii) more complex likelihood equations. These challenges have traditionally been circumvented by employing maximum composite likelihood (MCL) methods: methods that approximate a network’s full likelihood by computing a product of simpler likelihoods [7, 9, 10]. This approach tends to be highly accurate and significantly faster than maximum likelihood approaches [8, 11].

Despite this progress, existing MCL methods, including SNaQ [7], have been restricted to level-1 networks, a class of phylogenetic networks whose biconnected components each contain at most 1 hybrid node (see Figure 1). Although, level-1 networks represent only a small subset of the topological complexities found in phylogenetic networks [12], the restriction remains popular in inference methods (*e.g.*, SQUIRREL Holtgrefe et al. [13], PhyNest Kong et al. [10], and NANUQ+ Allman et al. [14]) due to quartet identifiablility [7, 15] and computational tractability. Under the MSNC, the expected quartet concordance factors (qCFs) for any network can be expressed as a weighted average of the qCFs from the quartet trees induced by the network. In the case of level-1 networks, these weighted averages come in only a handful of closed forms that can be easily expressed and enumerated in software (see SI of [7]). This mathematical convenience, however, comes at a biological cost: the restricted level-1 class cannot represent many biologically realistic reticulate histories. For example, lineages that have experienced successive rounds of hybridization, hybrid taxa that have themselves hybridized with other species, or organisms with complex histories of introgression across multiple time points all violate the level-1 assumption. As a result, many biologically plausible reticulate evolutionary histories have remained outside the reach of statistical inference under the MSNC framework.

**Fig. 1:**
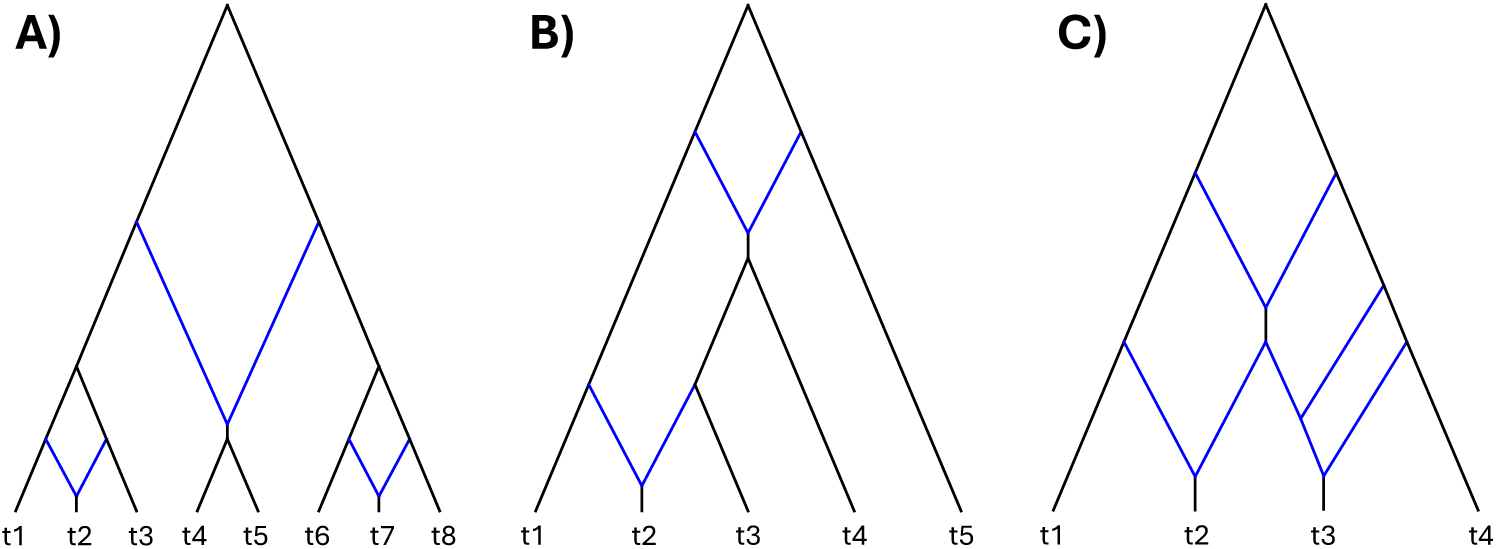
**Black lines: tree edges. Blue lines:** hybrid edges, parent to a hybrid node. **A)** Level-1, tree-child, and galled network. **B)** Level-2, tree-child, and not galled network. **C)** Level-4, not tree-child, and not galled network.

Here, we extend SNaQ to infer arbitrary semi-directed, metric phylogenetic networks by implementing a modified version of the recursive algorithm for computing expected qCFs under the MSNC given in Ané et al. [16]. For non-level-1 networks, qCF weighted averages grow rapidly in length and complexity, requiring recursive computation, substantially increasing the computational cost of likelihood evaluation. To offset this burden, we introduce gradient-based optimization with efficient forward differentiation and other implementation improvements that yield substantial runtime gains.

Guided by recent results showing that tree-child galled (TCG) networks have identifiable topologies under the MSNC [17], we focus our simulation study on this class. A semi-directed network is considered *strongly tree-child* [18] (hereafter simply “tree-child”) if all rootings of the network result in each hybrid node being parent to at least one tree node. A semi-directed network is considered *galled* [19] if each of its cycles contains at most one hybrid node (see Figure 1 for illustrative examples and Kong et al. [12] for further details). We evaluate inference accuracy both when the true network satisfies TCG conditions and when it does not.

Additionally, we introduce a flexible implementation that allows users to restrict the space of networks searched by SNaQ to a pre-specified domain of topological properties. This allows users to restrict inference to level-1 networks if desired, but also allows the inference of networks from any other class, networks with other arbitrary topological restrictions, or networks without topological restrictions. Theoretical and algorithmic details are described in Methods and implemented in the Julia package SNaQ.jl [20] (new version v2.0).

Finally, we reanalyze the evolutionary history of *Xiphophorus* (Poeciliidae), a genus that has been extensively studied and shows evidence of a complex reticulate history [7, 13, 21–26]. Using several search strategies, we evaluate network models inferred with our framework and show that relaxing SNaQ’s level-1 restriction yields significantly better-fitting models which reveal a more highly reticulated evolutionary history than previously inferred under the level-1 constraint.

## 2 Materials and Methods

### 2.1 Composite Likelihood Computation

Under the multi-species network coalescent (MSNC) model [4–6], the likelihood of a network *N* given a set of gene trees {*G_i_*} can be calculated exactly. The complexity of these computations, however, quickly becomes computationally intractable as the number of taxa in *N* increases and as the size of {*G_i_*} increases. To simplify these calculations, we can instead look at each of the quarnets (networks with 4 taxa) induced by *N*. For a given set of 4 taxa (say *a*, *b*, *c*, *d*), there are 3 possible unrooted gene tree topologies, which we can represent as the splits *ab*|*cd*, *ac*|*bd*, and *ad*|*bc*.

Using the MSNC, we compute the probability of each quartet topology given that the true underlying network induces the quarnet *Q*; we call these the *expected quartet concordance factors* (expected qCFs), denoted *E_N_* (*X_ab_*_|_*_cd_*), *E_N_* (*X_ac_*_|_*_bd_*), *E_N_* (*X_ad_*_|_*_bc_*), which we compute using a modified version of the recursive algorithm introduced in Ané et al. [16] (hereafter referred to as Algorithm 1). For each gene tree in {*G_i_*} containing *a*, *b*, *c*, *d*, we then record the frequency with which each quartet topology appears; we call these the *observed qCFs* (or *estimated qCFs* when {*G_i_*} consists of estimated gene trees), denoted *X_ab_*_|_*_cd_*, *X_ac_*_|_*_bd_*, *X_ad_*_|_*_bc_*. These observed qCFs follow a multinomial distribution with probabilities given by the expected qCFs.

From here, marginal composite log-likelihood (MCLL) and gradient equations follow:

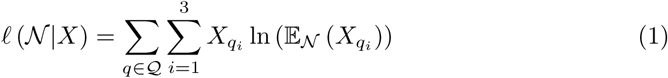

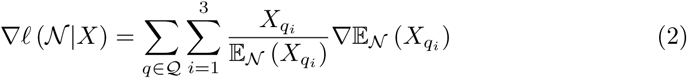

The original implementation of SNaQ.jl utilized these MCLL equations but restricted its search space to level-1 networks due to identifiability concerns [7]. This restriction improved computational feasibility due to the reduced size of searchable network space and the ability to hard code the small set of fixed forms that the MCLL equations could take. This original implementation also used these equations to optimize the parameters of a network via the gradient-free BOBYQA optimization algorithm [27]. Here, we loosen the level-1 restriction on the MCLL equations to allow computation of the MCLL for any network, and manually implement forward differentiation to compute the gradient of the MCLL with respect to each numerical parameter. We then use these gradients to optimize network parameters via L-BFGS [28]. Incorporating information from the gradient in this way allows for significantly more efficient parameter optimization [29].

One additional modification that improves the computational efficiency of this algorithm is to first assume, for each quarnet in *N*, that the quarnet is treelike. Computing expected qCFs for treelike quarnets does not require recursively reducing the network, yielding significant speedups for treelike quartets. If a quarnet is not treelike, we fall back to the full recursive algorithm. A second, more substantial modification is described below.

### 2.2 Gradient Computation

We further modify Algorithm 1 from Ané et al. [16] by storing the equations to compute a network’s expected qCFs in a computation graph. This allows us to efficiently recompute expected qCFs and their gradients for different parameter values without needing to make repeated, costly calls to this recursive algorithm. To this end, we construct a directed, multifurcating computation graph as we recursively iterate through Algorithm 1 (see an example of such a graph and its corresponding quartet in Figures S1 and S2, respectively). Every such graph has root node “1” and multifurcates outwards depending on the specific structure of its associated quarnet’s topology. The expected qCF of a network can be computed from such a graph by summing the products of all unique paths from the root node “1” to each of the graph’s leaf nodes. Each leaf node in these graphs is a vector with three elements, corresponding to each of the three values that comprise a qCF.

While every possible qCF equation under the MSNC can be contained in such a graph, each node of the graph can only take the form of one of 11 unique formulas. These formulas correspond to each of the return values in Algorithm 1 from Ané et al. [16] and are enumerated along with their gradients with respect to each type of parameter (edges *t_i_* or inheritance proportions *γ_i_*) in Table S1. Our implementation also lets the user specify the inheritance correlation parameter *ρ* [30] to control the inheritance correlation throughout the network, but this parameter is not optimized. Efficient computation of the expected qCF gradients takes advantage of two key insights. First, each unique parameter in the network’s topology appears in at most one node in each path from the root to any tip. This is a property of the recursive deconstruction of the network that occurs throughout Algorithm 1 from Ané et al. [16]: at each step of the algorithm, whenever a parameter contributes to the expected qCF, its corresponding topological component is removed from the quarnet. Thus, it will not appear again in the same path in the computation graph. Second, a quarnet’s expected qCFs are constructed from a sum of products, as described above. These products are represented in the computation graph as the unique paths from the root to each tip. Taken together, these details reveal that computing the gradient of each product (i.e., path in the computation graph) with respect to some parameter *t_i_* or *γ_i_* requires simply computing the derivative of the component of the product that contains *t_i_* or *γ_i_*. Thus, to compute the derivative of a given product with respect to some parameter *t_i_* or *γ_i_*, we only need to find the node in the product’s corresponding path in the computation graph that contains *t_i_* or *γ_i_* and then compute the derivative of that node. Or, if the parameter does not appear in any nodes in a given path, the corresponding derivative is 0. Consequently, we can compute all three expected qCFs for a given quarnet along with their derivatives with respect to all of a network’s parameters in a single walk of the computation graph with minimal overhead and bookkeeping.

Additionally, we leverage the fact that qCFs for a given 4-taxon subset always sum to 1 and their gradient components always sum to 0 to further improve performance by only computing two expected qCFs according to the computation graph (and their respective gradients). We use these values to determine the third qCF (or final gradient) with simple arithmetic.

The primary remaining runtime and memory cost to this algorithm comes from repeatedly copying *N*, reducing it to a 4-taxon subnetwork, then further recursively reducing the network. Future work could vastly improve the efficiency of these computations further by computing expected qCFs in-place without copying or modifying *N* in software.

### 2.3 Simulation Study

To evaluate the performance of SNaQ under different topological restrictions, we simulated sequence data, estimated quartet concordance factors, and inferred networks across different network spaces. Guided by recent results showing that tree-child galled (TCG) networks have identifiable topologies under the MSNC [17], we focus our simulation study on this class, but we additionally expanded the simulation space to features beyond topologically identifiable networks. First, ground truth networks were simulated from the space of tree-child and galled (TCG) networks, tree-child but not galled (TCNG) networks, and neither tree-child nor galled (NTCNG) networks. To assess the robustness of SNaQ to model violations, network inference was always restricted solely to the space of TCG networks. These networks were simulated with the SiPhyNetworks R package [31]. To adjust the level of incomplete lineage sorting (ILS) in the ground truth networks, edge lengths were multiplied by a constant such that the average edge length in each network was fixed (2.0 coalescent units for low ILS, 0.5 for high ILS). For each network, 100-10,000 gene trees were then generated under the MSNC with the PhyloCoalSimulations.jl Julia package [30] and multiple sequence alignments (MSAs) were simulated from each gene tree using seq-gen [32] under the HKY85 substitution model [33]. Estimated gene trees were inferred with IQ-TREE v3.0.0 [34] without specifying the underlying model of evolution. We additionally used true gene trees, low ILS, 10,000 loci, and 20 taxa to evaluate the performance of estimating the non-TCG networks in an ideal inference setting.

#### 2.3.1 Topological Accuracy

To quantify the error in the inferred networks, we computed the semi-directed hardwired cluster dissimilarity (SDHWCD) between the true network and the inferred network of each simulation. The hardwired cluster dissimilarity (HWCD) [35] is a dissimilarity measure computed between two rooted networks. The SDHWCD is a generalization that is computed between two semi-directed networks, and is defined as the minimum HWCD that is attainable by any re-rooting of the two semi-directed networks that does not introduce directional conflicts. Note that this is only a true distance metric for certain networks (*e.g.* normal level-1) and thus we refer to it as a dissimilarity.

Additionally, we computed the negative composite log-likelihood (NCLL) score of each inferred network and compared it to that of the true network. These measures allow us to simultaneously evaluate whether SNaQ is finding networks that sufficiently explain the data (low NCLL scores) and whether the networks inferred by SNaQ are topologically similar to the true network (low SDHWCDs).

#### 2.3.2 Numerical Parameter Estimation

To assess SNaQ’s ability to accurately infer a network’s numerical parameters, namely edge lengths *t* and inheritance probabilities *γ*, we evaluate the absolute error (AE) in *γ* estimates (Equation 3) and the relative absolute error (RAE) in *t* estimates (Equation 4). Given true and inferred networks *N* and *N̂* with edges *E^N^* and *E^N̂^*, respectively, we denote true and inferred edge length and inheritance probability parameters as 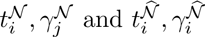, respectively. Then, the equations for AE and RAE are given by:

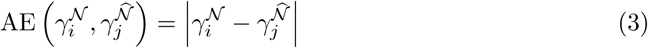

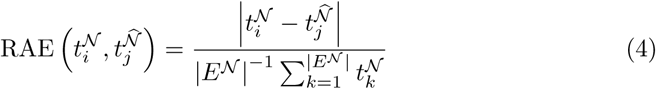

AE is clear and interpretable when evaluating *γ* estimates due to their restriction to the interval [0, 1], but AE can be misleading when evaluating edge length estimates. Edge lengths take values in [0, ∞), so an AE of 1.0 is relatively negligible when 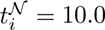, but is severe when 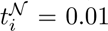. Thus, errors in edge length estimation are more meaningful and interpretable when weighted as in Equation 4. We weight the errors by the average edge length in the true network rather than 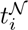 to avoid individual 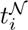 values that are particularly small or large artificially inflating or deflating the RAE value, respectively.

#### 2.3.3 Accuracy in Identifying Hybrid Ancestry

To evaluate the accuracy of reconstruction of reticulate ancestry in our simulations, we implement a weighted *F*_1_ score on the true and inferred hybrid ancestries. The *F*_1_ score (Equation 7) is a metric that balances precision (positive predictive value, Equation 5) and recall (true positive rate, Equation 6). This metric takes values in [0, 1] where a value of 1 indicates a perfect match between the true and inferred sets *S_true_* and *S_inferred_*, respectively. Conversely, a value of 0 indicates that there is no overlap between the true and inferred sets. We utilize this metric because it provides a natural penalty when taxa are incorrectly placed as descendants to hybrid nodes in the inferred network.

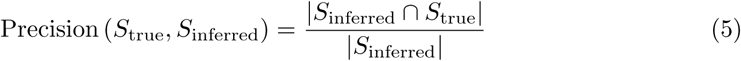

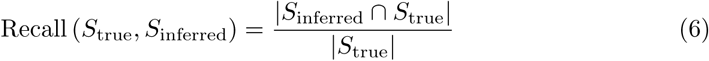

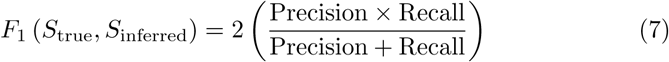

For a given pair of true and inferred networks (*N* and *N̂*, respectively), let *h* be the total number of hybrid nodes in the true network. Then, following the notations and definitions of [16], call 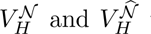 the sets of hybrid nodes belonging to each network. We then denote the set of descendants of a given hybrid node (*V*) as *D*(*V*). This set is in general unambiguous due to the semi-directed nature of the networks *N* and *N̂*. It follows that the *F*_1_ score between a single true and inferred hybrid node is given by:

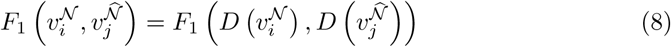

where 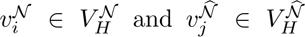. Call *P* the set of all permutations of the set {1, 2,…, *h*}, then the so-called weighted *F*_1_ score between the true and inferred networks is:

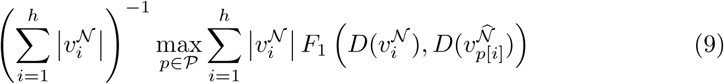

Weighting this metric by the size of each true hybrid node is important to avoid hybrid nodes with fewer descendant taxa having as much impact on the metric as hybrid nodes with many descendant taxa. Taking the maximum over *P* in this manner avoids the difficult problem of matching nodes in non-isomorphic graphs.

We used the semi-directed hardwired cluster dissimilarity (SDHWCD) metric to evaluate the topological similarity between true and inferred networks. Note that a SDHWCD value of 0 does not strictly imply isomorphism, so networks with SDHWCD values of 0 may have weighted *F*_1_ scores below 1.

When inferring a network with SNaQ, the maximum number of hybrid nodes allowed in the network (*h_max_*) must be specified a priori. In our simulations, we set *h_max_* = h (true value), but, in a small number of cases, the network inferred by SNaQ had *ĥ* < *h* hybrid nodes. In this case, for each *j* = *h* + 1,…, *h_max_*, we defined 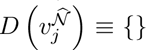 to represent the missing nature of this hybrid node.

### 2.4 Phylogeny of *Xiphophorus* (Poeciliidae)

To evaluate our approach on empirical data, we reanalyze the evolutionary history of *Xiphophorus* (Poeciliidae), a genus of platyfishes and swordtails whose relationships have been extensively studied and show strong evidence of reticulate evolution [7, 22].

To illustrate the versatility of SNaQ’s new framework, we infer networks from the following search spaces:

1. Networks that are both tree-child and galled (TCG-space)
2. Completely unrestricted (U-space)
3. Completely unrestricted search with the TCG-space results used as the starting point of each run (TCGU-space)
4. Heavily restricted by clade with further details below (C-space)

*Xiphophorus* (Poeciliidae) can be divided into four clades: northern swordtails (NS), southern swordtails (SS), northern platyfishes (NP), and southern platyfishes (SP) [22]. C-space consists only of tree-child and galled networks with the following property: any pair of taxa with a shared parent node (often referred to as “sisters”, “siblings”, or “cherries”) must belong to the same clade. This restricts the search space to networks where close, treelike sibling relationships must be monophyletic without uniformly disallowing polyphyletic relationships. The best-fitting level-1 networks from the original SNaQ Solís-Lemus and Ané [7] (and the best-fitting TCG-space networks inferred here; more in Results) exhibited the C-space property, so we theorized that only considering such networks could drastically accelerate inference by reducing the number of inferior topologies that are analyzed.

For each search space, networks were inferred with a sequentially increasing maximum number of hybrid nodes *h_max_* = 0, 1, 2,…, 7. Inference in TCGU-space began at *h* = 2 because TCGU-space is equivalent to U-space for *h <* 2. For each value of *h_max_* in each search space, networks were inferred across 100 independent runs. Starting topologies for *h* = 0 in each case were a tree inferred with ASTRAL v5.7.1 [36].

We utilize the same 1,183 gene trees inferred by MrBayes 3.2.1 [37] and quartet concordance factors inferred by BUCKy as in [7, 22]. Additionally, we compare our results to the original level-1 network inference results (L1-space) from SNaQ.jl v1.0 [7]. These networks were inferred with 0 ≤ *h* ≤ 5.

To select the best-fitting network in each search space, we used the data-driven slope estimation (DDSE) model selection approach implemented in the CAPUSHE R package [38]. DDSE calibrates a loss penalty by assuming that the NCLL scores of the most complex models are roughly linear with respect to the number of parameters in the model, then selects the model with the best penalized NCLL score. A visual inspection of Figure 6 validates this linearity assumption.

## 3 Results

### 3.1 Topological Accuracy

Network topological similarity improves under simpler evolutionary conditions and with increased data (Figure 2). Networks with fewer taxa and lower levels of ILS were inferred more accurately, and accuracy further increased as both the number of loci and the sequence length grew. Counterintuitively, when the simulations had only limited data (100 loci in particular), some estimated networks had better NCLL scores than the true generating network, and thus were preferred by SNaQ, despite having poor topological accuracy (high SDHWCD). This effect was especially pronounced for datasets with 10 taxa and only 100 or 1,000 loci, where nearly all inferred networks scored higher NCLL values than the true network regardless of their accuracy.

**Fig. 2:**
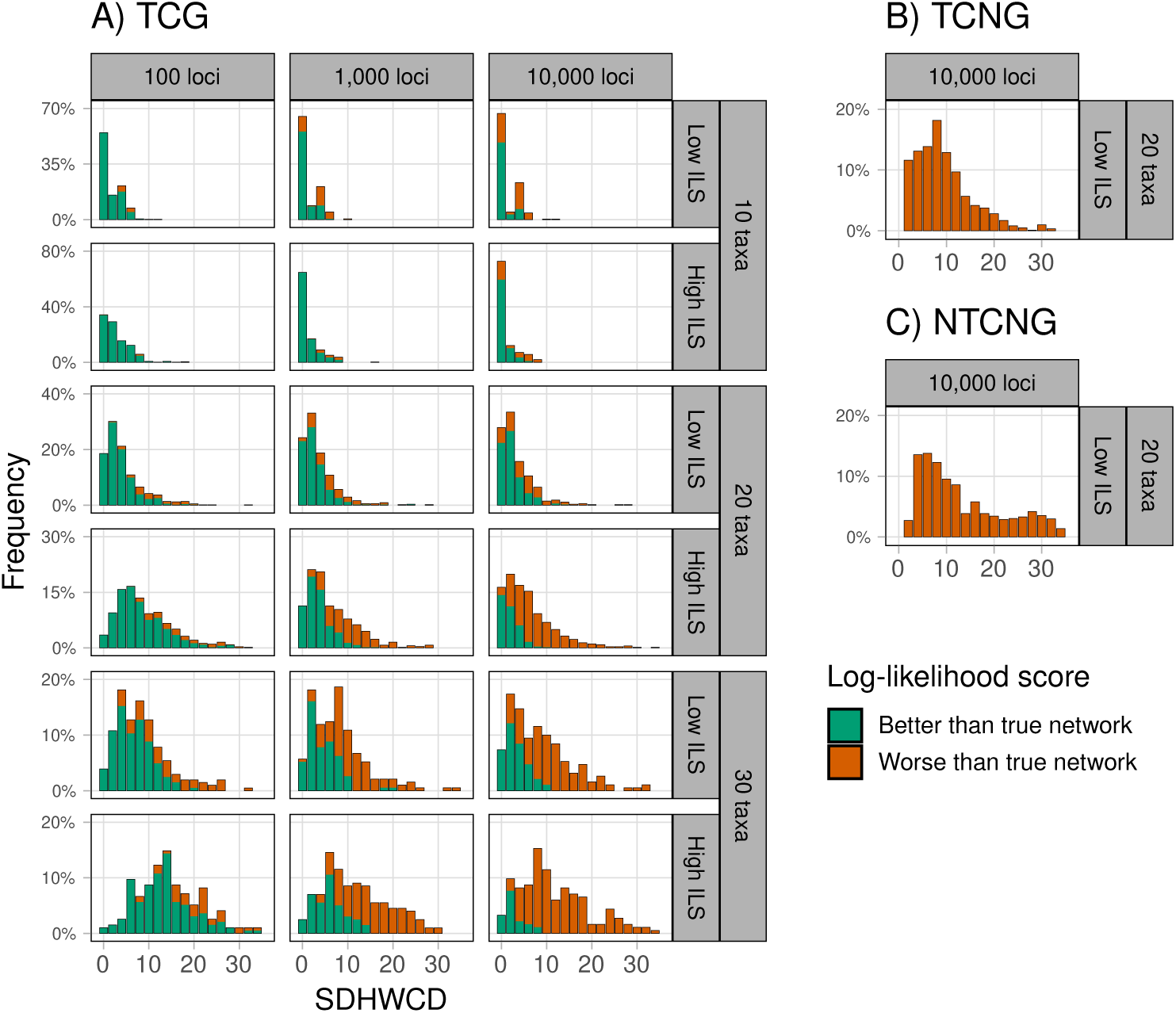
Distribution of semi-directed hardwired cluster dissimilarities (SDHWCD) between the true network and the network inferred by SNaQ across different simulation settings. Each histogram bar is filled *green* proportionally to how many inferred networks in each bucket yielded better negative composite log-likelihood (NCLL) scores than the ground truth network. The remaining proportion of networks, filled with *orange*, had worse NCLL scores than the ground truth network. Ground truth networks belonged to different network spaces, including **A) TCG** (tree-child and galled), **B) TCNG** (tree-child and not galled), and **C) NTCNG** (not tree-child and not galled), but all inferred networks were restricted to TCG-space.

Simulations in which the true network was either TCNG or NTCNG yielded substantially worse SDHWCD values than simulations with TCG networks. In fact, perfect recovery (SDHWCD = 0) was impossible in these cases because the estimated networks were restricted to the TCG space. Although it is theoretically possible for a TCG network to achieve a higher NCLL score than the true network under these conditions, this did not occur in our simulations.

Distributions of SDHWCD values were highly dependent on individual replicates in TCNG and NTCNG cases (Figures S10-S11). The SDHWCDs of some networks in these classes were concentrated heavily around 2 (the minimum possible, and therefore the best possible, in this case), while the SDHWCDs of others were highly concentrated around or above 30, and the SDHWCDs of others still were spread between these values. In the NTCNG case, we found a strong correlation between the SDHWCD of the best inferred network and the SDHWCD of the TCG network that is as similar as possible to the true NTCNG network (Figure S12). We found no such correlation in the TCNG case.

### 3.2 Numerical Parameter Estimation

Of the 125 total unique topologies in the TCG-space simulations, 11 were identified as outliers and removed from *γ* and *t* parameter estimate analyses due to extremely high AE and RAE values. Of the outlier networks, four contained 10 taxa, six contained 20 taxa, and one contained 30 taxa. All but two of these networks had the recurring topological feature of having at least one hybridization with exactly two descendants (see Figure S4 for two examples). The reamining two outlying networks contained 10 taxa with 1 hybrid (Figure S5) and 20 taxa with 2 hybrids (Figure S6). Some inlier networks contained this topological feature, but they all either had another reticulation with many descendants or several more reticulations each with one or two descendants, reducing the average error across the network. Of the two outlier networks without this topological feature, both had theoretically identifiable topologies and parameters, but parameter estimates for two edges in the 10 taxa, 1 hybrid network (Figure S5) were highly unstable. For example, the true topology along with the relevant true parameter values of this network is given in Figure S5(b), but, with ground-truth data, a near-perfect NCLL score can be achieved by setting *t̂*_1_ = 25 (*t*_1_ = 2.444) and *t̂*_2_ = 0.963 (*t*_2_ = 0.772), *t̂*_3_ = 0.600 (*t*_3_ = 0.483) and *γ̂* = 0.432 (*γ* = 0.349). Notably, 25 is the maximum edge length SNaQ can infer, because at this point coalescent probabilities are indistinguishable from 1. The NCLL score with these parameter adjustments is distinct from that of the true network in a theoretical sense, but it is close enough that it presents a clear challenge for numerical optimization software. We were unable to identify any features of the last outlying network (Figure S6(d)) that were not present in the inlying networks.

The accuracy of parameter estimates tended to improve with increased data, but were not noticeably improved under simpler evolutionary conditions (lower ILS) except when only 10 taxa were present (Figure 3). This was the case regardless of whether SDHWCD was zero or not, which lends nuance to the prevailing wisdom that inference is strictly more difficult in higher ILS conditions. When the network’s topology was correctly inferred (SDHWCD = 0), parameter estimates improved as the number of taxa in the network grew; this could be the result of the number of qCFs that a network has scaling quartically with the number of taxa in the network (*O*(*n*^4^)) whereas the number of parameters in a network scales roughly linearly (*O*(*n*)).

**Fig. 3:**
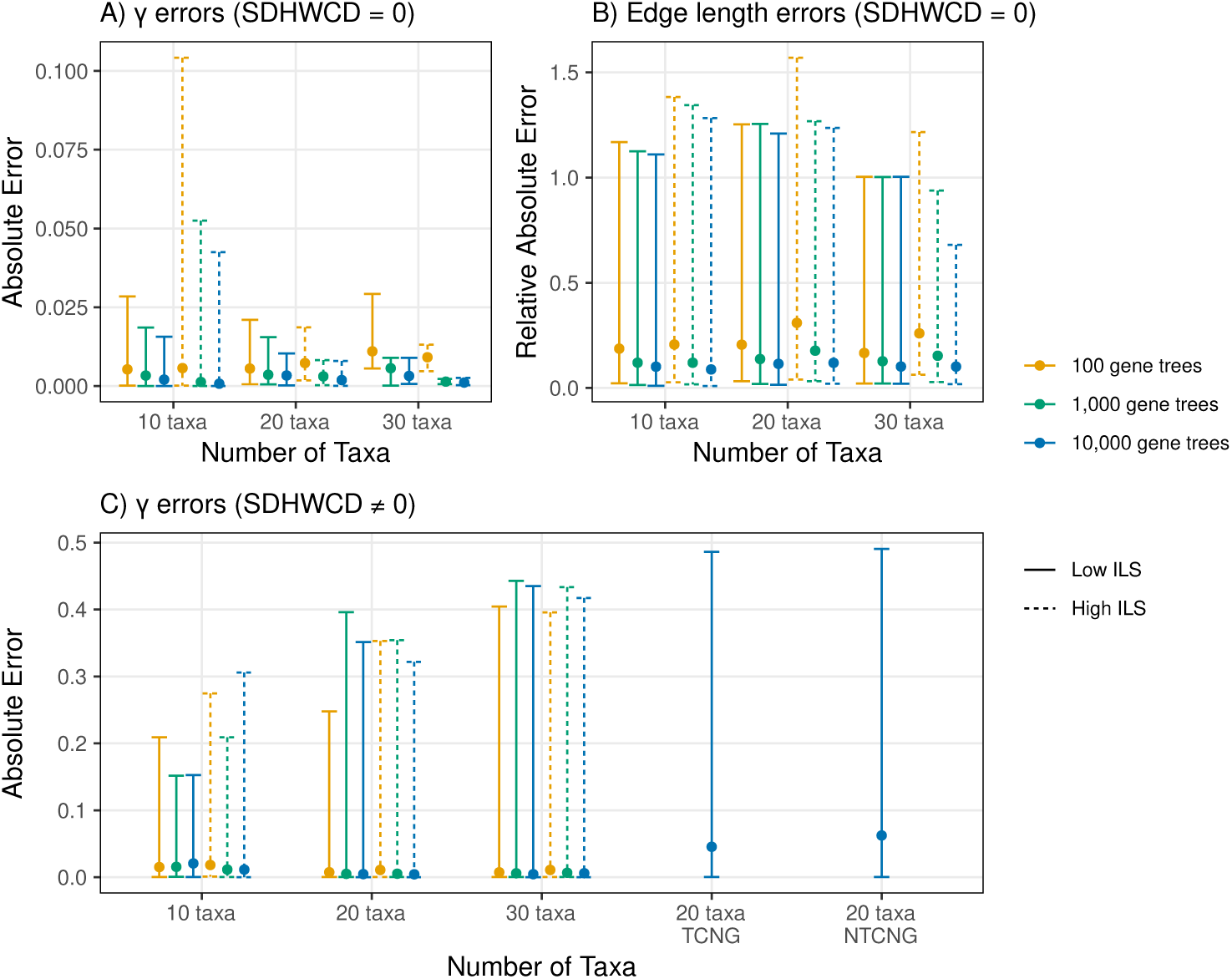
Distributions of observed parameter estimation errors across simulation parameters. Lines: 95% observed confidence regions (*q*_0.025_*, q*_0.975_) for inference errors. Points: median observed inference errors. **A)** Absolute error (AE) in inferred inheritance parameters (*γ*) when the semi-directed hardwired cluster dissimilarity (SDHWCD) between the true and inferred networks was 0. Four outlier networks were removed from each of the 10 taxa and 20 taxa simulations. **B)** Relative absolute error (RAE; Equation 4) in edge length parameters when the SDHWCD between the true and inferred networks was 0. **C)** Absolute error (AE) in inferred inheritance parameters when the SDHWCD between the true and inferred networks was not 0. *γ* parameters were matched between networks in this case using the process used to compute *F*_1_ scores in Equation 9 described in Section 2.3. Eleven outlier networks out of 125 total networks were removed from these results.

When the simulation model matched the search space (TCG), numerical parameter estimates were accurate; median AEs for *γ* were near zero while median RAEs for *t* were no higher than 0.25 and approached 0 as the number of gene trees used increased (Figure 3). For TCG simulations, when the network’s topology was correctly inferred, the distributions of *γ* estimates were tight around the median in most cases. The only scenario where this was not the case was for 10 taxa networks with high ILS, likely due to the simultaneously high amounts of discord and lower relative abundance of quartet data. When SNaQ failed to infer the correct network topology, median *γ* estimates were still near-zero, but the variance in *γ* estimates dramatically increased relative to when topolgies were correctly estimated. Variance around branch length and gamma estimates when the network’s topology was not correctly inferred was larger both as the size of the network grew and when the model was misspecified.

### 3.3 Accuracy in Identifying Hybrid Ancestry

To evaluate the accuracy of estimated hybrid clades, we computed the weighted average of *F*_1_ scores between the multisets of hybrid descendants in the true network and the inferred network.

Hybridization events were accurately recovered when the true network was a part of the TCG class (Figure 4). F_1_ scores were consistently high in these simulations, though they declined slightly as the number of hybrid nodes increased. Increasing the number of gene trees used for inference marginally improved F_1_ scores on average. Notably, this trend was independent of the SDHWCD values of the inferred networks. In contrast, simulations in which the true networks fall outside the TCG class (TCNG or NTCNG) yielded markedly lower F_1_ scores, as expected when the exact hybrid descendancy structure lies outside the space of inference.

**Fig. 4:**
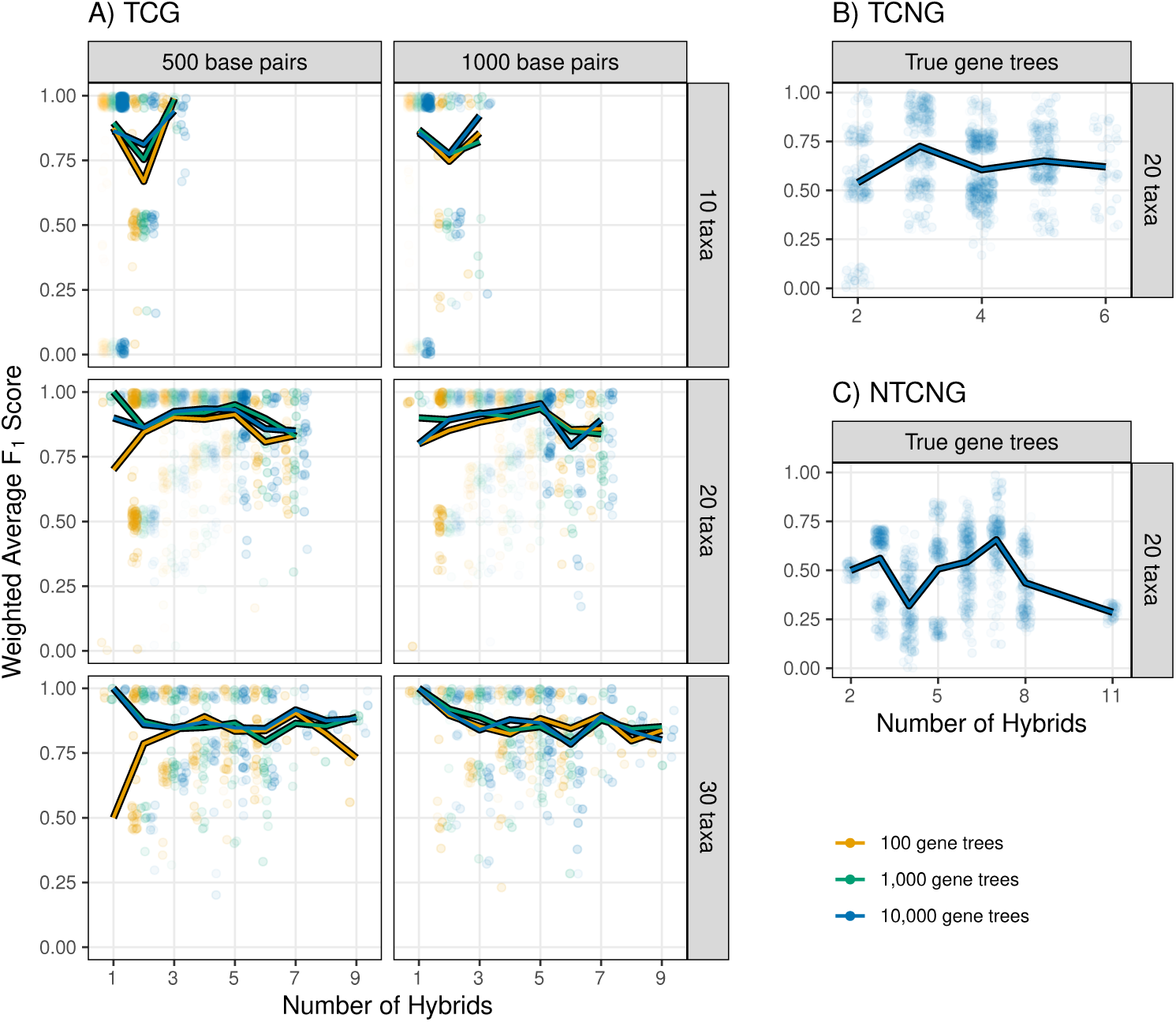
Weighted average *F*_1_ scores comparing the descendants of each hybrid node in the true network and the descendants of that node’s corresponding match in the inferred network (Equation 9). **TCG** (tree-child and galled), **TCNG** (tree-child and not galled), and **NTCNG** (not tree-child and not galled) refer to each of the spaces from which the ground truth networks were simulated. All inferred networks belonged to the TCG space.

### 3.4 Computational Improvements

Composite log-likelihood equations for arbitrary phylogenetic networks are not straightforward, particularly as the number of reticulations increases. Gradient-based parameter optimization, however, has significantly improved the speed of parameter optimization and thus expanded the scale of networks that can reasonably be inferred with our extended framework (Figure 5). Runtimes can be found in Figure S13.

**Fig. 5:**
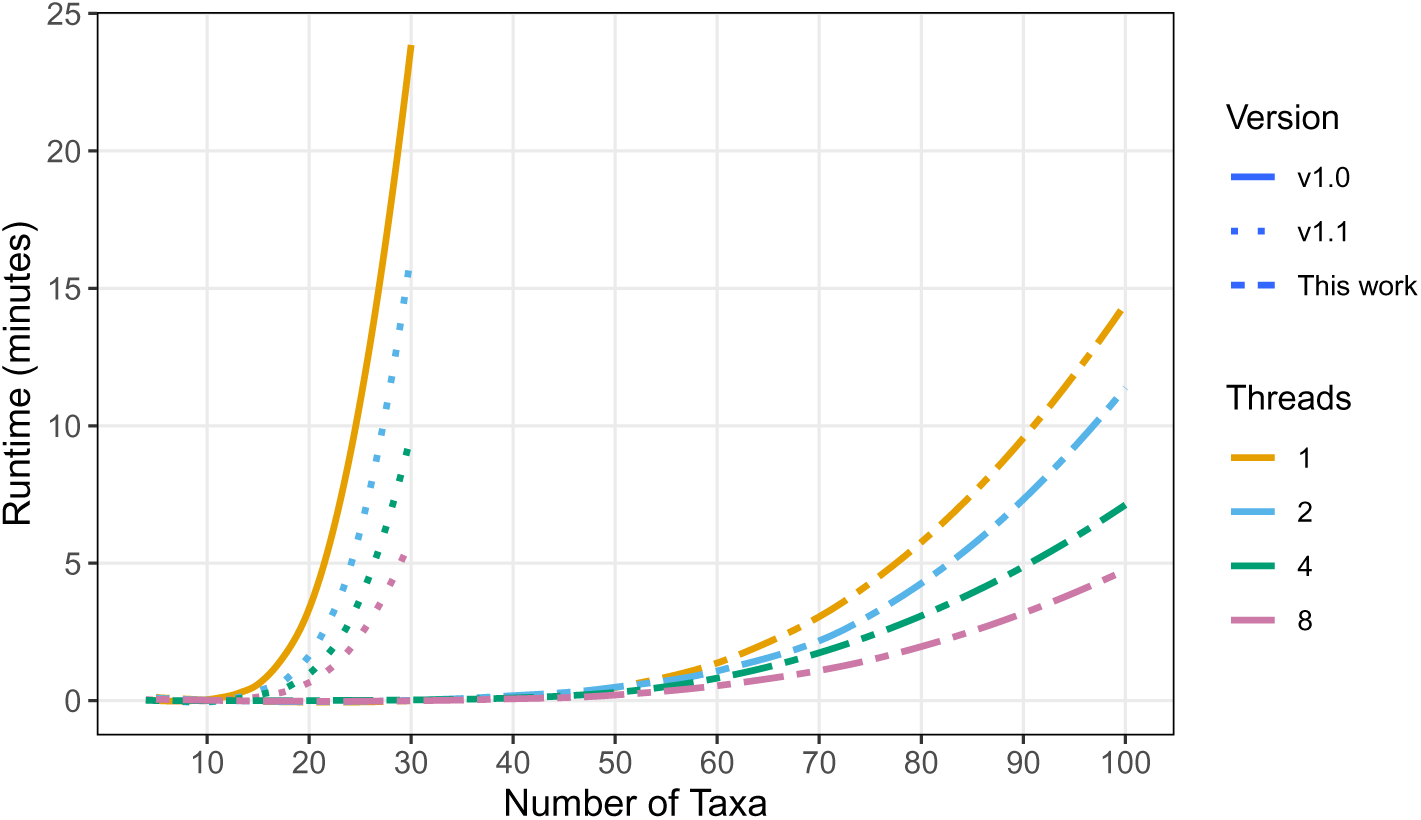
Time (in minutes) to optimize the parameters of a single network topology with varying numbers of taxa using various versions of SNaQ.jl (v1.0 [7], v1.1 [20]) by different number of threads. Results are averaged across randomly generated networks with 0-3 hybridizations.

### 3.5 Phylogeny of *Xiphophorus* (Poeciliidae)

Compared to the original L1-space analyses performed with SNaQ.jl v1.0, we found drastically better log-likelihood scores from the TCG-, TCGU-, and U-spaces when *h* ≥ 2 (Figure 6). All search spaces found the same optimal tree when *h* = 0, but networks inferred in C-space yielded worse NCLL scores than all other spaces except L1-space when *h >* 0. TCG-space was anomalously outperformed by L1-space when *h* = 1, but estimated networks with better NCLL scores when *h >* 1. When *h* = 1, L1- and TCG-space found the same network but with the direction of the reticulation reversed (Figures S17 and S23), explaining why the TCG-space network had a worse NCLL. C-space and U-space both inferred the same optimal network with *h* = 1.

**Fig. 6:**
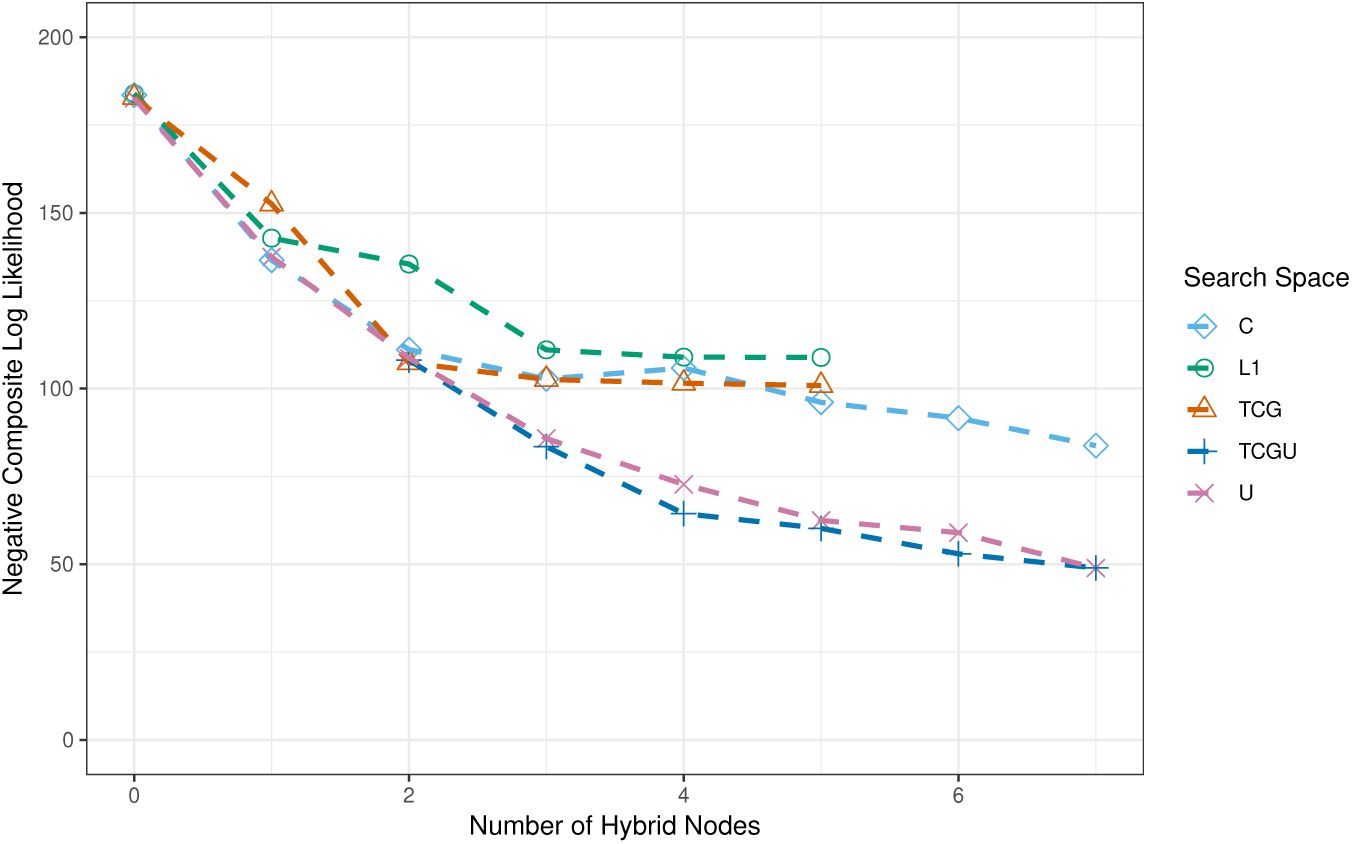
Negative composite log-likelihood (NCLL) scores of the best-fitting *Xiphophorus* species networks from each search space with each number of hybrid nodes *h*. Results from L1-space were taken from Solís-Lemus and Ané [7]. Inferences in TCG-space with *h_max_* = 6, 7 were conducted, but the best-fitting networks in these cases only ever contained at most *h* = 5 hybrid nodes.

The TCG-, TCGU-, and U-space networks with *h* = 2 were identical (Figures S24, S28, and S36, respectively), but when *h >* 2, L1-space was drastically outperformed by TCG-space, as was TCG-space by U- and TCGU-space. TCGU-space yielded marginally better results at each value of *h* than U-space (Figure 6), but it is unclear whether this is the result of a superior search strategy or because TCGU-space searches were effectively given more search iterations.

Despite performing inference in TCG-space with *h_max_* = 6, 7, the best-fitting networks in these cases all had only 5 hybrid nodes. This is due to the addition of a sixth hybrid node not improving the NCLL score enough to surpass the default optimization tolerances.

Best-fitting inferred networks in each search space are shown in Figures S16-S72, and model selection curves are given in Figure S15. The levels of the best-fitting networks for each space are given in Table 1. When analyzing all results at once, the best-fitting model is the network with 4 hybrid nodes from TCGU-space (Figure 7).

**Fig. 7:**
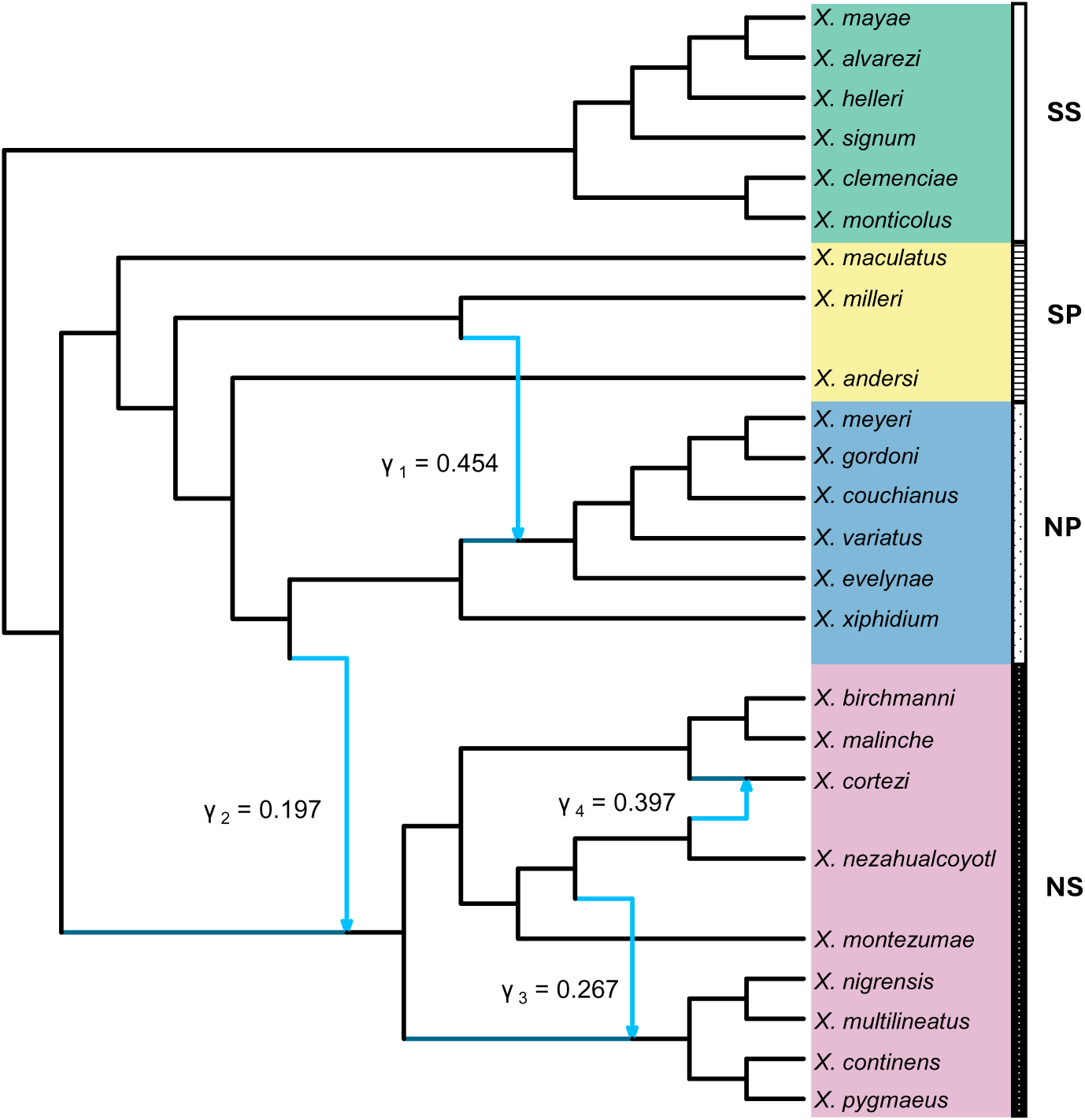
Best-fitting phylogeny of genus *Xiphophorus* inferred by SNaQ.jl after model selection, rooted with the southern swordtails (SS) outgroup clade. SP: southern platyfishes, NS: northern swordtails, NP: northern platyfishes. Black edges: internal tree edges (inheritance parameter *γ* = 1). Blue edges: internal hybrid edges with *γ̸*≠ 1. Inheritance parameters for minor edges (with *γ <* 0.5) in each pair are given.

**Table 1:** Level of the best-fitting networks for the *Xiphophorus* data from each search space with each number of hybrid nodes. Networks from L1-space are excluded because they are always level-0 when *h* = 0 and level-1 otherwise. *h* = 0 and *h* = 1 are excluded because they are always level-0 and level-1, respectively. **Bold, underlined**: the model selected by CAPUSHE [38] for each search space.

| Search Space | $h = 2$ | $h = 3$ | $h = 4$ | $h = 5$ | $h = 6$ | $h = 7$ |
| --- | --- | --- | --- | --- | --- | --- |
| TCG | <u>2</u> | 3 | 4 | 5 | N/A | N/A |
| TCGU | 2 | 2 | <u>2</u> | 3 | 3 | 6 |
| U | 2 | <u>2</u> | 2 | 5 | 5 | 6 |
| C | <u>2</u> | 1 | 1 | 1 | N/A | N/A |

Interestingly, networks selected by DDSE from each search space were all tree-child and galled, even in the TCGU- and U-spaces. Additionally, the best networks according to NCLL score from U-space with *h* ≤ 3 and TCGU-space with *h* ≤ 4 were tree-child and galled. Networks inferred in TCG-space were all level-*h* networks, where the level of the network was equal to the number of hybrid nodes in the network. For values of *h >* 5, no networks in TCGU- or U-space were tree-child and galled.

Across all search spaces, the network selected by DDSE was the network with *h* = 4 from TCGU-space (Figure 7), which we denote as the “best-fitting network”. This network contains two hybridizations in the NS clade, one hybridization in the NP clade and one hybridization from NS to NP. The SP clade is polyphyletic both in the major tree reported by Solís-Lemus and Ané [7] and in the major tree of the best-fitting network (Figure 7). In Solís-Lemus and Ané [7], the three clades NS, NP, and SS were each recovered as monophyletic. In contrast, our best-fitting network estimated the NP clade as polyphyletic, though only by a single taxon (*X. xiphidium*). Notably, if the inheritance probability *γ*_1_ were greater than 0.5 (here *γ̂*_1_ = 0.454), the NP clade would also be monophyletic in the best-fitting network, consistent with the topology reported in the original study.

Further, the reticulation corresponding to *γ*_2_ in the best-fitting network was the same as the level-1 network presented in Figure 10 of Solís-Lemus and Ané [7]. The reticulation corresponding to *γ*_3_ was contained in the level-1 network as well but with its direction reversed (with *X. nezahuacoyotl* as the hybrid descendant). Manually reversing this reticulation to match the direction of Solís-Lemus and Ané [7] in the best-fitting network and re-optimizing the network’s parameters yielded a worse NCLL score, suggesting that the new direction presented here better fits the data.

We build on the results of C-space in Appendix C.1 by removing the restriction of C-space to only tree-child and galled networks (denoted CU-space). Results from CU-space were highly comparable with results from U-space. Additionally, we experimented with the optimization parameters of SNaQ in CU-space and, in doing so, further reduced the cumulative runtime of network inference 13-fold while actually *improving* the NCLL score of the inferred networks. Further details are provided in Appendix C.1 and results are shown in Figure S14.

## 4 Discussion

A clear takeaway from our simulation study is that phylogenetic network analyses need to be interpreted very carefully, particularly those without a surplus of data. In 88% of TCG-space simulations with 100 gene trees, networks that were incorrectly inferred (SDHWCD > 0) had a better NCLL score than the true network (93% of 10 taxa networks, 86% of 20 taxa networks, and 74% of 30 taxa networks). This indicates that with limited data, the best topological estimates may be very distinct from the true evolutionary history, even when model assumptions are met. The issue occurs in all simulations but becomes less severe with more gene trees. Most notably, estimated concordance factors are extremely accurate when using 10,000 gene trees as input (median absolute error of 0.9%), but this issue still persists (41% with 10 taxa, 40% with 20 taxa, and 22% with 30 taxa). These results suggest that SNaQ is successful in finding networks that fit the data very well, but that the quartet-based approach may lack the specificity necessary to distinguish between meaningfully different network topologies, even with ample input data.

Hybrid ancestries appear to be accurately captured in networks of all sizes when the true network belongs to TCG-space (Figure 4). This trend is largely independent of the SDHWCD between the inferred network and true network and is largely independent of the amount of input data used as well. This suggests that SNaQ tends to infer the correct reticulate relationships between taxa even when it incorrectly infers specific siblinghood or intraclade relationships.

However, when the true network lies outside of the space of inferred networks, *F*_1_ scores indicated a more limited ability to identify hybrid clades; for simulations of TCNG and NTCNG networks average F_1_ scores were 0.63 and 0.25, respectively. Notably, in these cases precision was constantly higher than recall (see Figure S9), indicating that the taxa placed as hybrids by SNaQ tend to have accurate ancestry, but that SNaQ does not always find every relevant hybrid descendant. This explains why our re-analyses of the phylogeny of *Xiphophorus* (Poeciliidae) in TCGU- and U-space led to drastically better fitting networks with more hybridizations than TCG-space networks.

In general, edge length and *γ* estimates both improved as the level of ILS decreased and the amount of input data increased, as expected. Additionally, estimates tended to improve as the number of taxa in the network increased due to the fact that more quartet data is inherently available as the number of taxa in the network increases. Figure 3 shows that median errors in edge length estimates are near-zero but that there is also a high degree of variance in these estimates, even though the correct network topology was identified. This comes both from the natural variance in qCF estimates induced by estimation error as well as from the model simplifications that a quartet-based approach takes. These results indicate that the general magnitude of edge lengths tends to be accurately captured by SNaQ, but that researchers should exercise caution when performing analyses that require precise edge length estimates. With the exception of when the true topology was outside the topological search space, the median absolute error in *γ* parameter estimates was near-zero, even if the network topology was not correctly inferred; though in these cases, there was high variance in the absolute error of *γ* parameter estimates. This suggests that while most *γ* parameters are inferred very accurately on average, care should be taken when drawing conclusions from the value of any specific *γ* parameter.

In the case of the outlying networks with very poor parameter estimates, even with an SDHWCD of 0, we see a clear connection to anomalous networks. Anomalous networks are phylogenetic networks where an unrooted gene tree topology that *is not* displayed in the network has a higher probability of occurring than an unrooted gene tree topology that *is* displayed in the network. Ané et al. [16] conjecture that all anomalous networks must contain a hybridization with two descendant taxa (a so-called 3_2_-cycle), and prove the result for a restricted set of quarnets. This is exactly the topological feature that we noticed in 9 of the 11 outlying network topologies. A quarnet may contain a 3_2_-cycle and not be anomalous, however, depending on its numerical parameters. The prevalence of anomalous quarnets in these outlying networks ranged from 0% to 10%, indicating that both anomalous quarnets and non-amomalous 3_2_-cycles may pose practical difficulties when it comes to numerical parameter estimation under the MSNC model. In particular, despite these poor parameter estimates, SNaQ correctly inferred the topologies of these outlying cases with the same frequency as non-outlying replicates, and the NCLL scores of the inferred networks in these outlying cases were either better than or very close to the NCLL scores of the respective true networks.

The phylogeny of *Xiphophorus* fishes is proposed to include both ancient and contemporary hybridization events [21, 23, 24, 39]. One such event is the hypothesized ancient hybridization between the southern platyfish (SP) and southern swordtail (SS) clades followed by repeated backcrossing in hybrid females. As a result, mitochondrial DNA places *X. clemenciae* within the SP clade, whereas nuclear DNA instead places it with the SS clade [21, 23, 39, 40]. Our analyses are based on gene trees constructed from nuclear DNA, which is why *X. clemenciae* is placed in SS without any signal of hybridization. Recent work has already utilized mtDNA to infer the hybrid ancestry of *X. clemenciae* [21], but future work could seek to model this event in the context of the broader phylogeny of *Xiphophorus* fishes.

Our best-fitting network (Figure 7) recovers hybrid ancestry that is broadly consistent with previous studies of the genus, while also resolving additional events not captured by earlier phylogenies. The hybrid event ancestral to several members of the NP clade, with parents *X. milleri* and *X. xiphidium* (*γ*_1_), matches the findings of Du et al. [21]. Two further events are consistent with Rosenthal et al. [23]: the mating of SP platyfish females with an ancestral swordtail lineage (*γ*_2_), and contemporary hybridization between *X. malinche* and *X. birchmanni* (*γ*_4_). The latter hybridization (*γ*_4_) additionally corroborates evidence that *X. nezahualcoyotll*’s transcriptome was largely derived from that of *X. cortezi* [24]. More broadly, our network also reflects the finding that *X. nezahualcoyotll* exhibits evidence of having ancient hybrid ancestry, but not contemporary hybrid ancestry [24]. Beyond these, we recover a novel hybrid event ancestral to a small portion of the NS clade (*γ*_3_ = 0.267) not previously documented in the literature.

It should be noted that more recent work has refuted the placement of *X. continens* as sister to *X. pygmaeus*, instead placing it sister to *X. montezumae*. This appears to be due to misidentification of materials in previous studies, including the data from which the gene trees used in these analyses were inferred [21, 41].

Expanding the space of inferable networks from level-1 to include tree-child and galled networks and further to all semi-directed phylogenetic networks led to clear, drastic improvements in the NCLL scores of the networks inferred on these species. Interestingly, every network in each space chosen by model selection was level-2. This may suggest that the true underlying model (or a close approximation of it) is level-2. Our highly restricted C-space analyses showed promise by yielding comparable or better networks than TCG-space for each value of h. Further, search through C-space yielded network topologies with 6 and 7 hybrid nodes whereas the search through TCG-space could not, even when *h_max_* > 5. We additionally conducted a similar search where C-space was not bound to tree-child and galled networks (denoted CU-space) and saw results that were comparable to the results of U-space (Figure S14). This indicates that restricting the network search space with topological information known a priori may be highly beneficial in reducing inference runtimes by filtering out obviously incorrect network topologies from the search.

On the other hand, we showed that searching only through identifiable network space is harmful to network inference. The best-fitting network overall belonged to TCG-space and C-space but was only able to be inferred by allowing the network search to traverse through the space of non-identifiable networks. Most computationally practical, model-based approaches up to this point have restricted practitioners and systematists to network spaces that are known to be statistically identifiable, even if it does not contain the true evolutionary history. While topological identifiability of various classes of networks is an area of active research [15, 42–44], waiting on these results holds back the progress of inference methods while simultaneously hindering the ability of researchers to infer evolutionarily accurate phylogenies today.

## Data Availability

The SNaQ method is available as open-source software written in Julia, hosted on GitHub at github.com/JuliaPhylo/SNaQ.jl. Simulation and empirical data have not yet been digitally archived but will be prior to publication.

## Acknowledgements

This work was supported by the National Science Foundation (DEB-2144367 to CSL). The authors thank Cécile Ané for discussions about network space.

## A Composite Likelihood and Restricted Search with SNaQ.jl

A small detail worth noting is that SNaQ technically optimizes and reports the value of the function given in 1 plus the constant 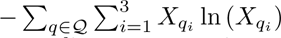. ln (X*_qi_*). This adjustment does not impact the optimization process, gradient computations, or other technical details, but does improve the interpretability of results because, with this adjustment, ℓ(*N* |*X*) = 0 when all *X_qi_* = *E_N_* (*X_qi_*). Thus, the optimal NCLL score with this adjustment is 0, whereas without this adjustment it is some arbitrary constant that will vary depending on the sample data.

The algorithm of Ané et al. [16] is defined on 4-taxon networks, but can be applied to *n*-taxon networks by, for each quartet *q* ∈ *Q*, copying the network, removing all taxa not in *q* from the copy, and running the algorithm on the resulting 4-taxon network.

**Table S1:** The structure of all possible values in an expected quartet concordance factor (qCF) computation graph and their corresponding derivatives. Note that every leaf in a graph is a vector with length 3 comprised of some combination of only the last four rows in this table.

| Expected qCF Node Value | $d/dt_i$ | $d/d\gamma_i$ |
| --- | --- | --- |
| $\gamma_i$ | 0 | 1 |
| $1 - \gamma_i$ | 0 | -1 |
| $\gamma_i(1 - \gamma_i)$ | 0 | $1 - 2\gamma_i$ |
| $\gamma_i^2$ | 0 | $2\gamma_i$ |
| $(1 - \gamma_i)^2$ | 0 | $-2(1 - \gamma_i)$ |
| $\exp\{-\sum_{e \in E} t_e\}$ | 0 if $i \notin E$, else $-\exp\{-\sum_{e \in E} t_e\}$ | 0 |
| $1 - \exp\{-\sum_{e \in E} t_e\}$ | 0 if $i \notin E$, else $\exp\{-\sum_{e \in E} t_e\}$ | 0 |
| $1/3 \exp\{-\sum_{e \in E} t_e\}$ | 0 if $i \notin E$, else $-1/3 \exp\{-\sum_{e \in E} t_e\}$ | 0 |
| $1 - 2/3 \exp\{-\sum_{e \in E} t_e\}$ | 0 if $i \notin E$, else $2/3 \exp\{-\sum_{e \in E} t_e\}$ | 0 |
| 1 | 0 | 0 |
| 0 | 0 | 0 |

Previous versions of SNaQ.jl were hard-coded to only consider networks that were level-1 and whose topologies were identifiable. Here, we extend SNaQ.jl to allow for user-specified search spaces with the default being set to the space of networks that are both tree-child and galled. To decide whether a network belongs to the user-specified space, SNaQ.jl checks its proposed network against the user-specified search function at each iteration. If the proposed network belongs to the search space then the algorithm continues as normal. If not, the proposal is immediately rejected, and the rejection does not increment the total number of sequentially failed iterations, and so does not interact with the parameter Nfail.

Well-implemented functions to check for restrictions in a network’s topology are many orders of magnitude faster than likelihood optimization functions, so such an approach does not lead to any notable increase in method runtime. For example, benchmarking on a network with 20 taxa and 7 hybrids, optimizing the network’s parameters took on average 10^9^ times as long as checking if the network was both tree-child and galled.

**Fig. S1:**
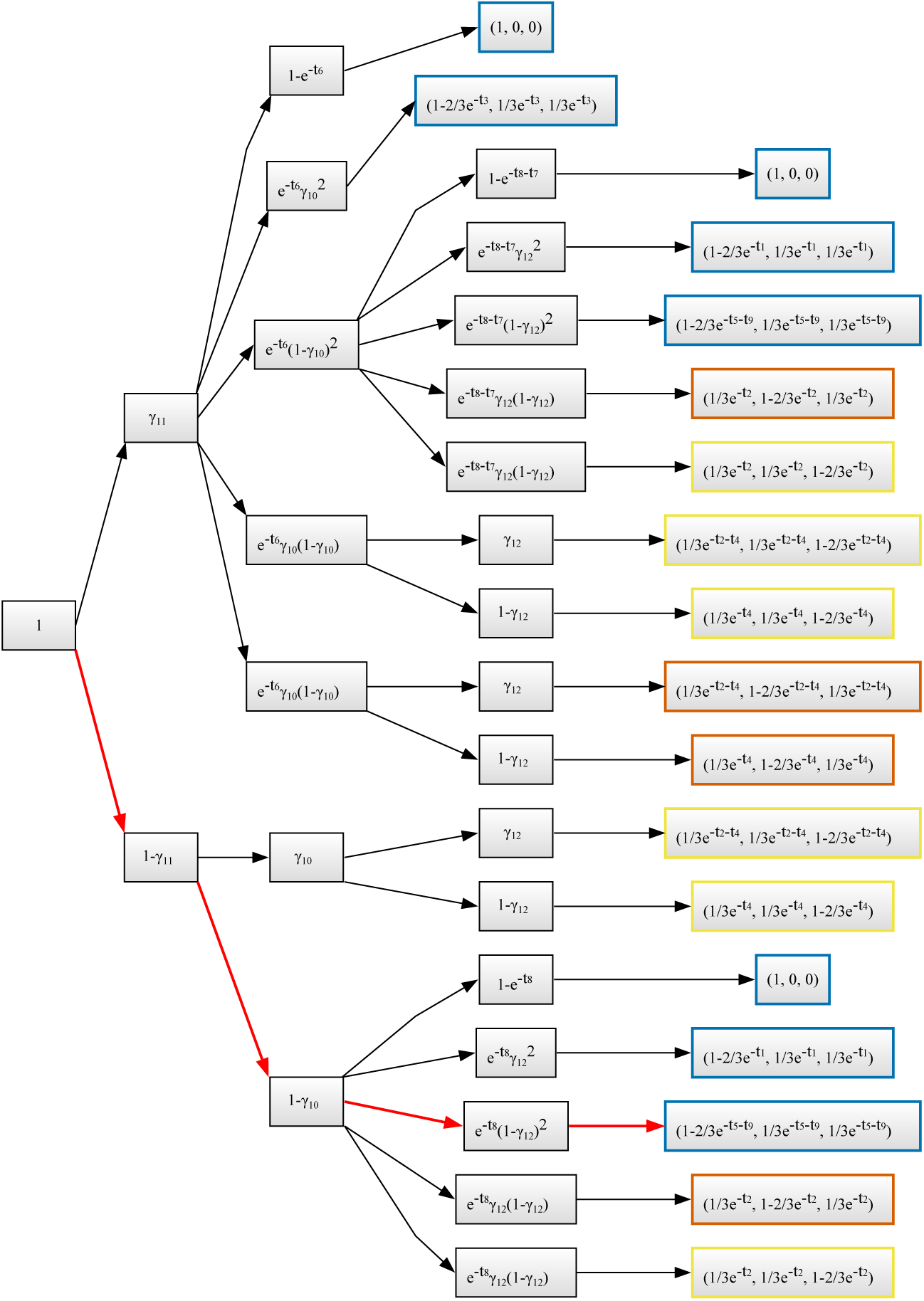
Example of the data structure used to store expected quartet concordance factor (qCF) equation information. The quartet that this graph was generated from is given in Figure S2. The leaf nodes in this graph (right-most nodes) are vectors each of three values. Each value corresponds to each of the three unrooted qCFs that can arise from a species network with four taxa, and more broadly to a parental tree of the quartet. **Red lines:** The specific coalescent history shown in the quarnet given in Figure S2. The corresponding coalescent histories in Figure S2 are halted as soon as the corresponding qCFs are determined. The probability of observing each possible unrooted quartet in this coalescent history is calculated by multiplying together each of the values of the nodes in the red highlighted path.

**Fig. S2:**
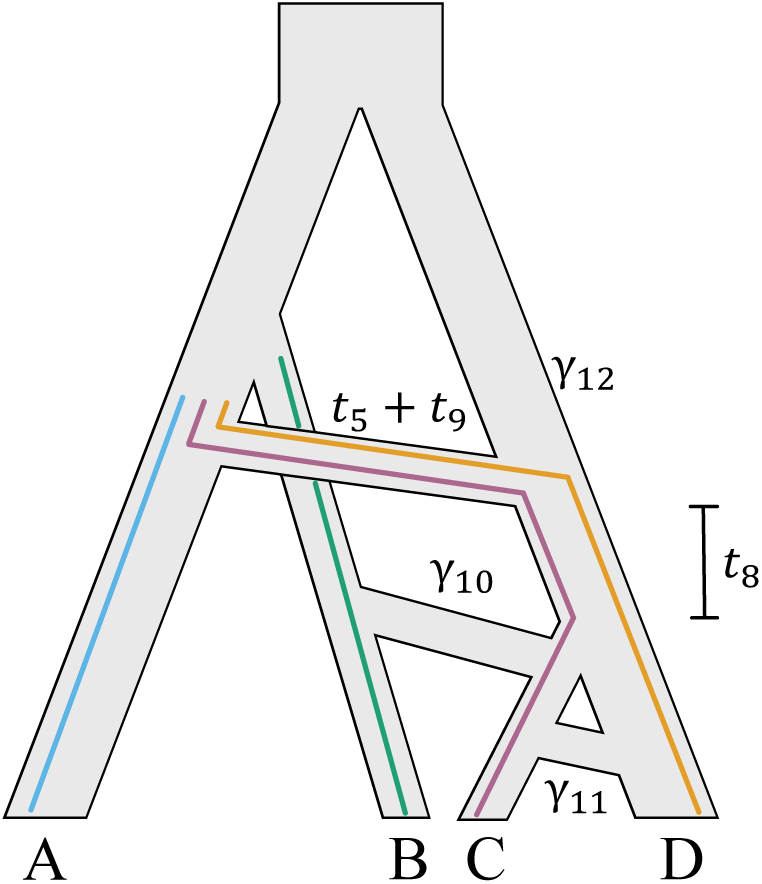
Species quarnet whose expected quartet concordance factor (qCF) computation graph is given in Figure S1. The probability of the specific coalescent history shown here can be computed by traversing the red lines in Figure S1 and taking the product of every node that is passed through. Probabilities of other coalescent histories can be calculated similarly. Minor hybrid edges are labelled with their corresponding inheritance proportions (*γ*). One edge has length *t*_5_ +*t*_9_ because this edge in the quarnet spans two edges in the larger network that this quarnet was extracted from.

Several other useful and common search spaces are written into SNaQ.jl, including restrictions for level-k networks and unrestricted search space. These spaces can be utilized with the following syntax, respectively:

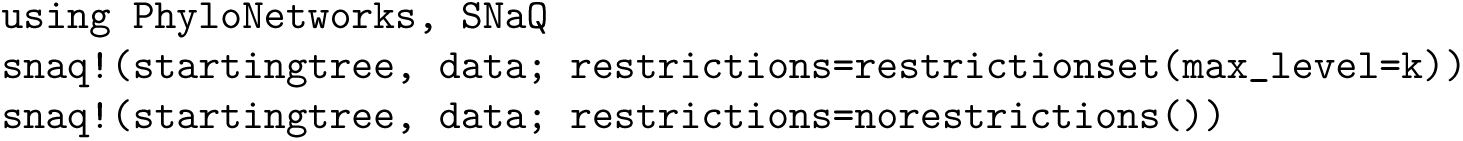

Additionally, custom restrictions can be written that leverage specific topological features of the network. The sole requirement of a custom restriction function is that it takes a HybridNetwork object from the PhyloNetworks.jl package [45] as input and that it returns a bool. Below is an example of a restriction that requires hybrid nodes to have at least 2 children. Such a restriction could be useful if all taxa in the network were sampled at least twice.

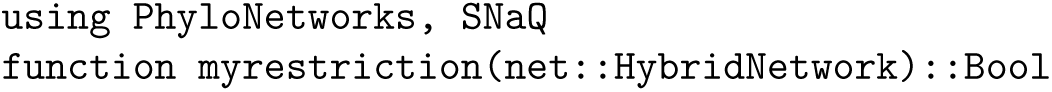
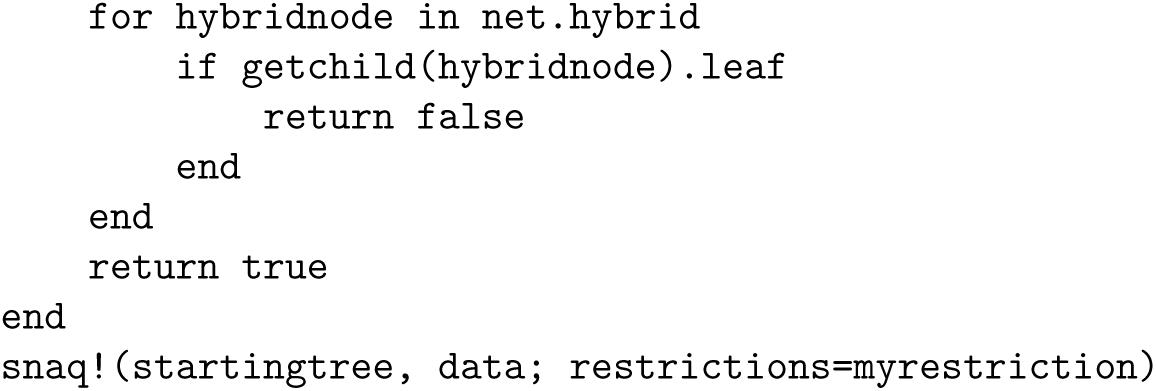

## B Simulation Study

### B.1 Simulated Data Generation

To generate ground truth networks, we used the SiPhyNetwork package in R [31] to simulate networks under a birth-death-hybridization model. These networks were simulated with 10, 20, and 30 taxa, low and high average densities of reticulations, and from the space of tree-child and galled networks (TCG), the space of tree-child but not galled networks (TCNG), and the space of neither tree-child nor galled networks (NTCNG). To obtain networks with varying average densities of reticulations, we varied the speciation rate (*λ*) and hybridization rate (*ν*) parameters in SiPhyNetwork as given in Table S2. The level of ILS in each network was varied by scaling the length of each edge by a constant such that the network’s average edge length was equal to a fixed value. The average edge length in the network was fixed at 2.0 coalescent units (cu) for low ILS simulations and 0.5 cu for high ILS simulations.

**Table S2:** Speciation rate (*λ*) and hybridization rate (*ν*) parameters for simulating ground truth networks with the SiPhyNetwork R package.

| Number of Taxa | Reticulation Density | $\lambda$ | $\nu$ |
| --- | --- | --- | --- |
| 10 | Low | 0.5 | 0.03 |
| 20 | Low | 0.5 | 0.02 |
| 20 | High | 0.6 | 0.05 |
| 30 | Low | 0.5 | 0.0125 |
| 30 | High | 0.6 | 0.03 |

Networks with 10 taxa were only simulated with low reticulation density because SiPhyNetwork almost never produced networks that were TCG with high reticulation density under any combination of *λ* and *µ*. The distribution of reticulations in networks belonging to each set of simulation parameters is given in Figure S3. All inheritance parameters *γ* were drawn from a Beta(10, 10) distribution. For each of the 5 combinations of number of taxa and reticulation density, 25 unique networks were simulated.

SiPhyNetwork does not provide any functionality for restricting the space of the generated network. In order to restrict the networks to each of the respective spaces, we simulated arbitrary networks and only kept networks that belonged to each corresponding space. This was repeated for a given space until 25 networks were obtained.

Ground truth gene trees were simulated from ground truth networks with the PhyloCoalSimulations Julia package [30]. Simulations were conducted with 100, 1,000, and 10,000 gene trees (loci), each containing all of the network’s taxa.

These gene trees were used to simulate multiple sequence alignments (MSAs) with seq-gen [32] under the HKY85 model of evolution [33] with rate parameters 0.3, 0.2, 0.2, 0.3 and either 500 or 1,000 base pairs.

Then, gene trees were inferred from the simulated MSAs with IQ-TREE [34] without specifying the true underlying model of evolution. Finally, quartet concordance factors were calculated from this set of inferred gene trees and used as input to the snaq! function in SNaQ.jl. For each simulation, 20 independent inference runs were conducted, and each run continued until 2,000 consecutive networks were proposed that did not improve the composite log likelihood past the default tolerances (the Nfail parameter of snaq!). Further, the space of networks searched was restricted to the space of networks that are both tree-child and galled with identifiable topologies. This was done by setting the named argument restrictions in the snaq! function to the tcgidentifiable function provided in SNaQ.jl, as below. Additionally, the maximum number of hybridizations in the inferred network (specified by hmax) was set to the number of hybridizations in the ground truth network (*h* below). In most cases, the inferred network had exactly this many reticulations, but in cases with particularly high values of *h*, this was not always the case.

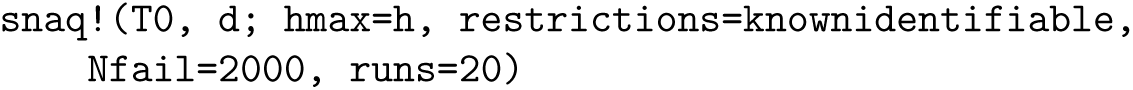

### B.2 Distribution of Number of Hybridizations

### B.3 TCG Outlier Networks

Below are the unique topologies of the networks that were identified as outliers with respect to parameter estimation (i.e., as shown in Figure 3).

**Fig. S3:**
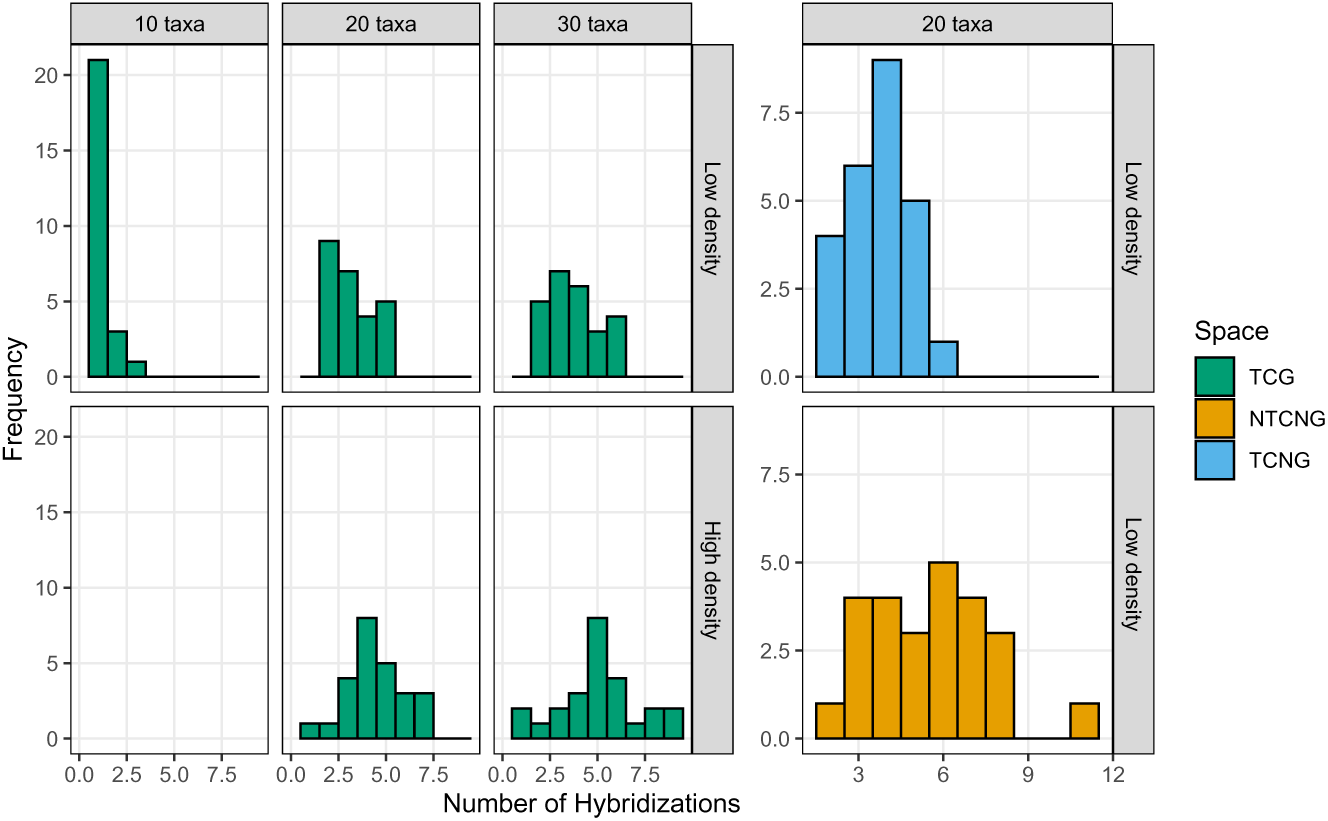
Distribution of the number of reticulations in each replicate number based on the number of taxa in the network, the reticulation density (“Low” and “High” density), and the network space.

**Fig. S4:**
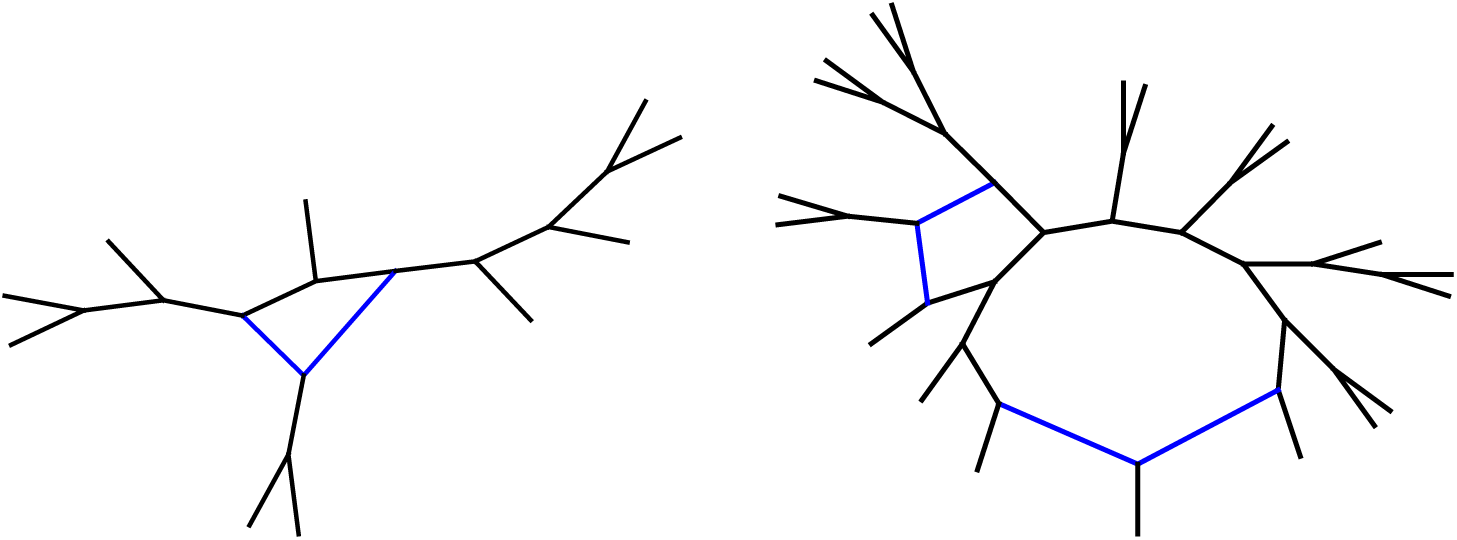
Two examples of the true (unlabeled) networks with outlying *γ* and *t* parameter estimates. Tree edges are drawn in black while hybrid edges are drawn in blue. **Left:** 10 taxa, 1 hybrid; **right:** 20 taxa, 2 hybrids. Note that both networks share the topological property of having at least one hybrid node with exactly two descendants.

**Fig. S5:**
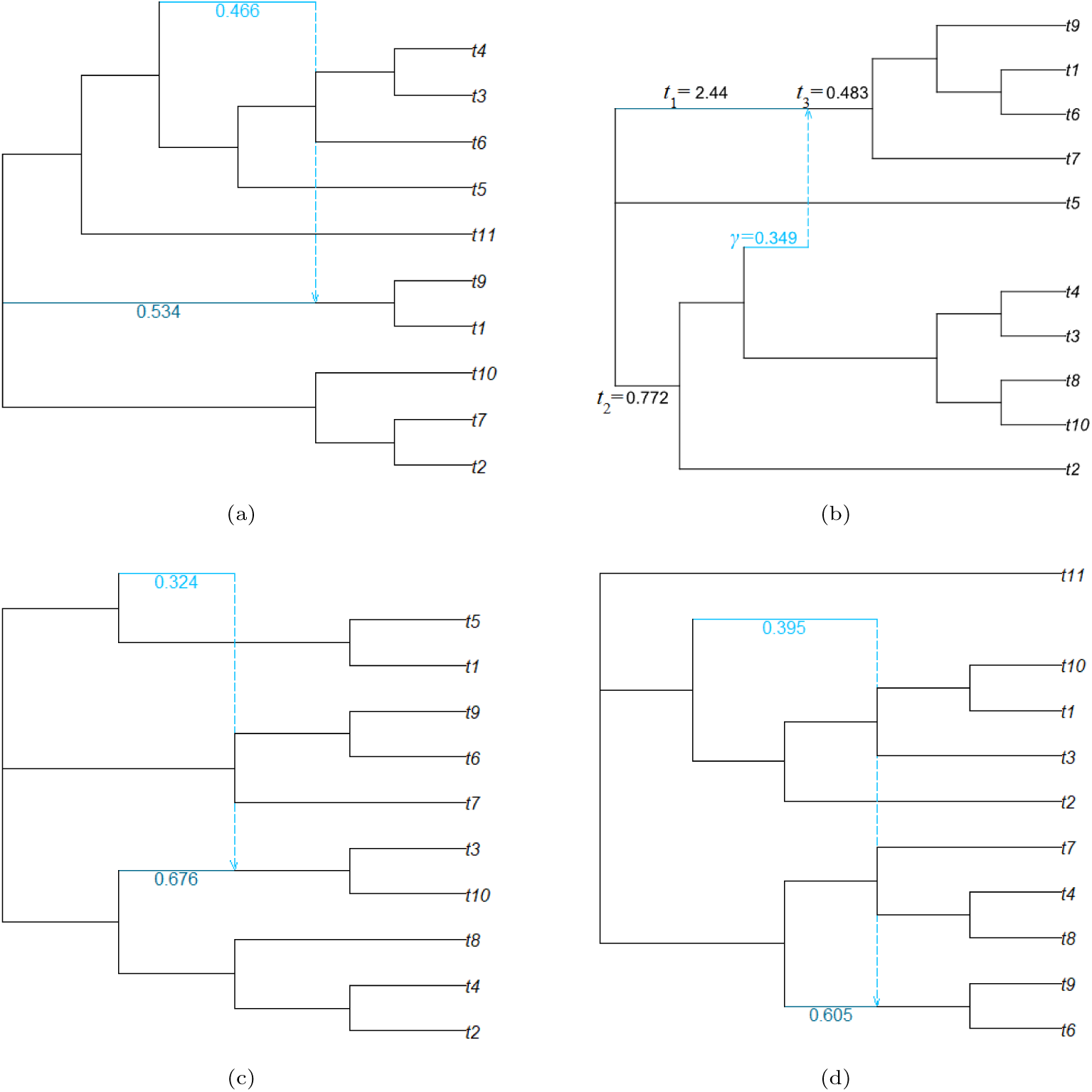
Outlier 10 taxa replicate networks. All networks are semi-directed with tree edges drawn in black and hybrid edges drawn in blue. Networks (a), (c), and (d) have their major and minor hybrid edges labelled with their respective *γ* values. Network (b) does not have the topological feature of one hybridization with two descendants but exhibits numerical instability around parameters *t*_1_*, t*_2_*, t*_3_, and *γ*.

**Fig. S6:**
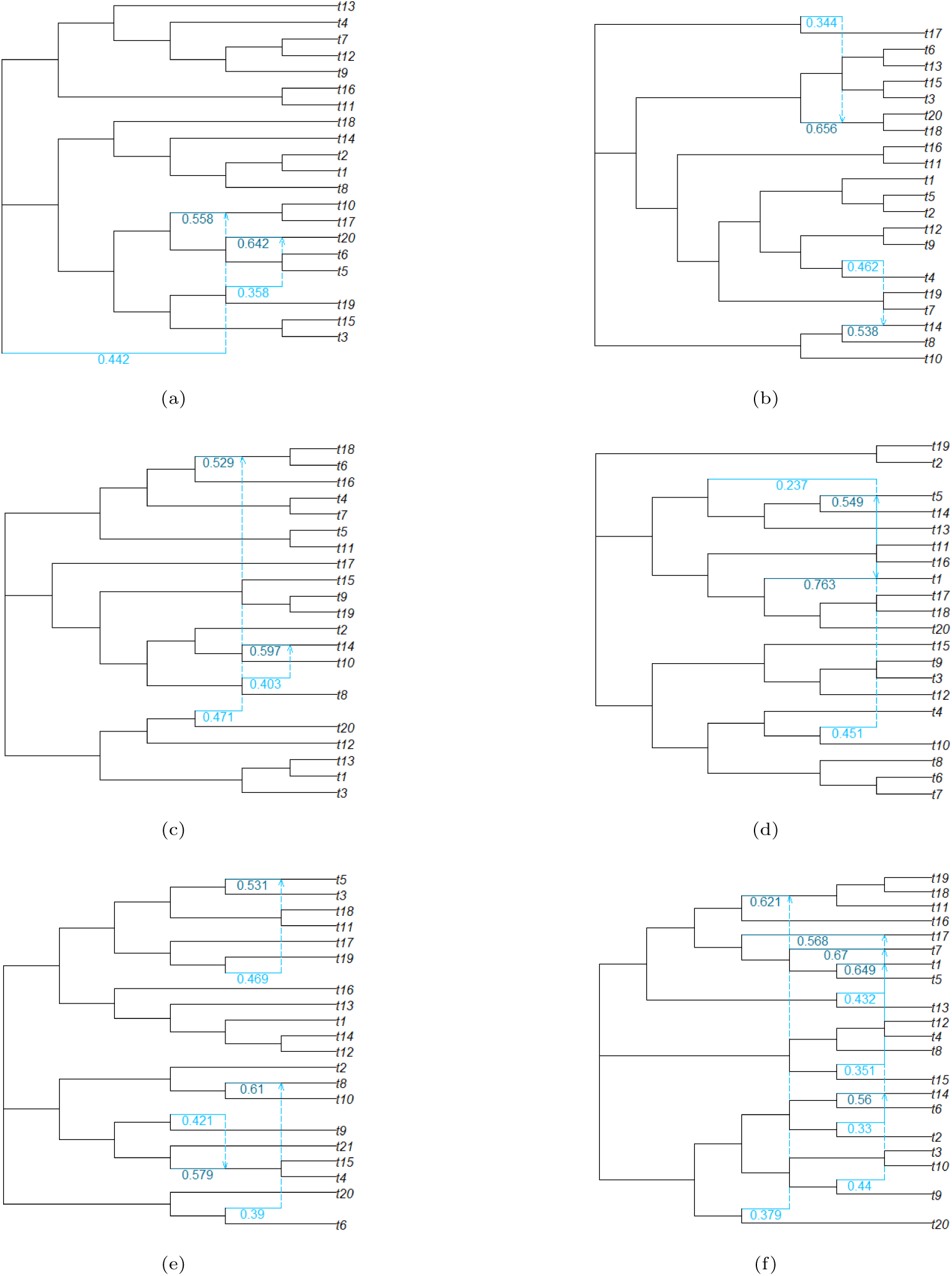
Outlier 20 taxa networks. All networks are semi-directed with tree edges drawn in black and hybrid edges drawn in blue. Major and minor hybrid edges are labelled with their associated inheritance probabilities. Networks (a)-(c), (e), and (f) exhibit the topological feature of a hybridization with two descendants discussed in the main text, whereas network (d) does not.

**Fig. S7:**
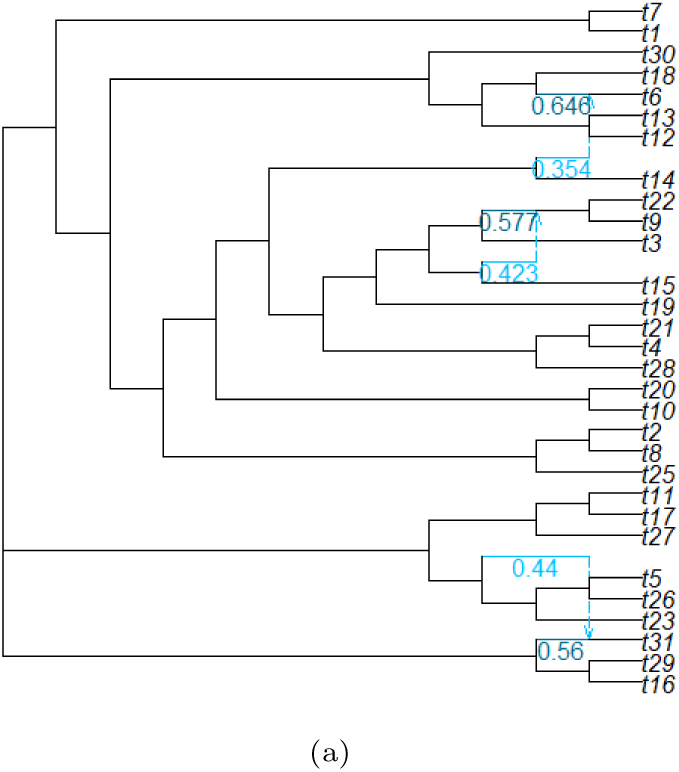
The only outlier 30 taxa network. Tree edges drawn in black and hybrid edges drawn in blue. Major and minor hybrid edges are labelled with their associated inheritance probabilities.

**Fig. S8:**
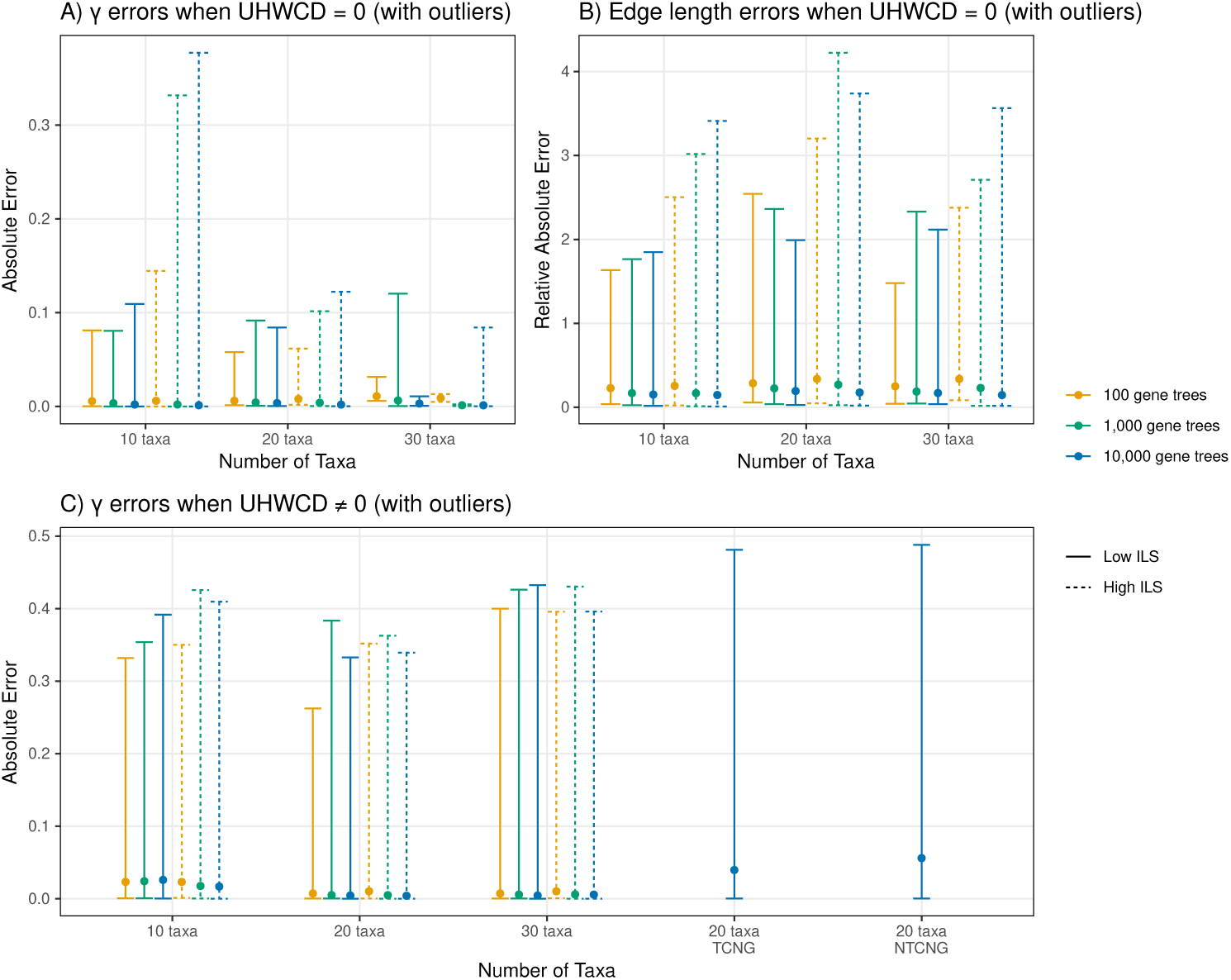
Distributions of observed parameter estimation errors across simulation parameters **including outliers**. Lines: 95% observed confidence regions (*q*_0.025_*, q*_0.975_) for inference errors. Points: median observed inference errors. **A)** Absolute error in inferred inheritance parameters (*γ*) when the semi-directed hardwired cluster dissimilarity (SDHWCD) between the true and inferred networks was 0. Four outlier networks were removed from each of the 10 taxa and 20 taxa simulations. **B)** Relative absolute error in edge length parameters when the SDHWCD between the true and inferred networks was 0. **C)** Absolute error in inferred inheritance parameters when the SDHWCD between the true and inferred networks was not 0.

### B.4 TCNG and NTCNG Simulations

**Fig. S9:**
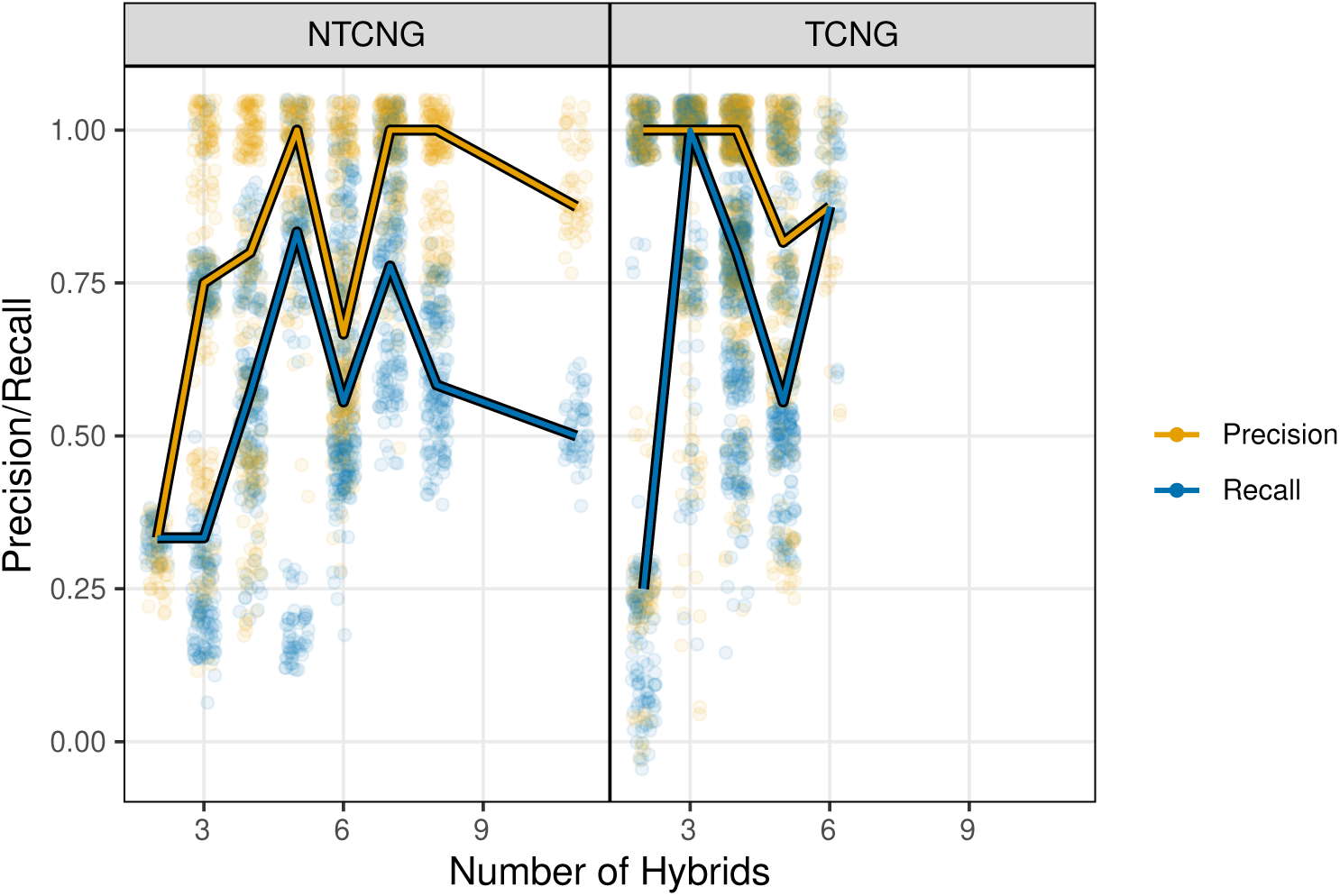
Distribution of the precision and recall values for simulations where the ground truth species network was TCNG (tree-child but not galled) and NTCNG (neither tree-child nor galled) with respect to placement of hybrid ancestry. Lines are median values for each metric. Points are all simulation values with small, random perturbations so that point density is readily apparent.

### B.5 Runtimes

Method runtimes increased polynomially with respect to the total number of taxa in the network and appear to increase roughly proportionally to the square root of the number of hybrids in the network (Figure S13). Median runtimes were comparable in all simulations, though TCNG and NTCNG simulations had marginally longer runtimes when compared to simulations with TCG networks.

**Fig. S10:**
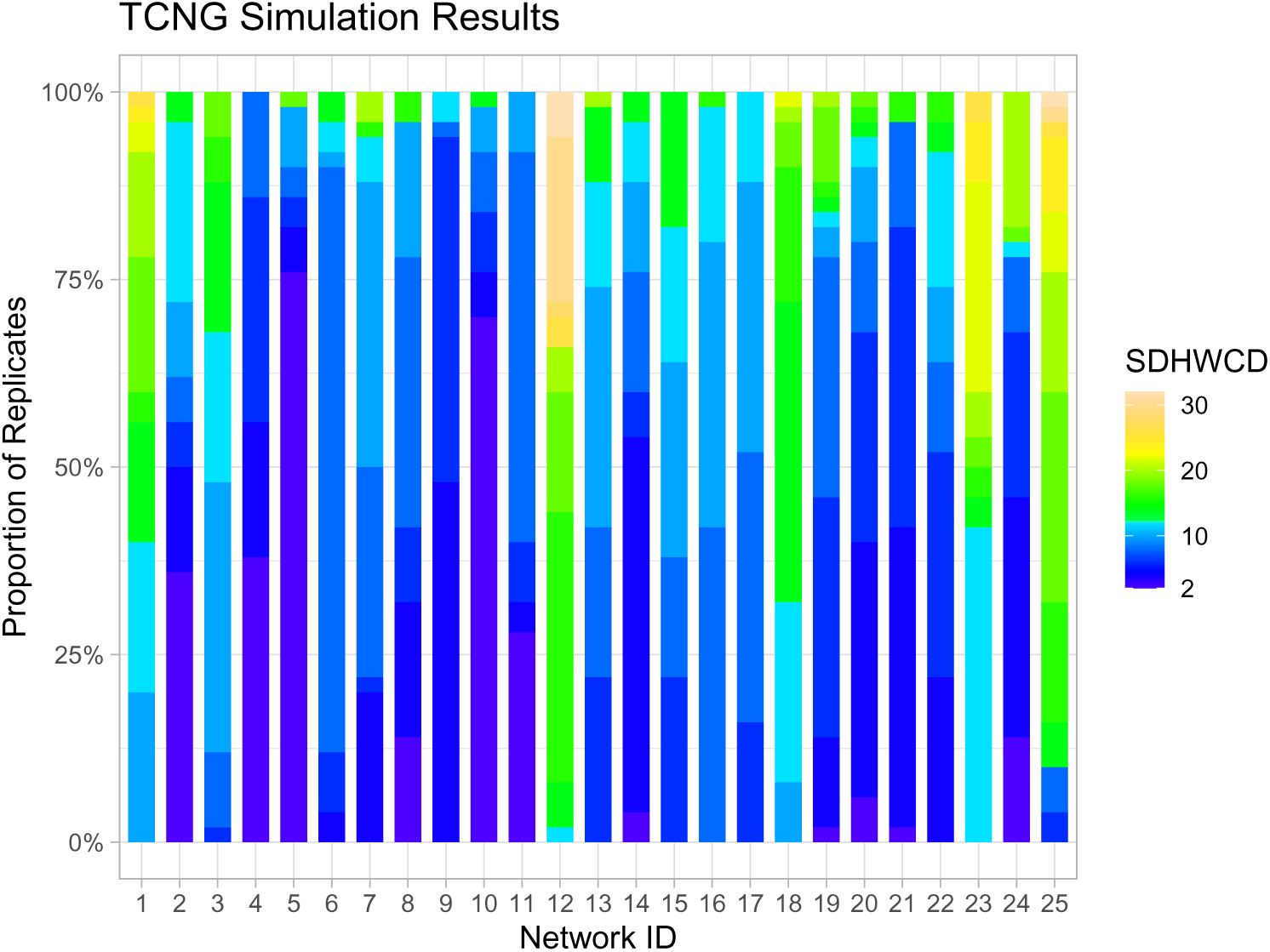
Distribution of semi-directed hardwired cluster dissimilarity (SDHWCD) results for networks inferred when the true network belonged to the class of tree-child and not galled (TCNG) networks. Network ID distinguishes the 25 distinct TCNG topologies that were simulated. The minimum SDHWCD shown is 2 because networks were inferred from TCG space while true networks were from TCNG space, so a SDHWCD of 0 is impossible.

**Fig. S11:**
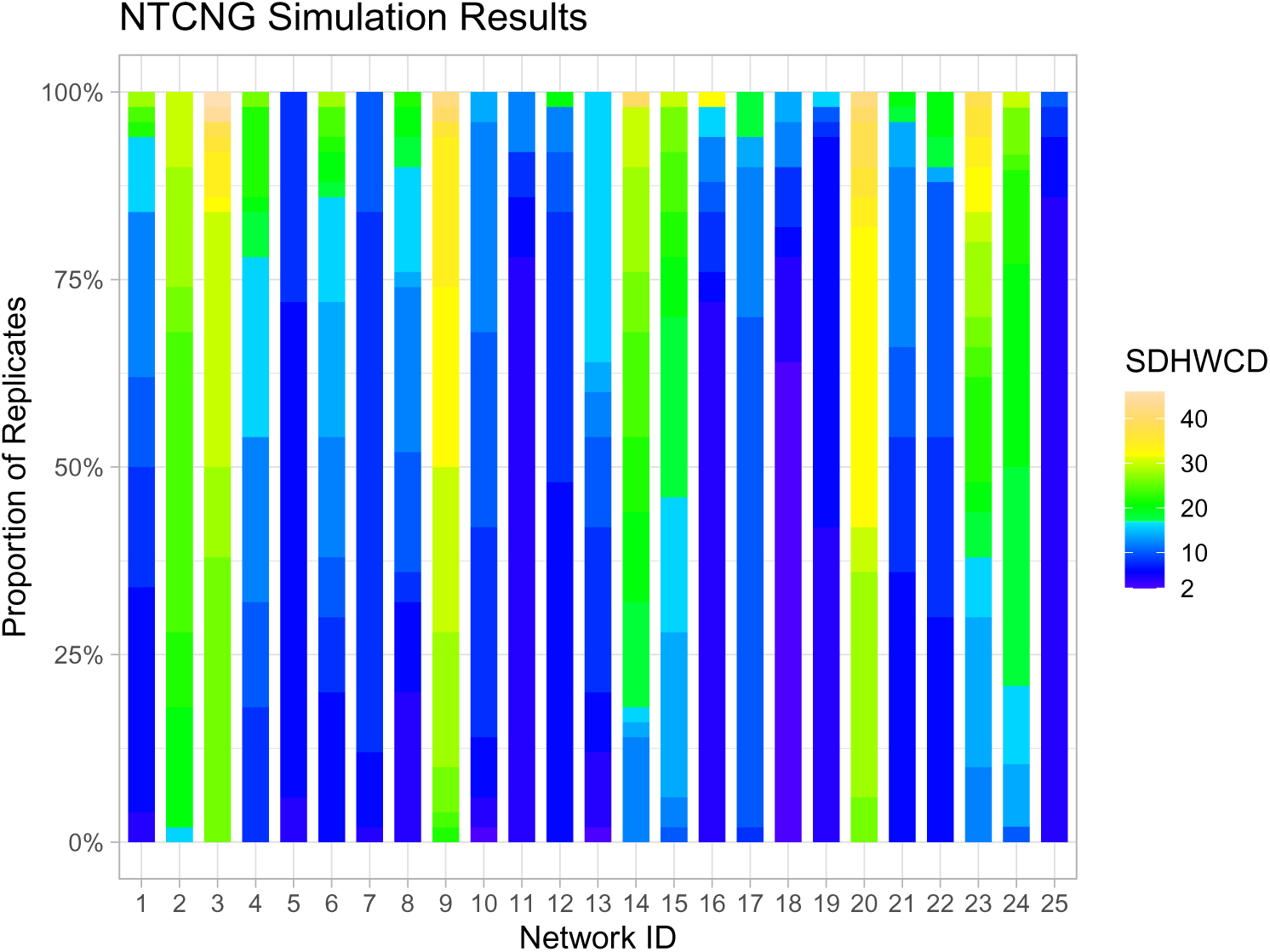
Distribution of semi-directed hardwired cluster dissimilarity (SDHWCD) results for networks inferred when the true network belonged to the class of not tree-child and not galled (NTCNG) networks. Network ID distinguishes the 25 distinct NTCNG topologies that were simulated. The minimum SDHWCD shown is 2 because networks were inferred from TCG space while true networks were from NTCNG space, so a SDHWCD of 0 is impossible.

**Fig. S12:**
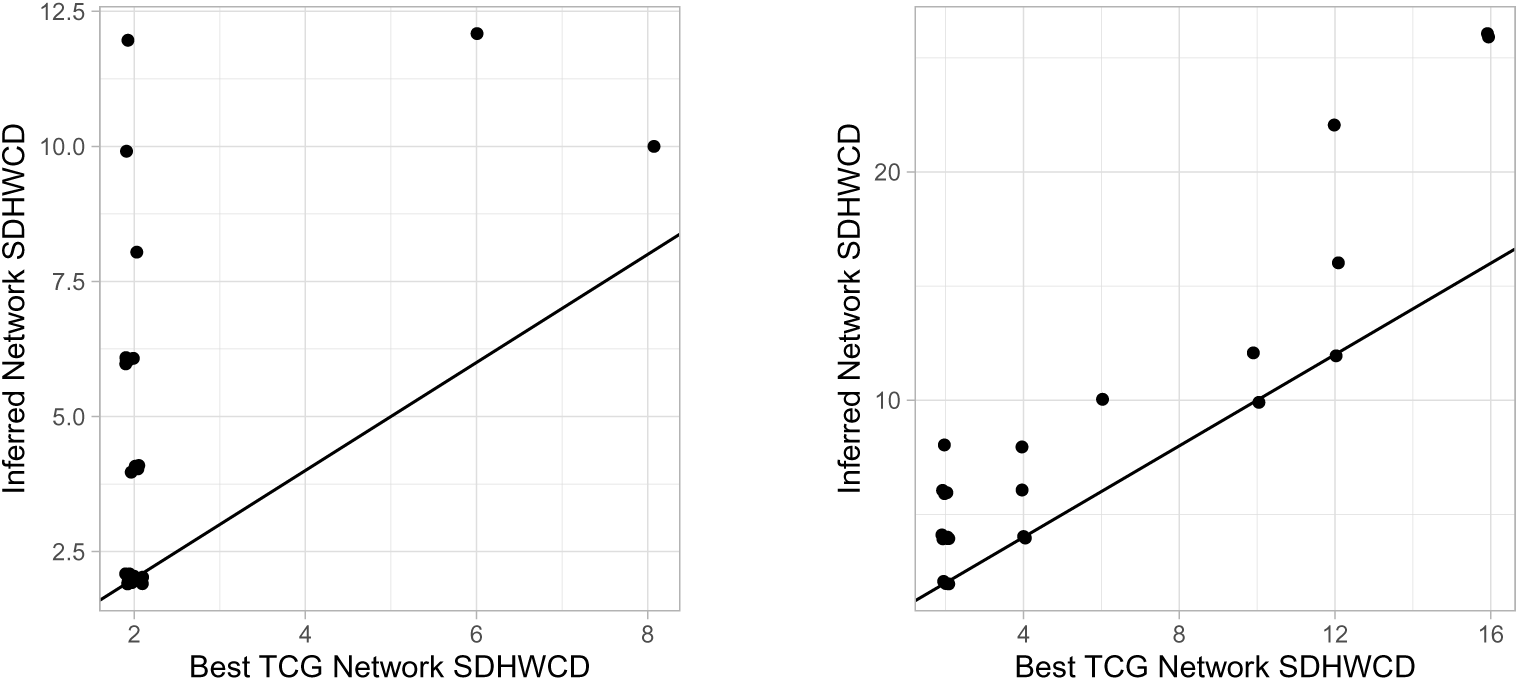
Semi-directed hardwired cluster dissimilarities (SDHWCD) of inferred networks where the true network was either tree-child and not galled (TCNG; left) or not tree-child and not galled (NTCNG; right) compared with the approximate best possible SDHWCD attainable with a tree-child and galled (TCG) network.

**Fig. S13:**
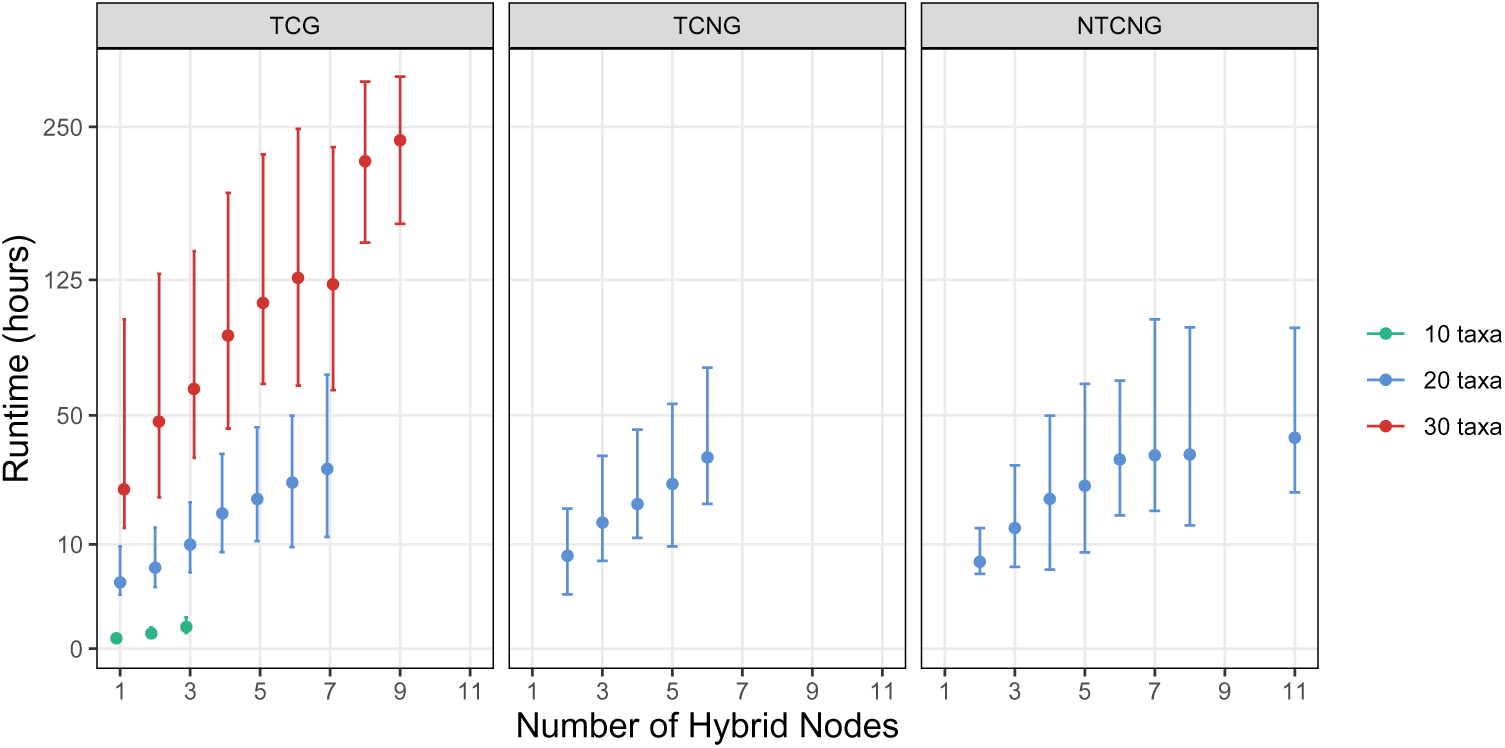
Distribution of wall-clock runtimes for SNaQ when inferring networks with ground truth data generated from **TCG** (tree-child and galled), **TCNG** (tree-child and not galled), and **NTCNG** (not tree-child and not galled) network spaces. All inferred networks belonged to TCG space. Lines: 95% observed confidence regions (q_0.025_, q_0.975_). Points: median runtimes.

## C Phylogeny of *Xiphophorus* (Poeciliidae)

### C.1 Additional Search Strategies

In the main text, we compared 6 different search spaces for inferring the phylogeny of *Xiphophorus* (Poeciliidae) from empirical data: L1-, TCG-, TCGU-, U-, C-, and CTCG-space. Following the positive results of inference in C-space, we chose to further examine the performance of SNaQ in C-space without the tree-child and galled restriction, and then with different strategies with respect to optimization. We examined the following spaces and strategies:

1. C-space but without the tree-child and galled restriction (CU-space).
2. CU-space with optimization parameters Nfail=50 and propQuartets=0.5 (denoted CUfast).
3. CU-space with optimization parameters Nfail=20 when *h* = 0, otherwise Nfail=40, propQuartets=0.05 for all *h*, and starting topologies selected according to Algorithm 1 for *h >* 0 (denoted CUfast500).

**Algorithm 1.**
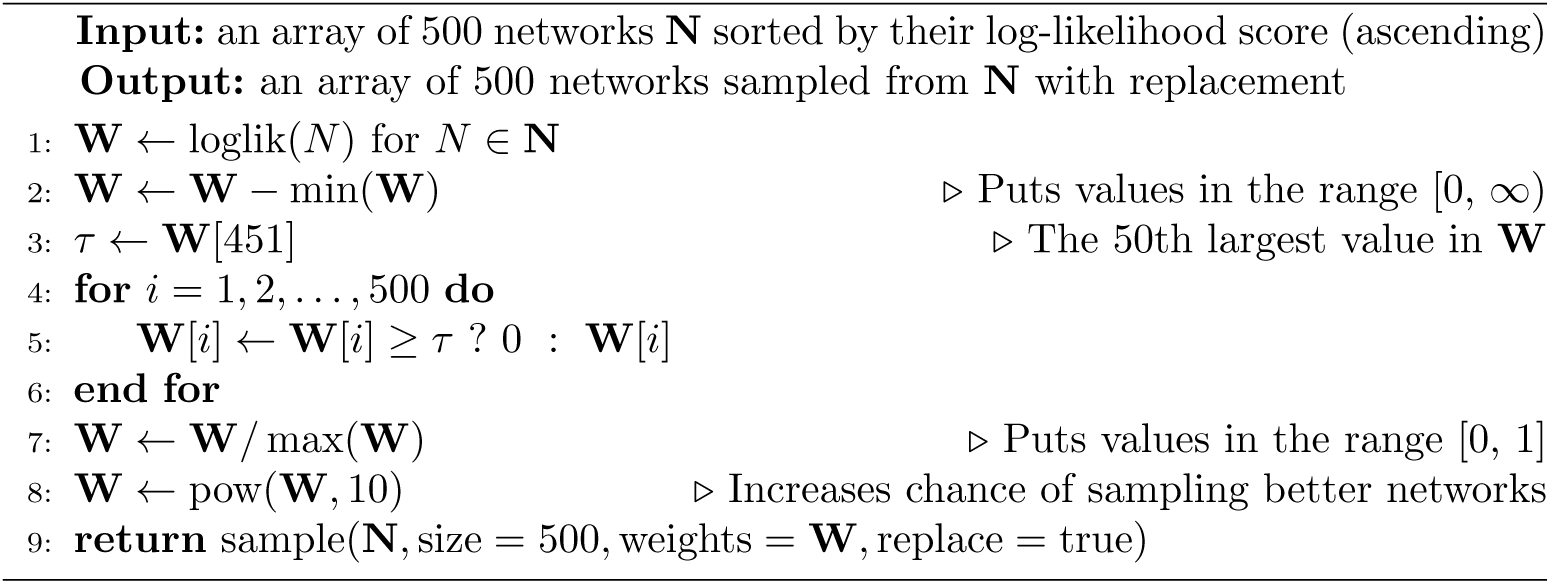
Weighted sampling strategy for the CUfast500 search strategy.

In short, Algorithm 1 picks starting networks for *h_max_* = *h* by performing weighted sampling on the 50 best networks inferred with *h_max_* = *h*−1. These weights are selected so that the single best network and those similar to it in terms of log-likelihood are sampled more often. We hypothesized that this sampling strategy would be particularly useful with propQuartets=0.05 due to the increased variability in the quality of inferred networks when so few quartets are used.

The CUfast and CUfast500 search strategies yielded dramatically faster runtimes without sacrificing network accuracy. In fact, the best networks with h = 4, 5, 6, 7 were inferred using the CUfast500 search strategy, though these results do not alter the conclusions discussed in the main text. This is especially important because the total cumulative runtime of the CUfast500 search strategy was only 7.5% of the U-space search total cumulative runtime (Figure S14).

It is unclear why the CU-space search took dramatically longer than other spaces. We would have expected its runtime to be similar to the runtime of U-space, similar to how C-space runtime is similar to TCG-space runtime. These simulations were run on a shared computing cluster and CU-space was the final search space for which inference was performed, so it is possible that the cluster was under unusually heavy load during CU-space analyses, leading to the increased runtimes. Due to the drastic spike in runtimes, we only inferred networks with CU-space up to h = 6.

**Fig. S14:**
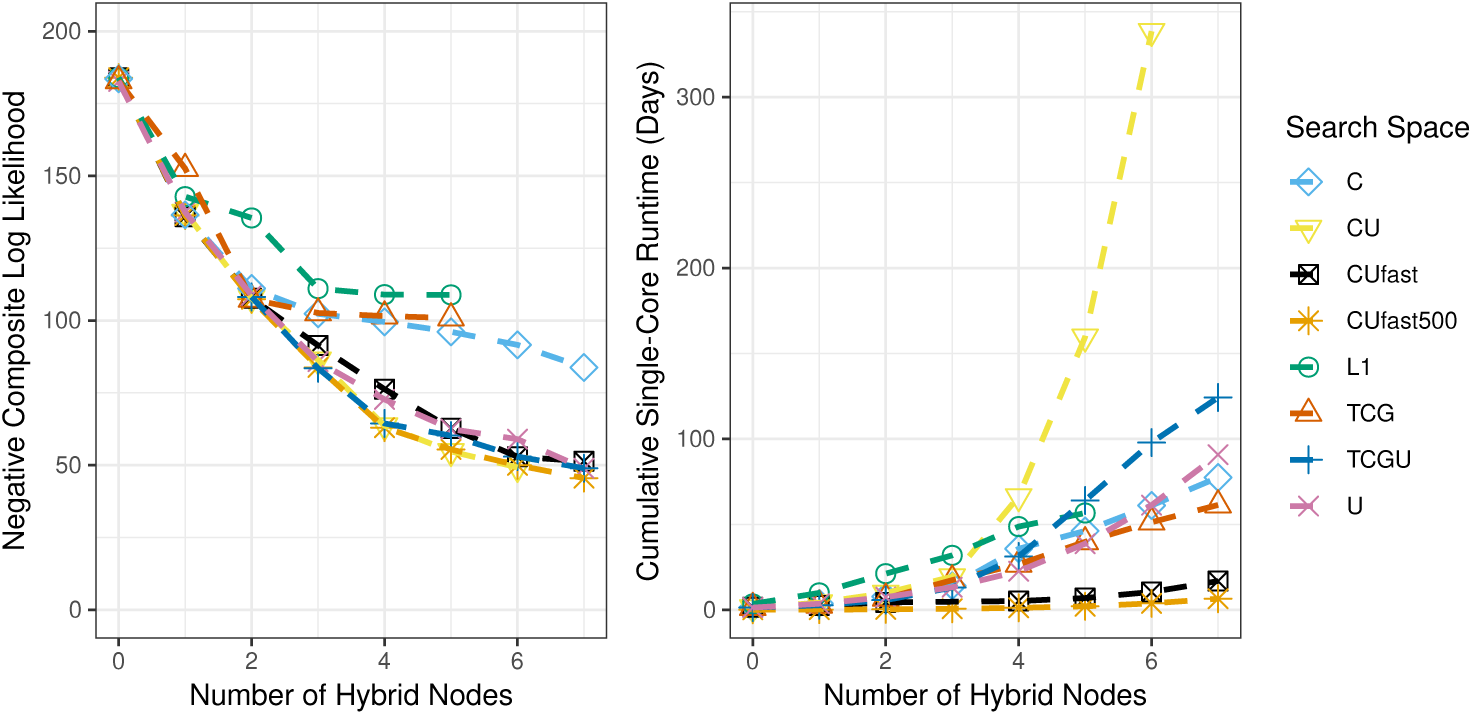
*Left:* Negative composite log-likelihood results with additional search strategies. *Right:* Cumulative runtimes in days to infer all networks under each search space and strategy with one processor core and two threads. For example, the cumulative runtime for a given space at *h* = 2 is the sum of all runtimes of each individual run ignoring parallelization across processors for *h* = 0, 1, 2. All spaces used 2 threads, except L1-space which was inferred with a version of SNaQ that did not feature parallelization across threads.

### C.2 DDSE Model Selection

**Fig. S15:**
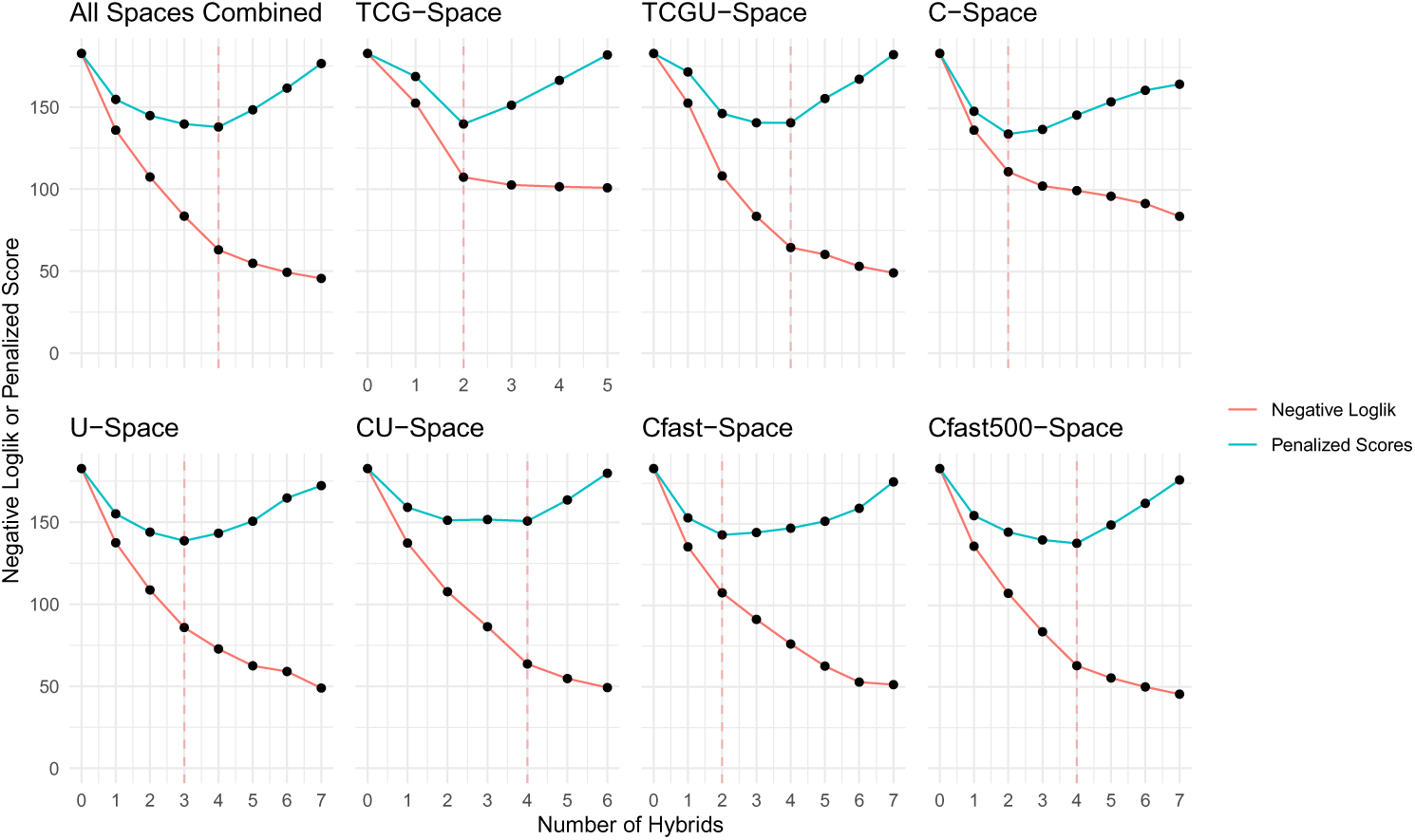
Data-driven slope estimation (DDSE) model selection curves used to select the best-fitting models from each set of inference spaces. **Solid red line:** best negative composite log-likelihood of all networks inferred with the given number of hybrids according to SNaQ.jl. **Solid blue line:** penalized score for each network calculated with DDSE using the CAPUSHE R package. **Dashed, red, horizontal line:** model with the lowest penalized score; the selected model.

### C.3 All Inferred Networks

Below are all of the best-fitting (highest NCLL) networks inferred with SNaQ.jl for each combination of parameter space and h_max_.

#### C.3.1 Level-1 Networks

**Fig. S16:**
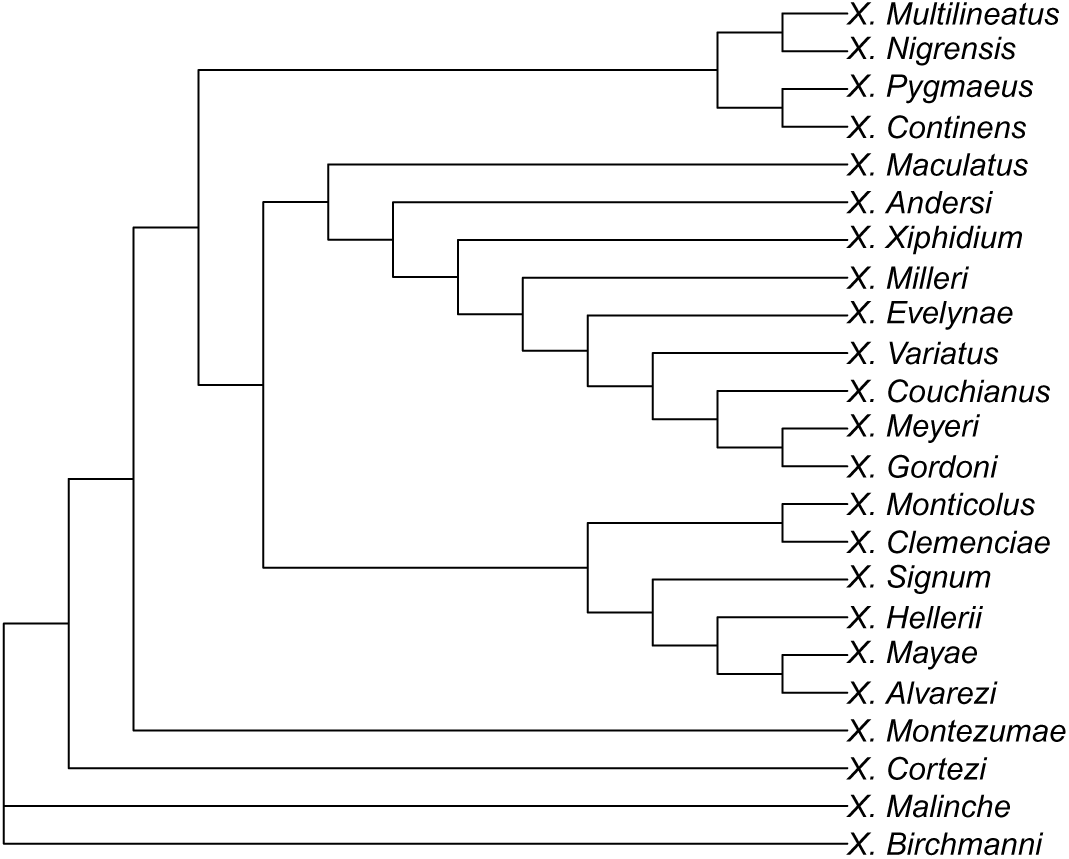
Level-1 (L1-space) network inferred for the phylogeny of *Xiphophorus* (Poeciliidae) with 0 hybridizations.

**Fig. S17:**
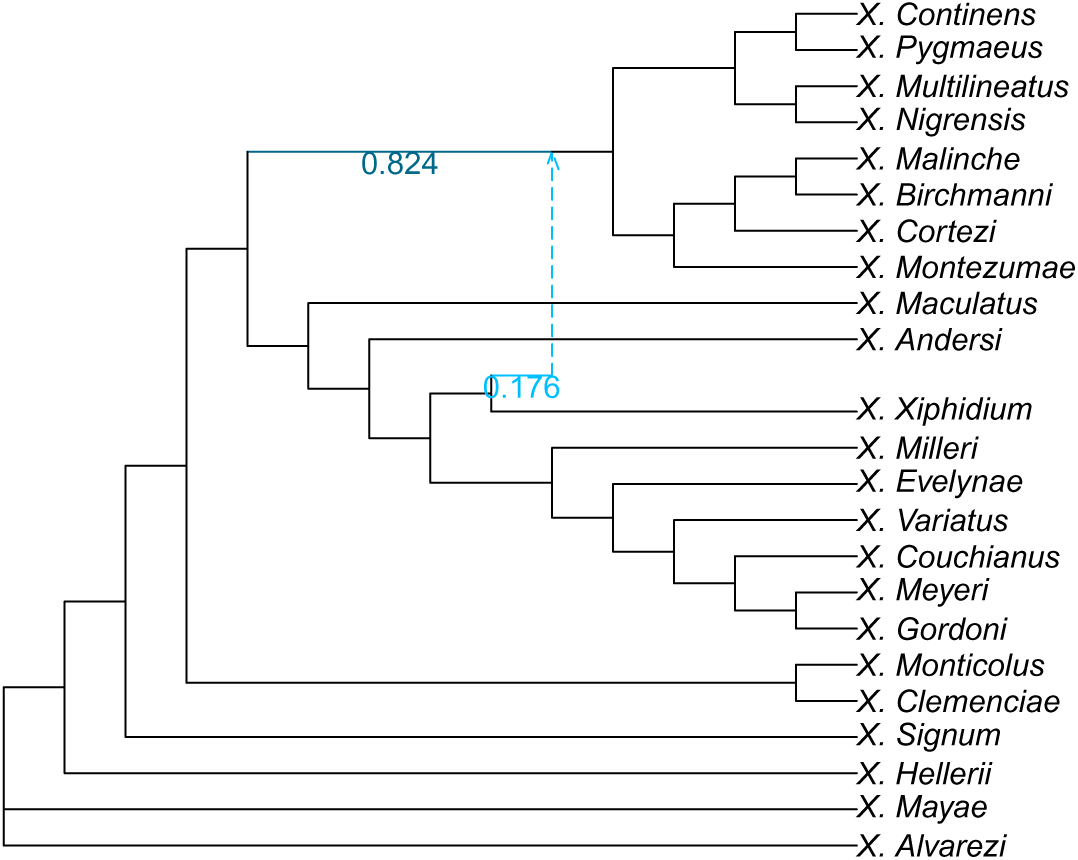
Level-1 (L1-space) network inferred for the phylogeny of *Xiphophorus* (Poeciliidae) with 1 hybridization.

**Fig. S18:**
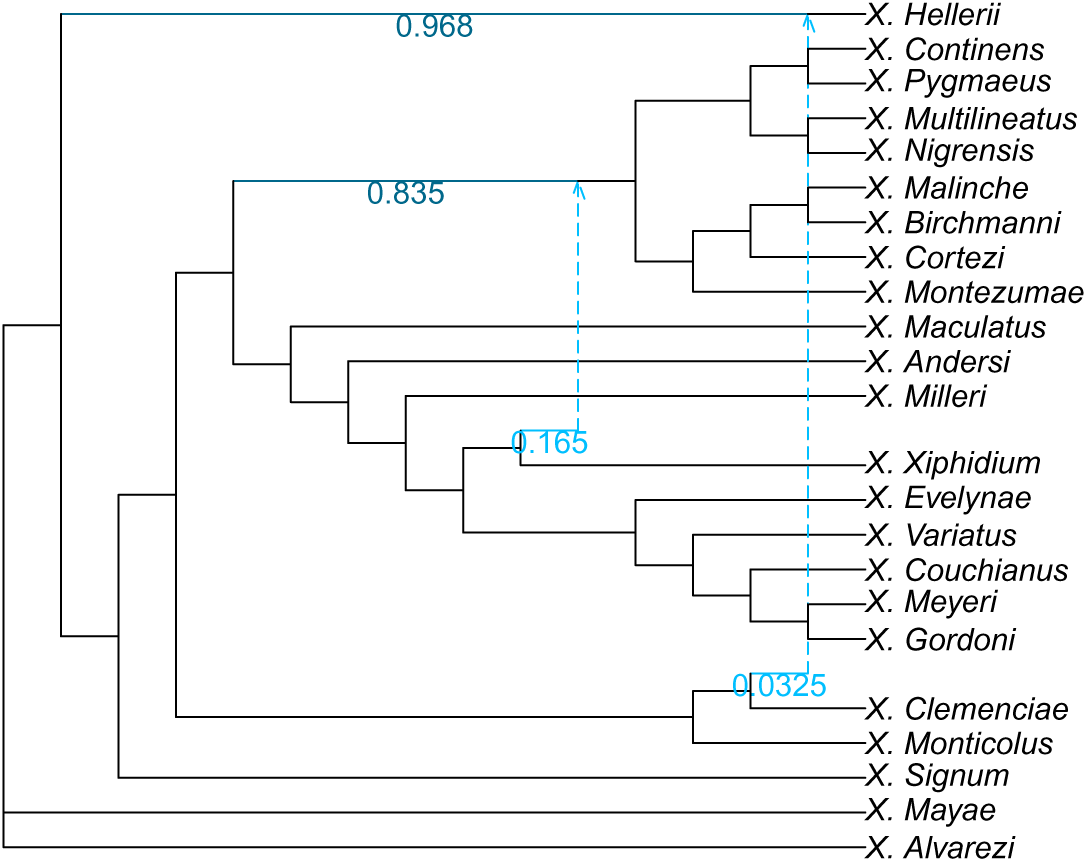
Level-1 (L1-space) network inferred for the phylogeny of *Xiphophorus* (Poeciliidae) with 2 hybridizations.

**Fig. S19:**
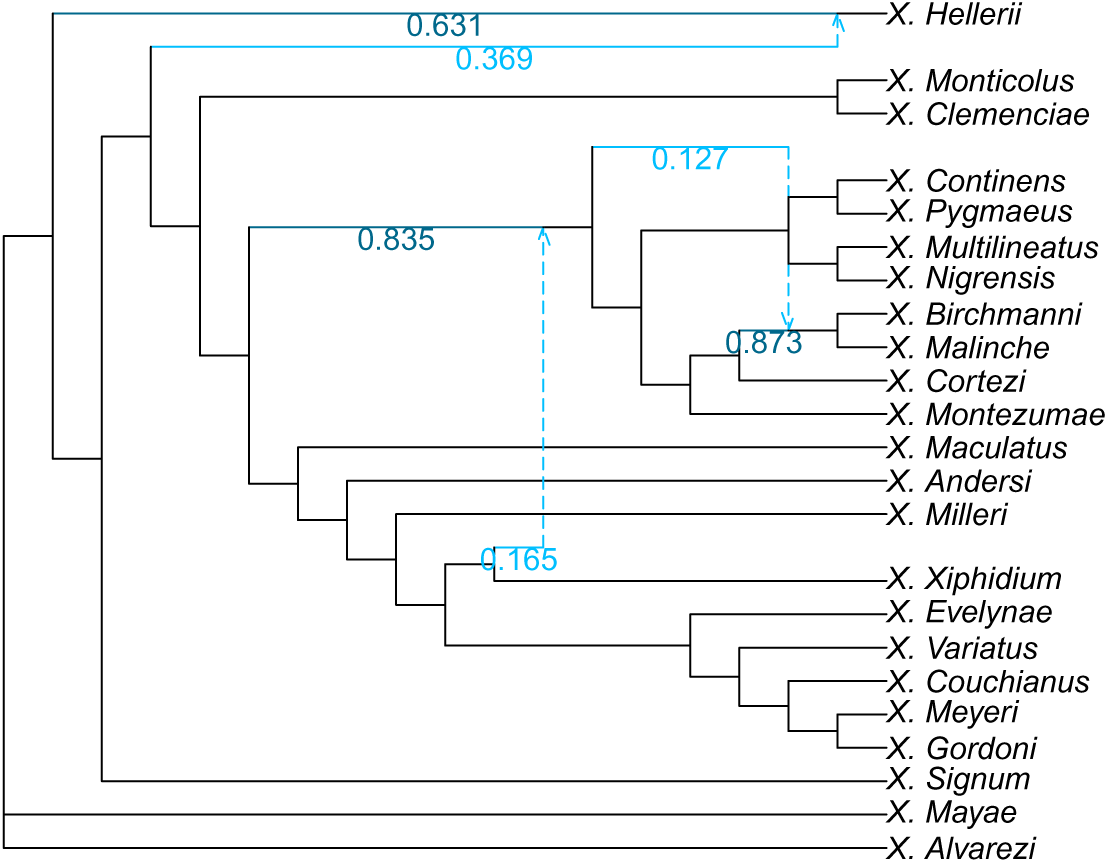
Level-1 (L1-space) network inferred for the phylogeny of *Xiphophorus* (Poeciliidae) with 3 hybridizations.

**Fig. S20:**
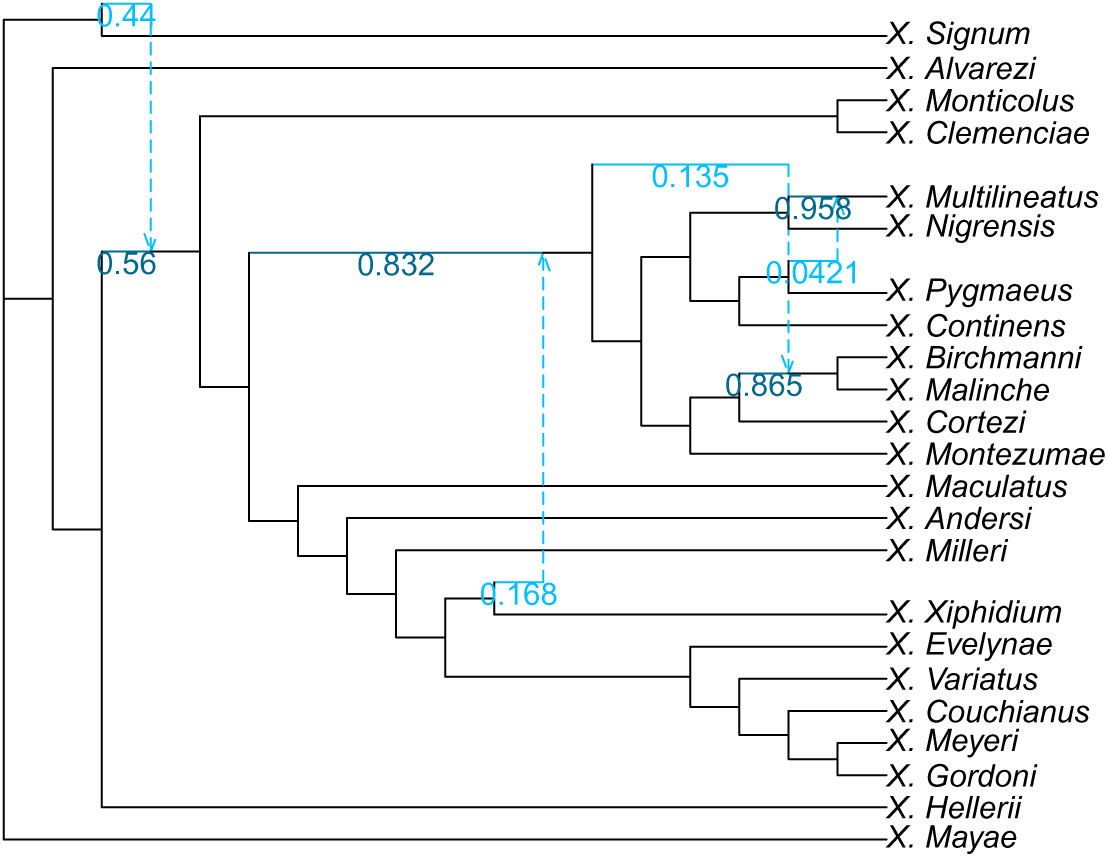
Level-1 (L1-space) network inferred for the phylogeny of *Xiphophorus* (Poeciliidae) with 4 hybridizations.

**Fig. S21:**
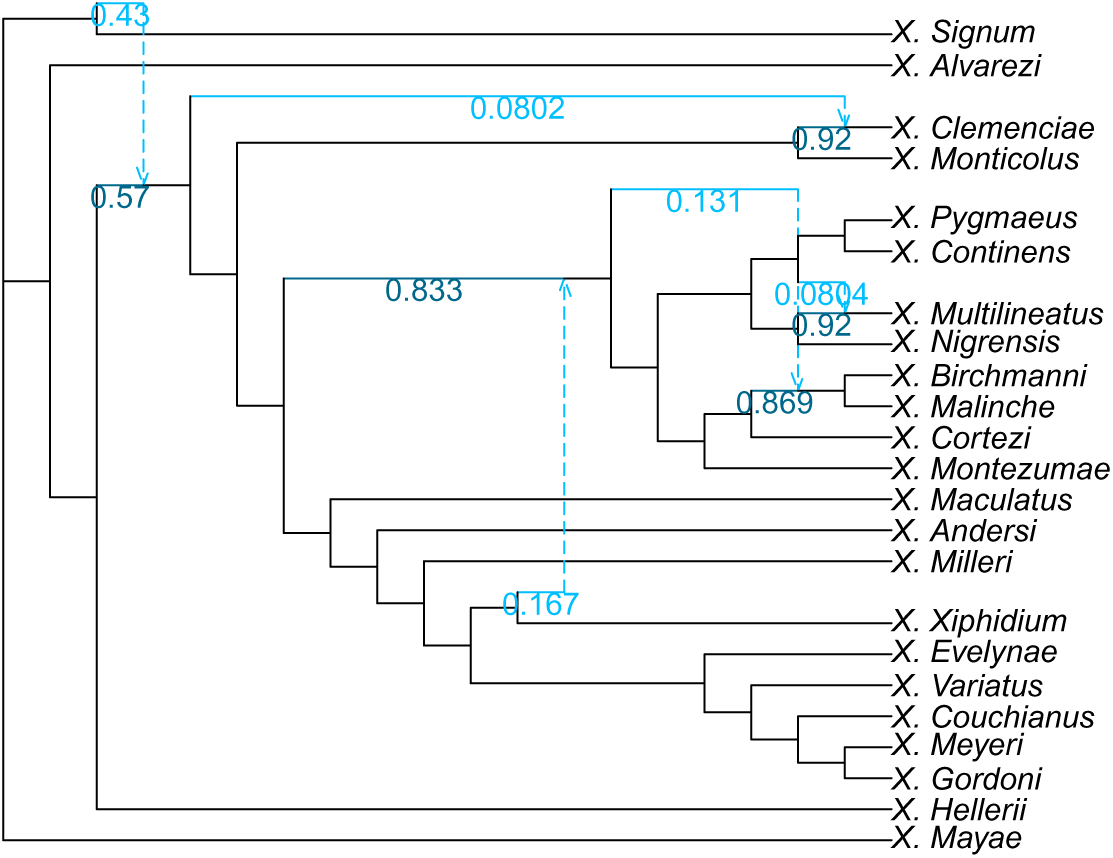
Level-1 (L1-space) network inferred for the phylogeny of *Xiphophorus* (Poeciliidae) with 5 hybridizations.

#### C.3.2 TCG-Space Networks

**Fig. S22:**
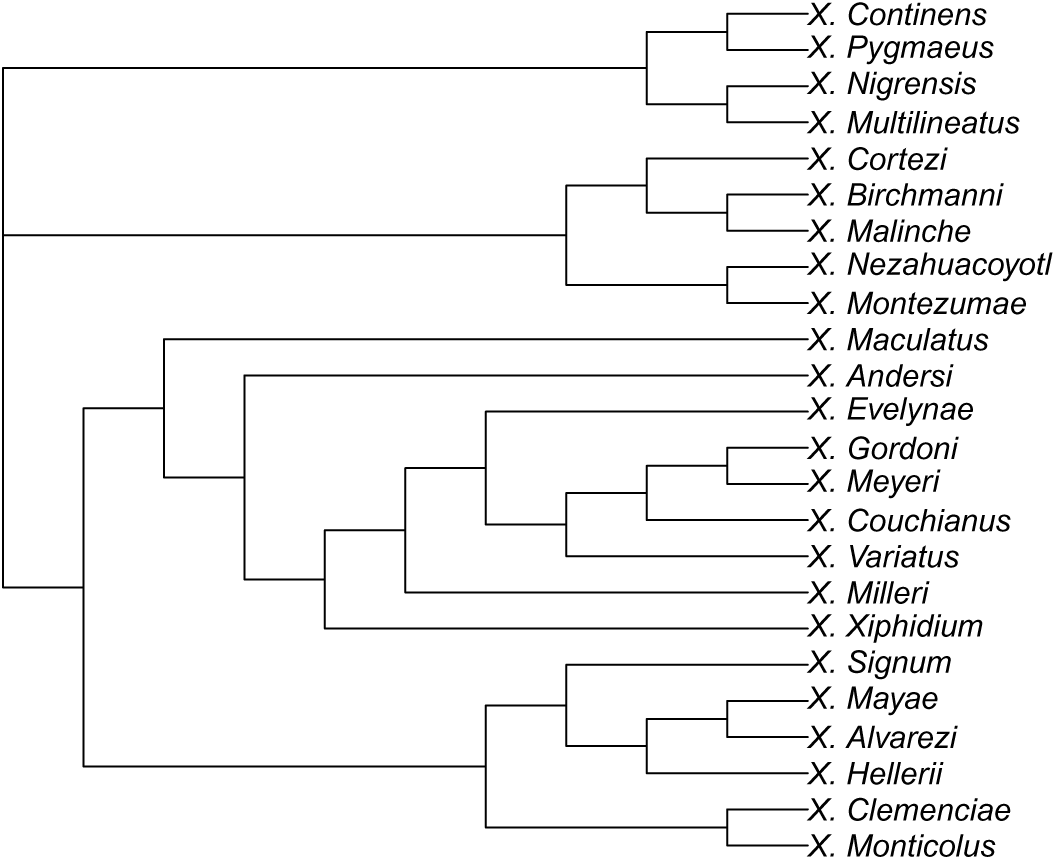
Tree-child and galled (TCG-space) network inferred for the phylogeny of *Xiphophorus* (Poeciliidae) with 0 hybridizations.

**Fig. S23:**
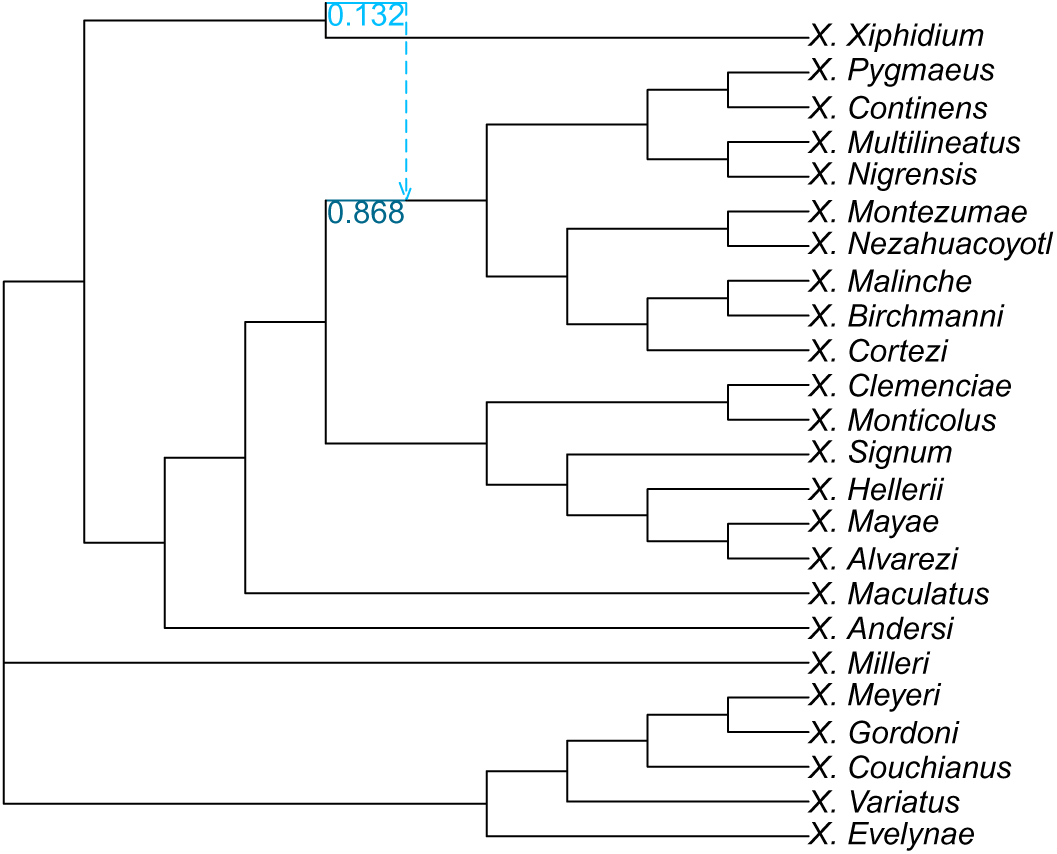
Tree-child and galled (TCG-space) network inferred for the phylogeny of *Xiphophorus* (Poeciliidae) with 1 hybridization.

**Fig. S24:**
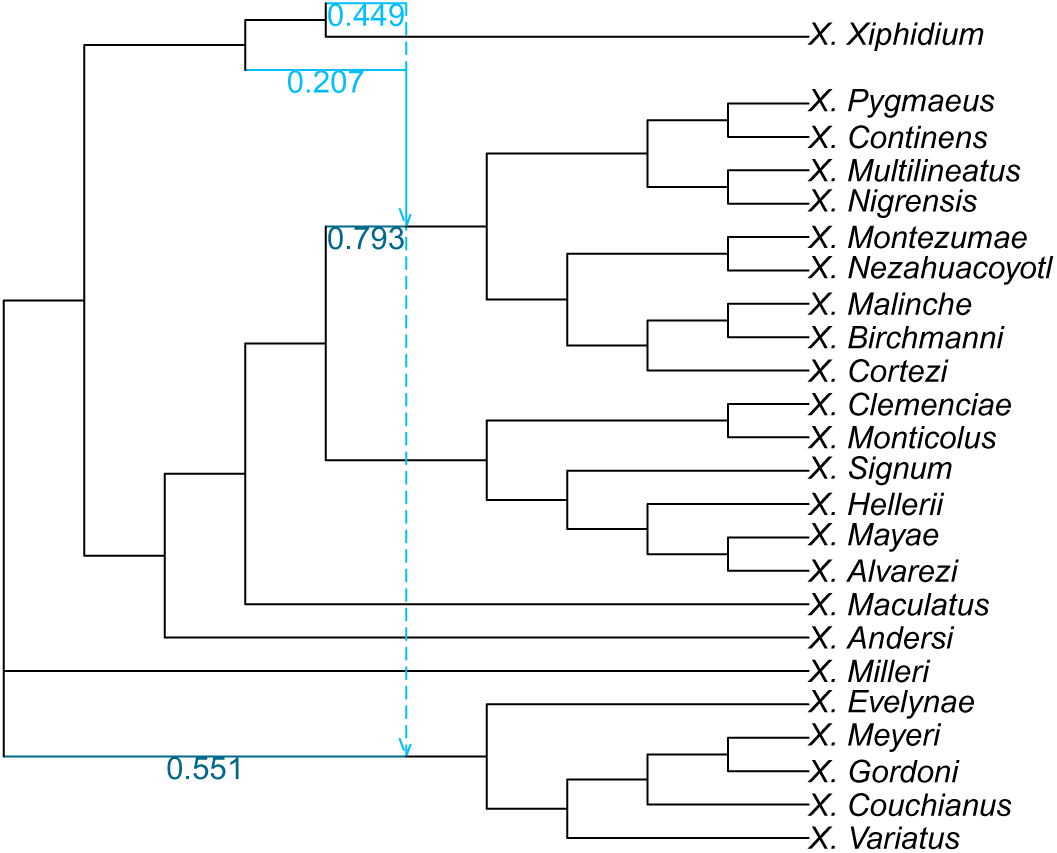
Tree-child and galled (TCG-space) network inferred for the phylogeny of *Xiphophorus* (Poeciliidae) with 2 hybridizations.

**Fig. S25:**
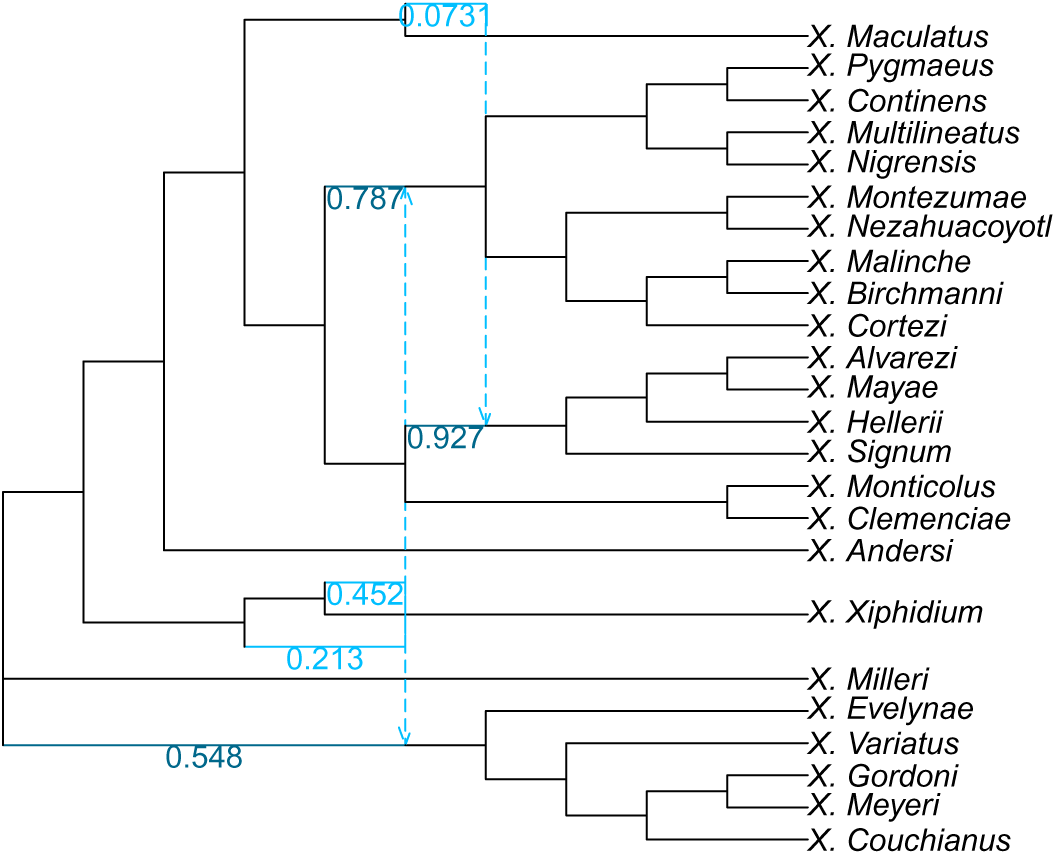
Tree-child and galled (TCG-space) network inferred for the phylogeny of *Xiphophorus* (Poeciliidae) with 3 hybridizations.

**Fig. S26:**
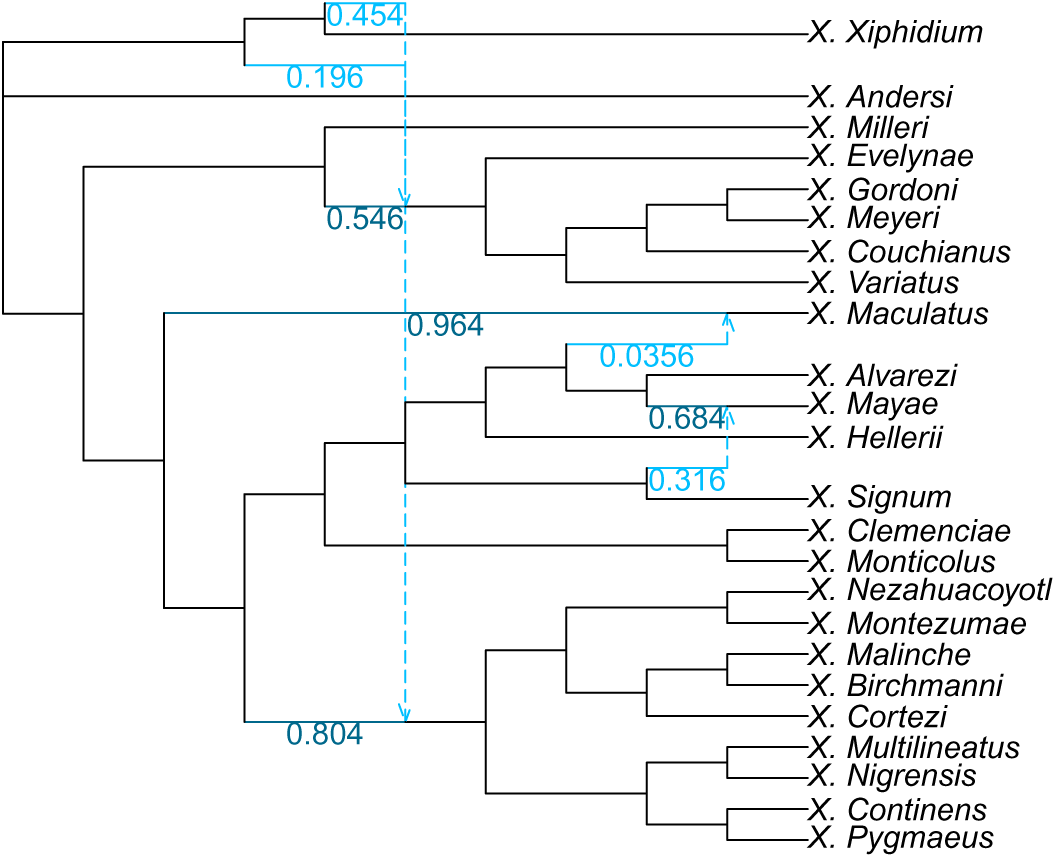
Tree-child and galled (TCG-space) network inferred for the phylogeny of *Xiphophorus* (Poeciliidae) with 4 hybridizations.

**Fig. S27:**
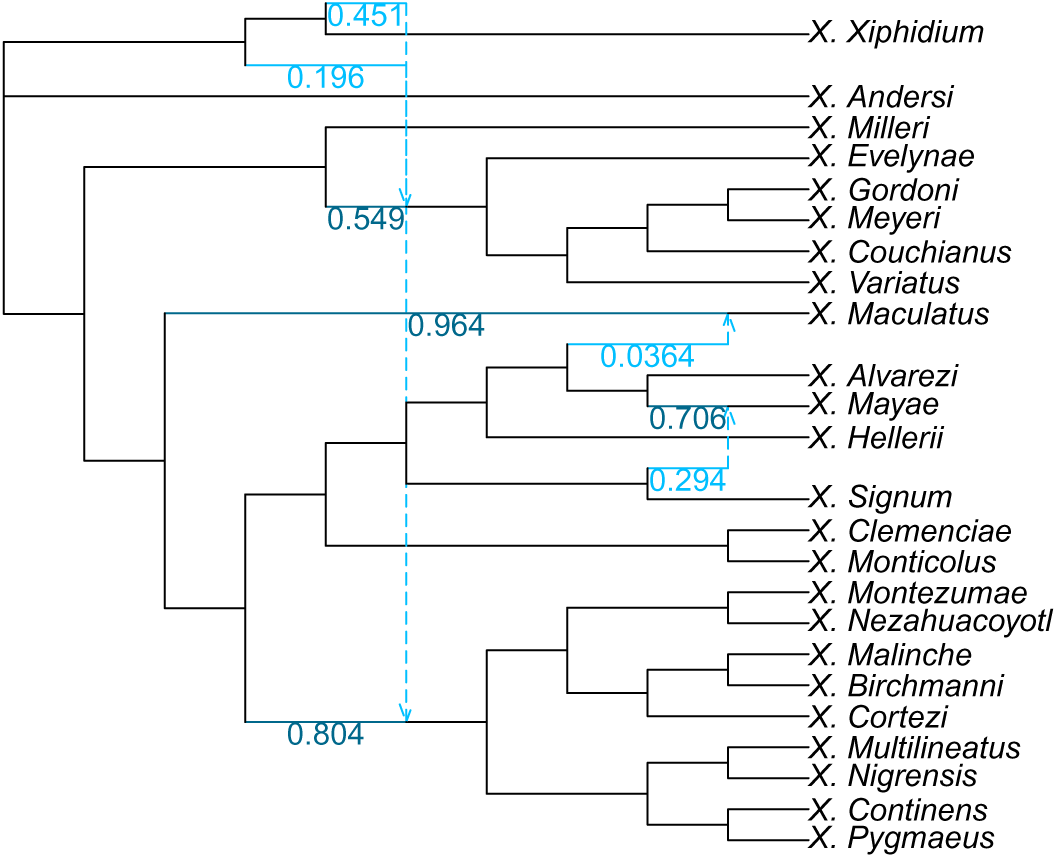
Tree-child and galled (TCG-space) network inferred for the phylogeny of *Xiphophorus* (Poeciliidae) with 5 hybridizations.

#### C.3.3 TCGU-Space Networks

**Fig. S28:**
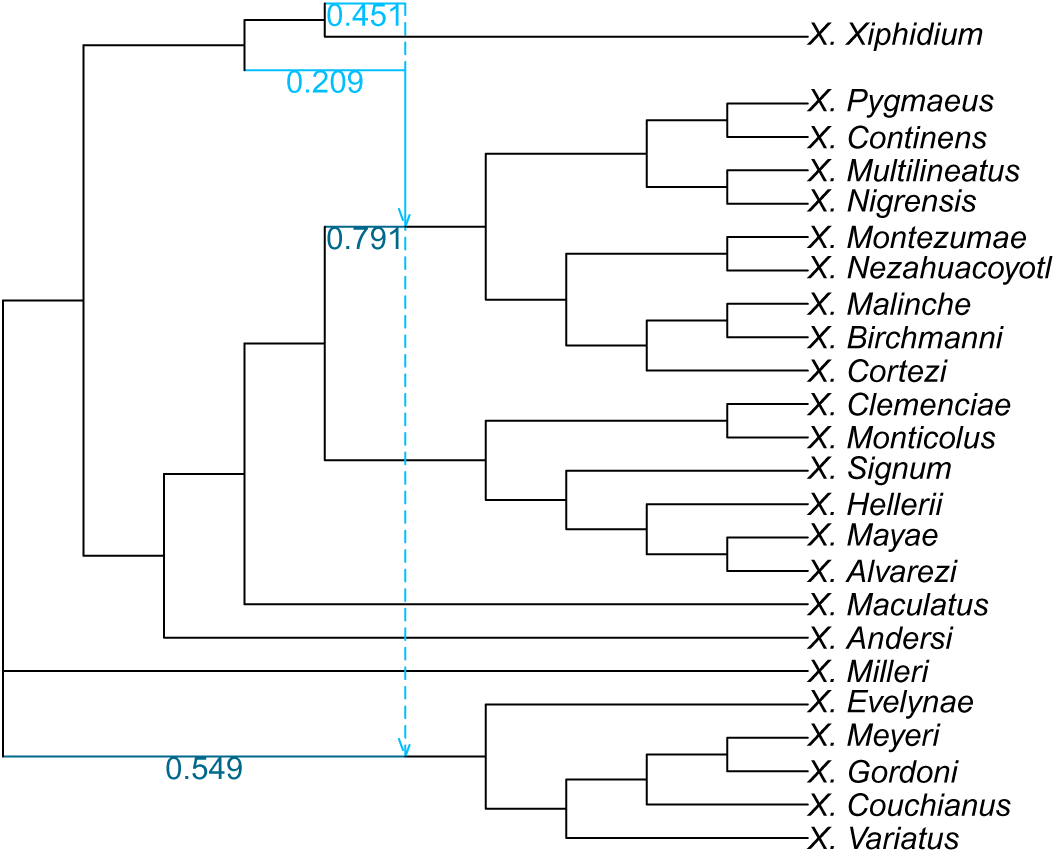
Unrestricted search network that used the corresponding TCG-space network as its starting point (TCGU-space) inferred for the phylogeny of *Xiphophorus* (Poeciliidae) with 2 hybridizations.

**Fig. S29:**
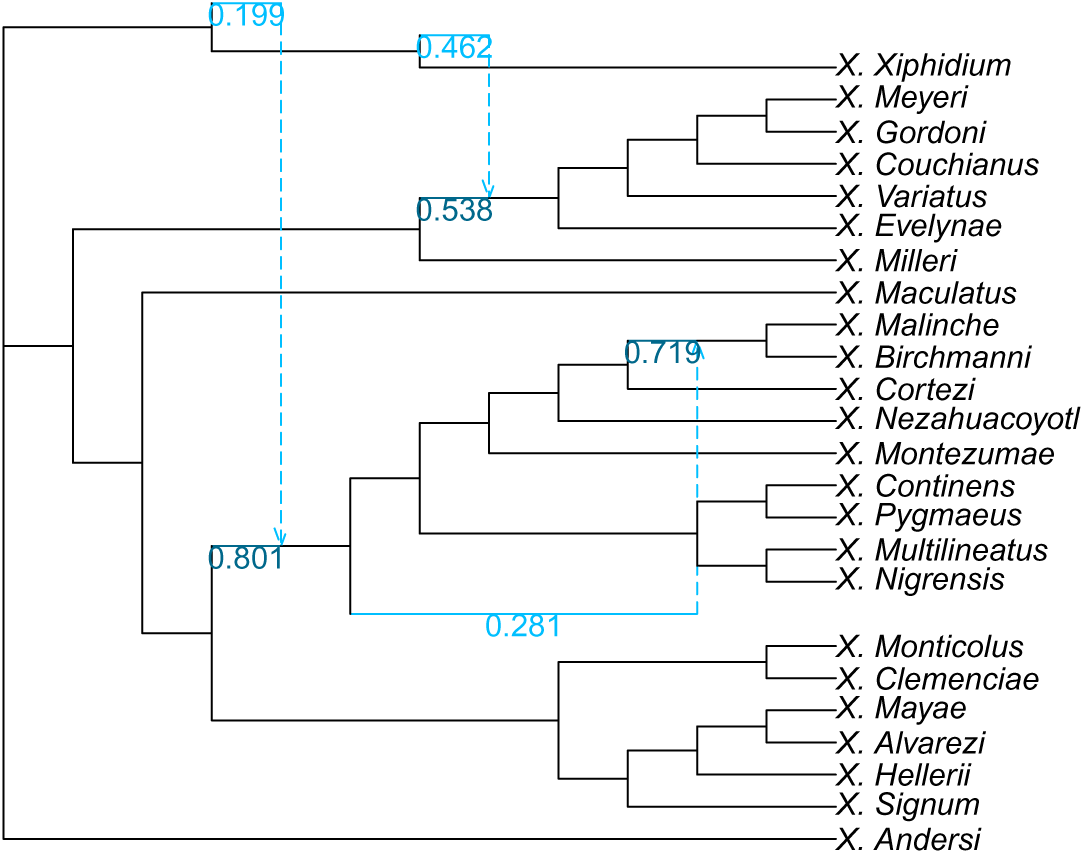
Unrestricted search network that used the corresponding TCG-space network as its starting point (TCGU-space) inferred for the phylogeny of *Xiphophorus* (Poeciliidae) with 3 hybridizations.

**Fig. S30:**
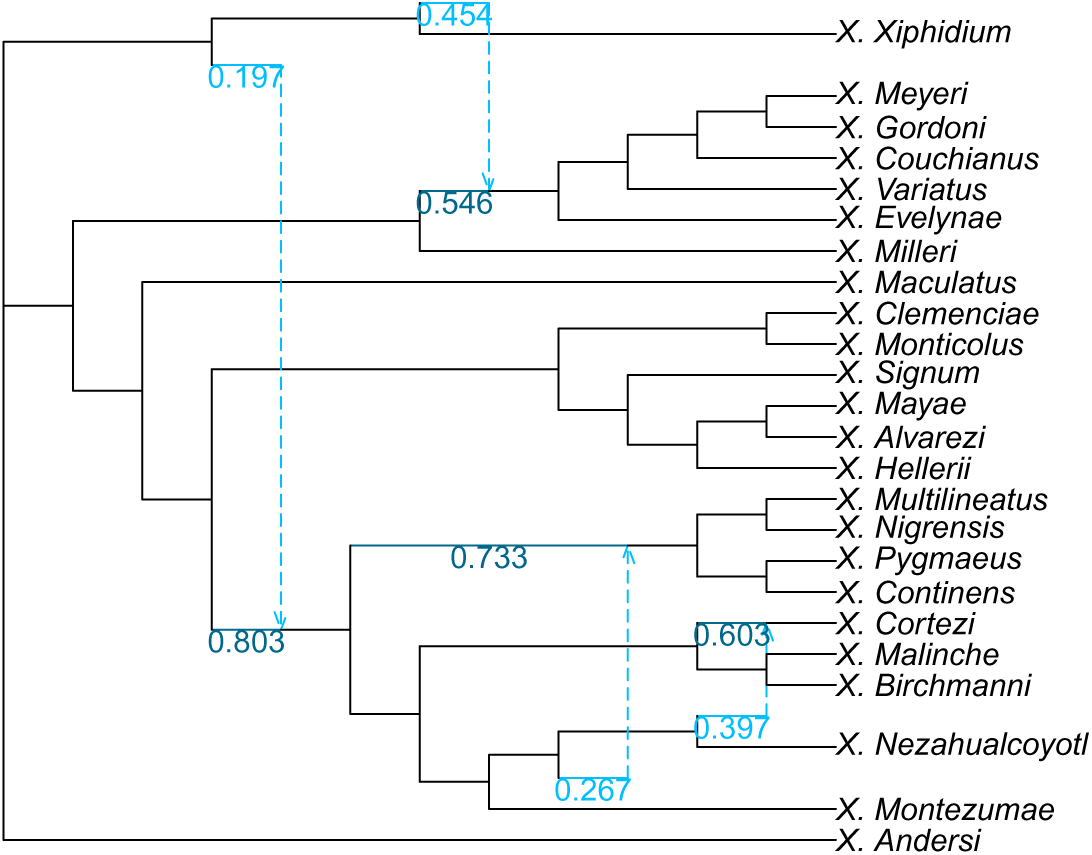
Unrestricted search network that used the corresponding TCG-space network as its starting point (TCGU-space) inferred for the phylogeny of *Xiphophorus* (Poeciliidae) with 4 hybridizations.

**Fig. S31:**
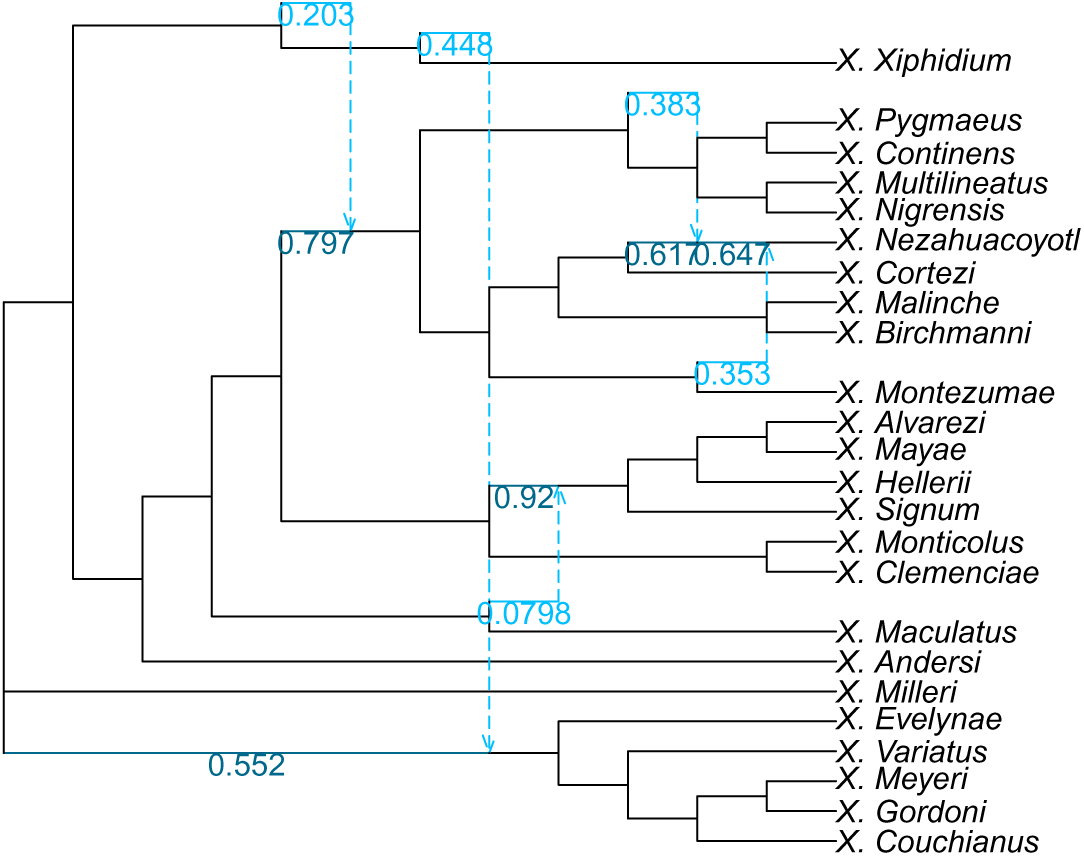
Unrestricted search network that used the corresponding TCG-space network as its starting point (TCGU-space) inferred for the phylogeny of *Xiphophorus* (Poeciliidae) with 5 hybridizations.

**Fig. S32:**
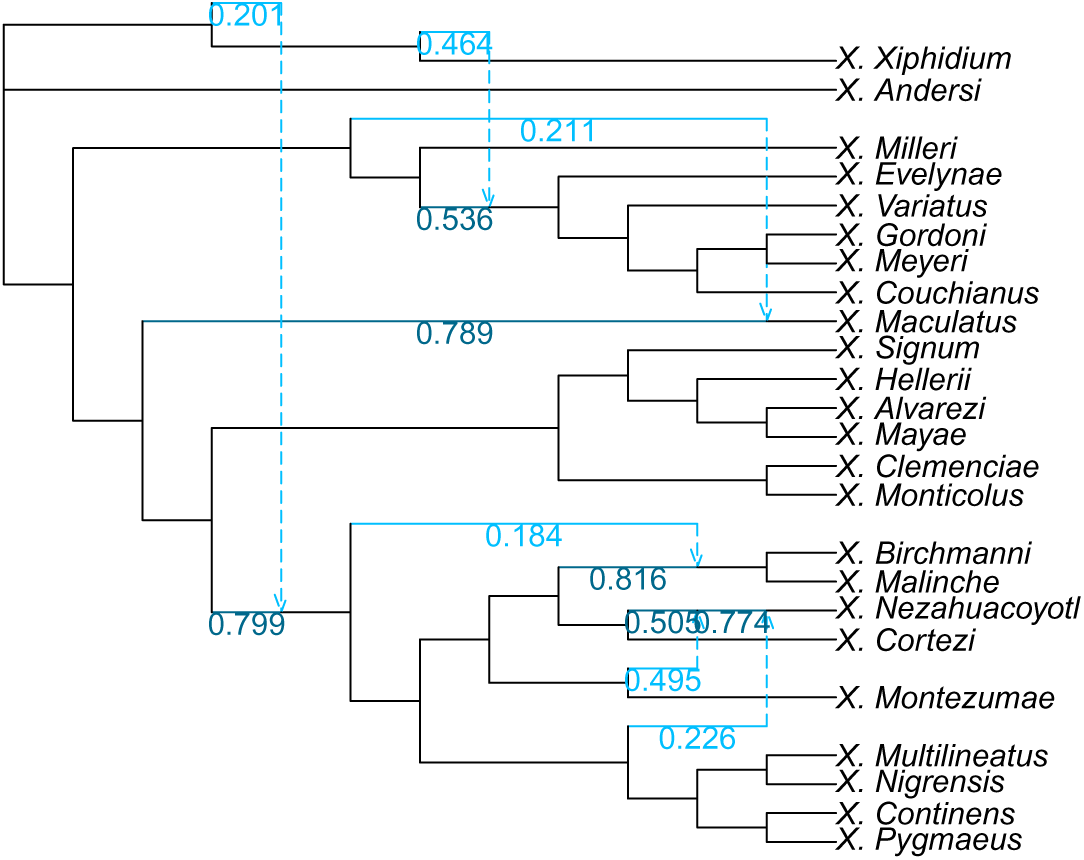
Unrestricted search network that used the corresponding TCG-space network as its starting point (TCGU-space) inferred for the phylogeny of *Xiphophorus* (Poeciliidae) with 6 hybridizations.

**Fig. S33:**
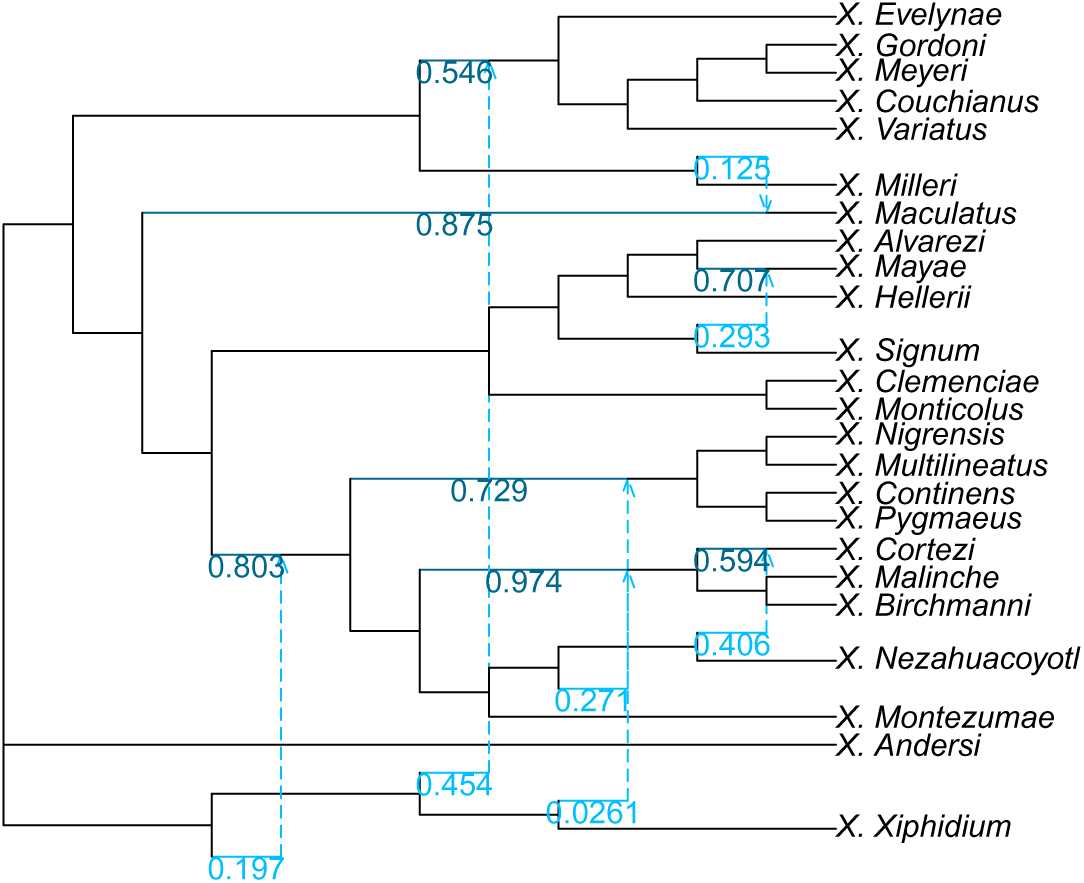
Unrestricted search network that used the corresponding TCG-space network as its starting point (TCGU-space) inferred for the phylogeny of *Xiphophorus* (Poeciliidae) with 7 hybridizations.

#### C.3.4 U-Space Networks

**Fig. S34:**
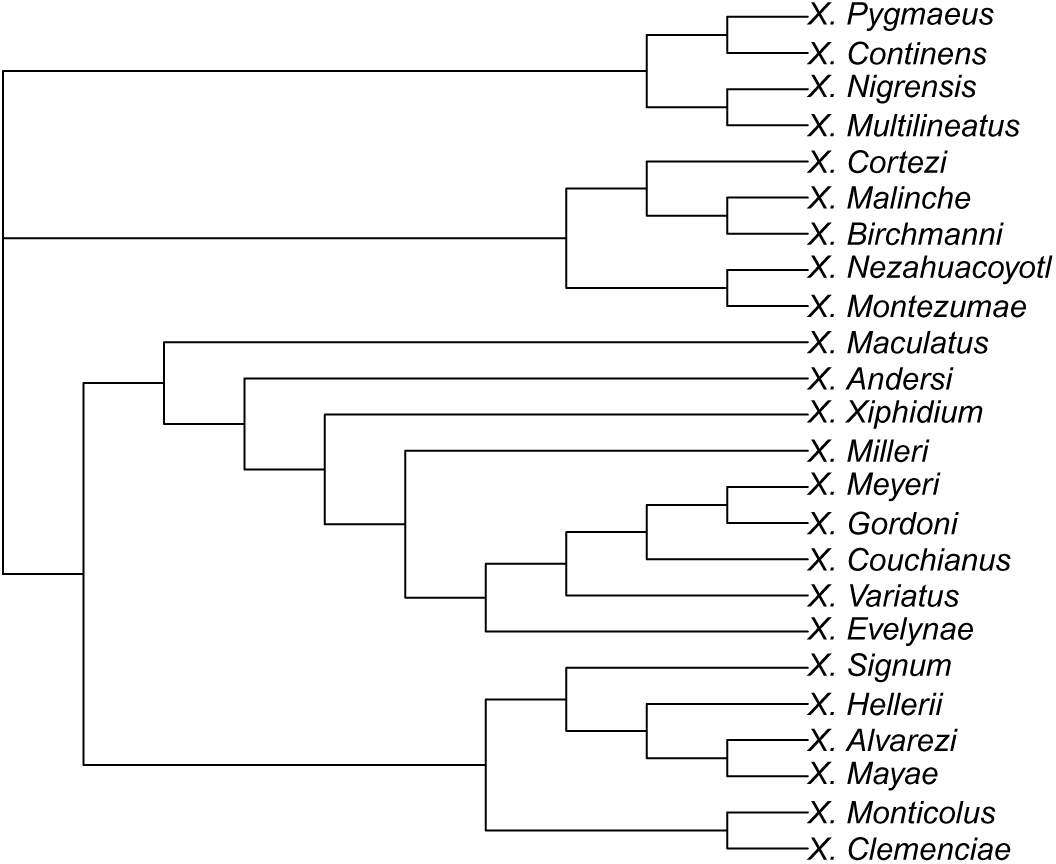
Unrestricted (U-space) network inferred for the phylogeny of *Xiphophorus* (Poeciliidae) with 0 hybridizations.

**Fig. S35:**
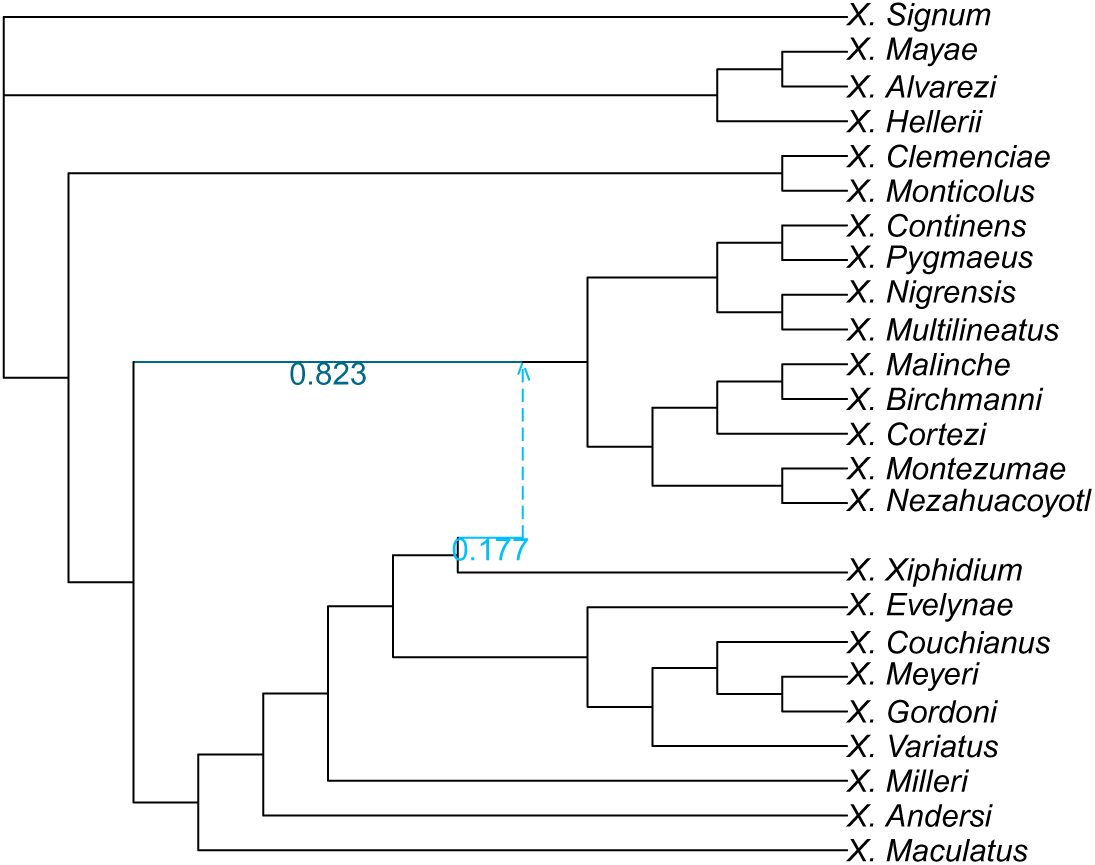
Unrestricted (U-space) network inferred for the phylogeny of *Xiphophorus* (Poeciliidae) with 1 hybridization.

**Fig. S36:**
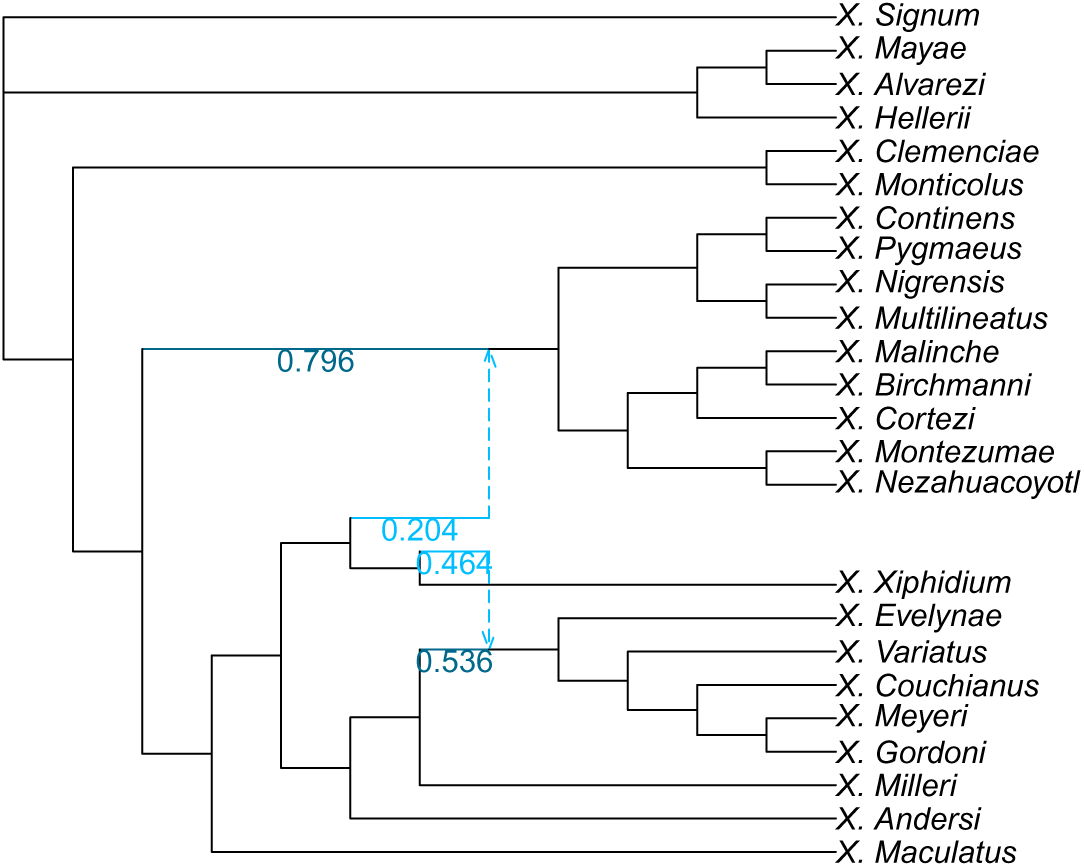
Unrestricted (U-space) network inferred for the phylogeny of *Xiphophorus* (Poeciliidae) with 2 hybridizations.

**Fig. S37:**
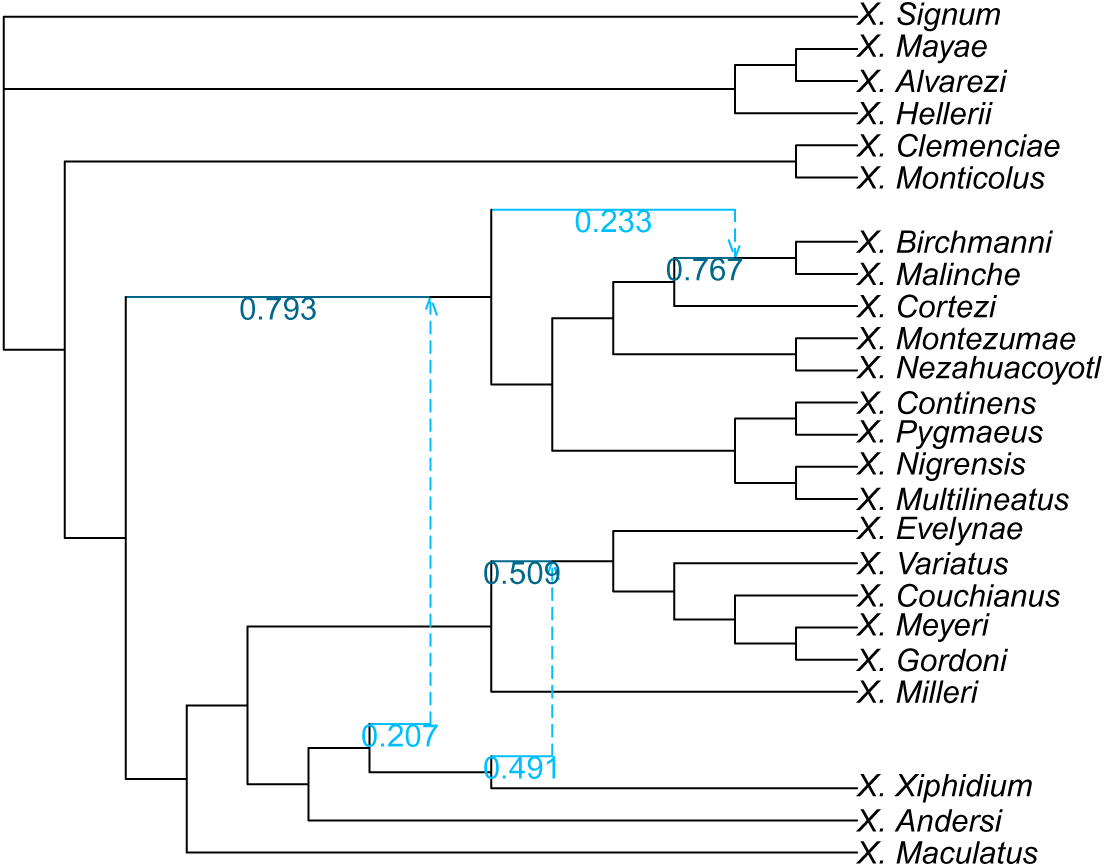
Unrestricted (U-space) network inferred for the phylogeny of *Xiphophorus* (Poeciliidae) with 3 hybridizations.

**Fig. S38:**
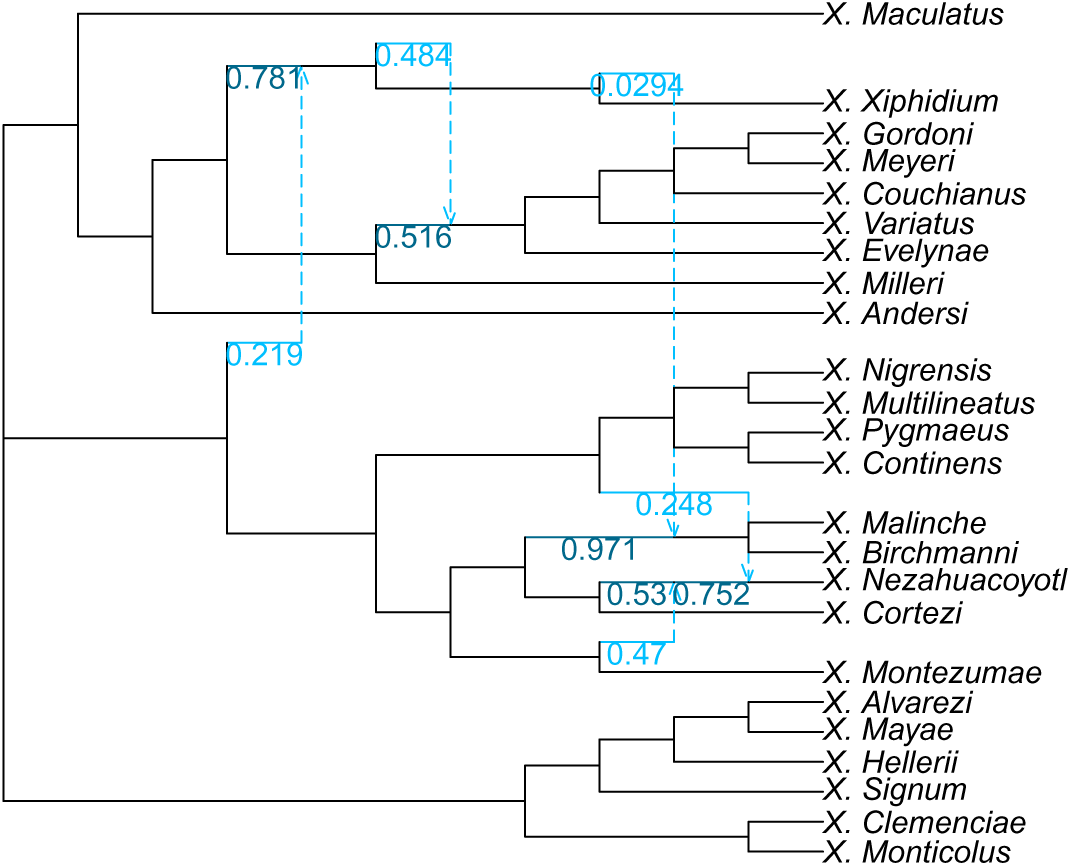
Unrestricted (U-space) network inferred for the phylogeny of *Xiphophorus* (Poeciliidae) with 4 hybridizations.

**Fig. S39:**
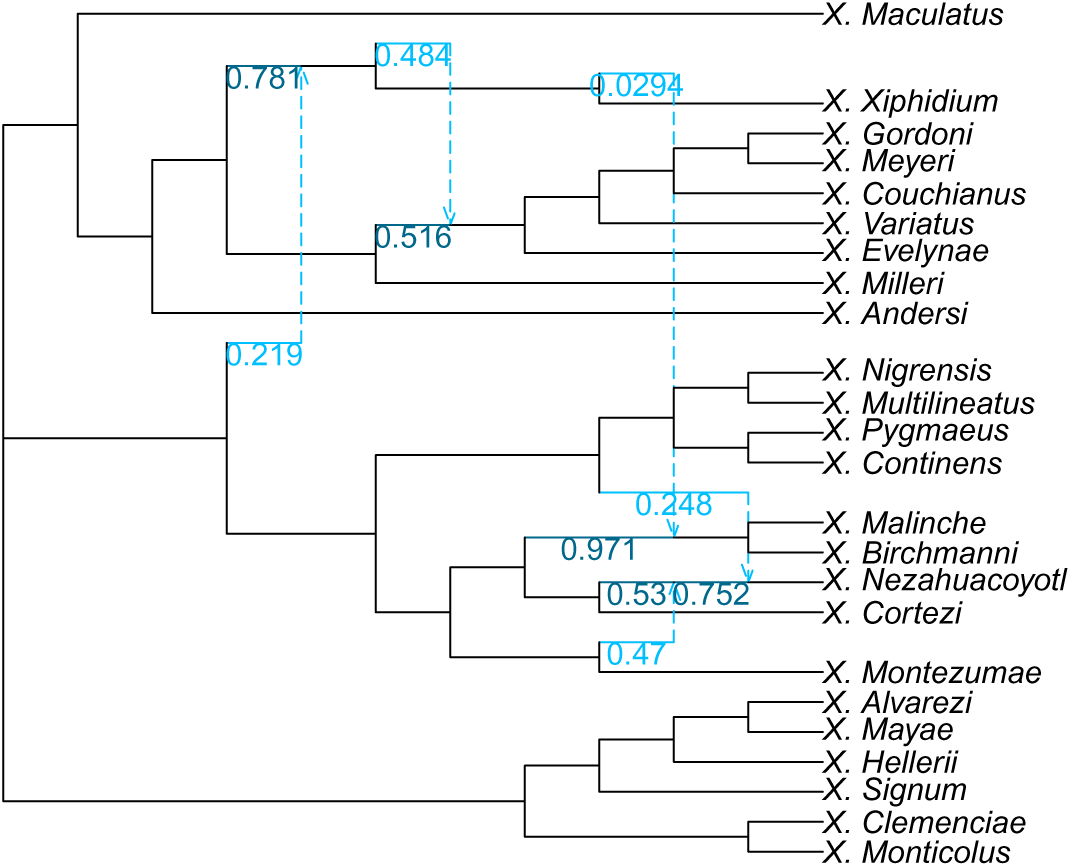
Unrestricted (U-space) network inferred for the phylogeny of *Xiphophorus* (Poeciliidae) with 5 hybridizations.

**Fig. S40:**
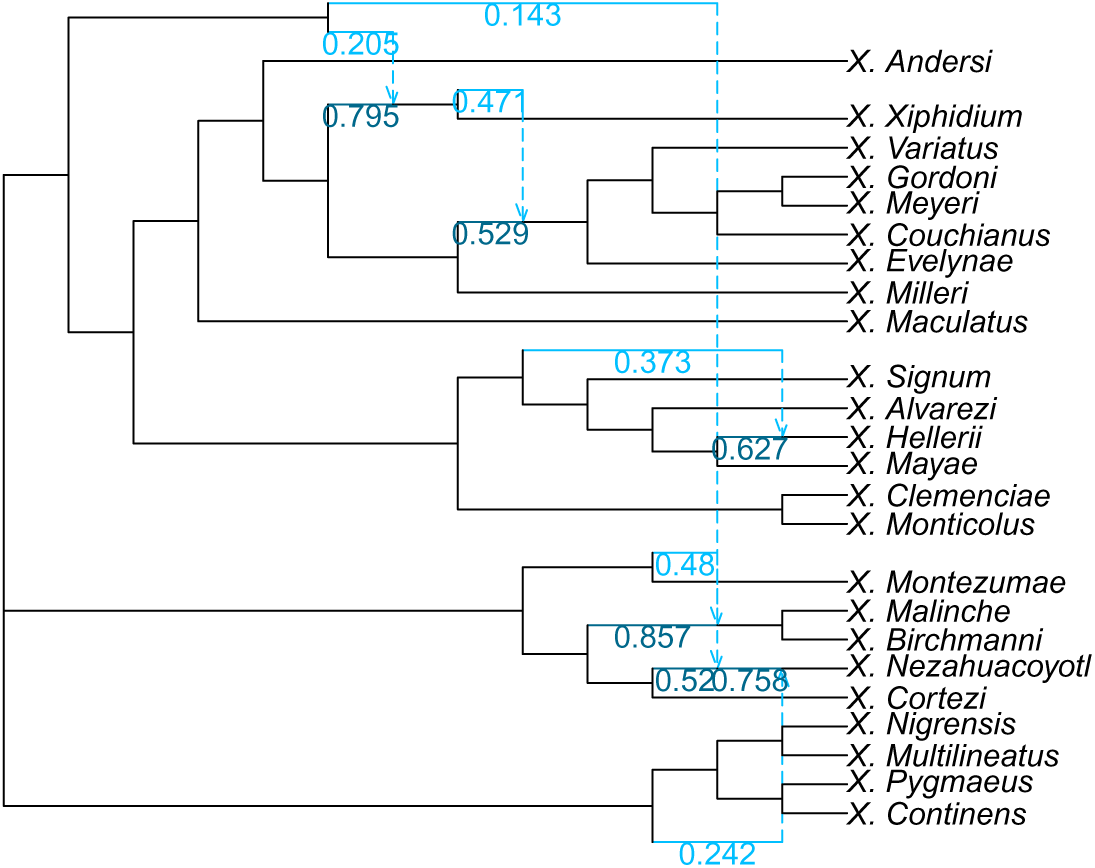
Unrestricted (U-space) network inferred for the phylogeny of *Xiphophorus* (Poeciliidae) with 6 hybridizations.

**Fig. S41:**
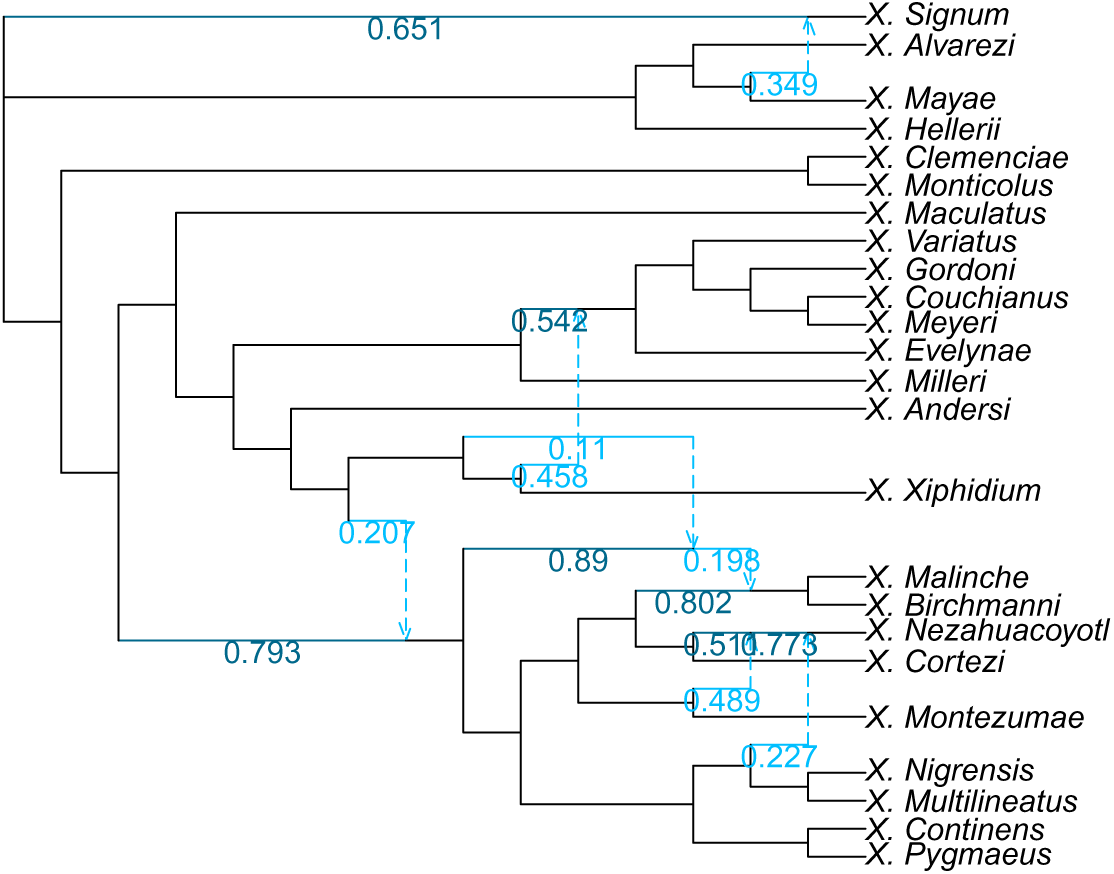
Unrestricted (U-space) network inferred for the phylogeny of *Xiphophorus* (Poeciliidae) with 7 hybridizations.

#### C.3.5 C-Space Networks

**Fig. S42:**
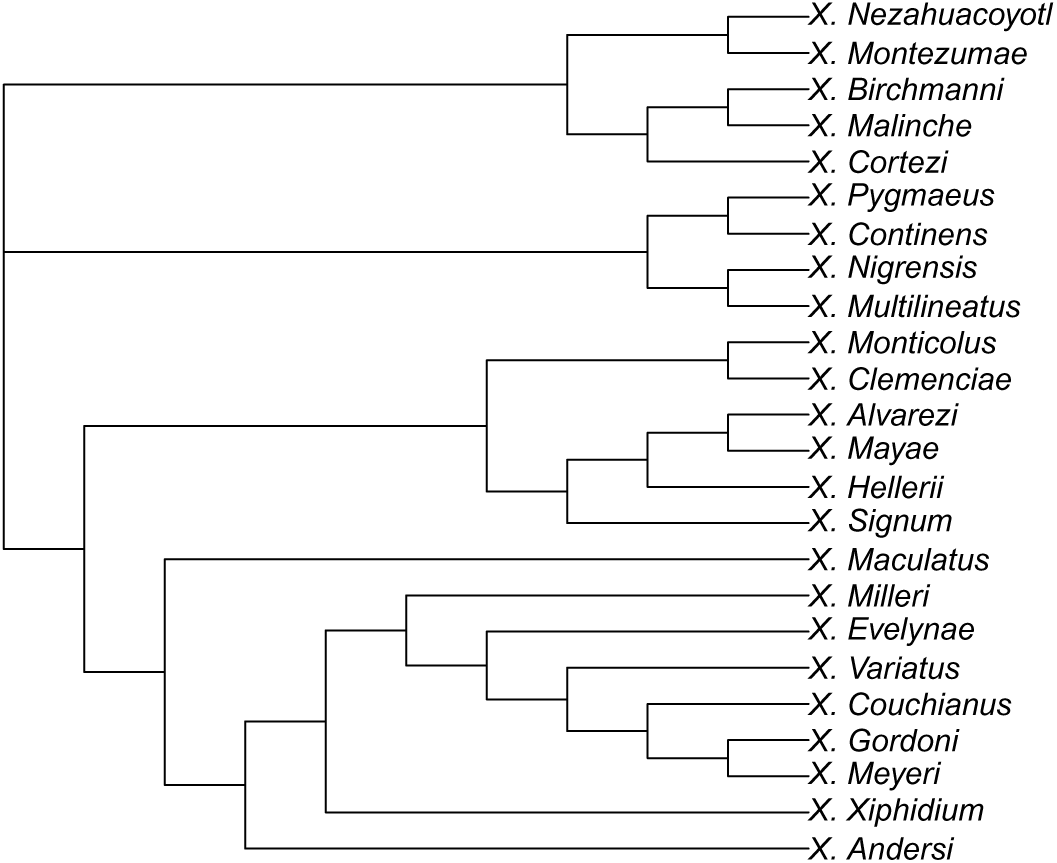
Clade-restricted (C-space) network inferred for the phylogeny of *Xiphophorus* (Poeciliidae) with 0 hybridizations.

**Fig. S43:**
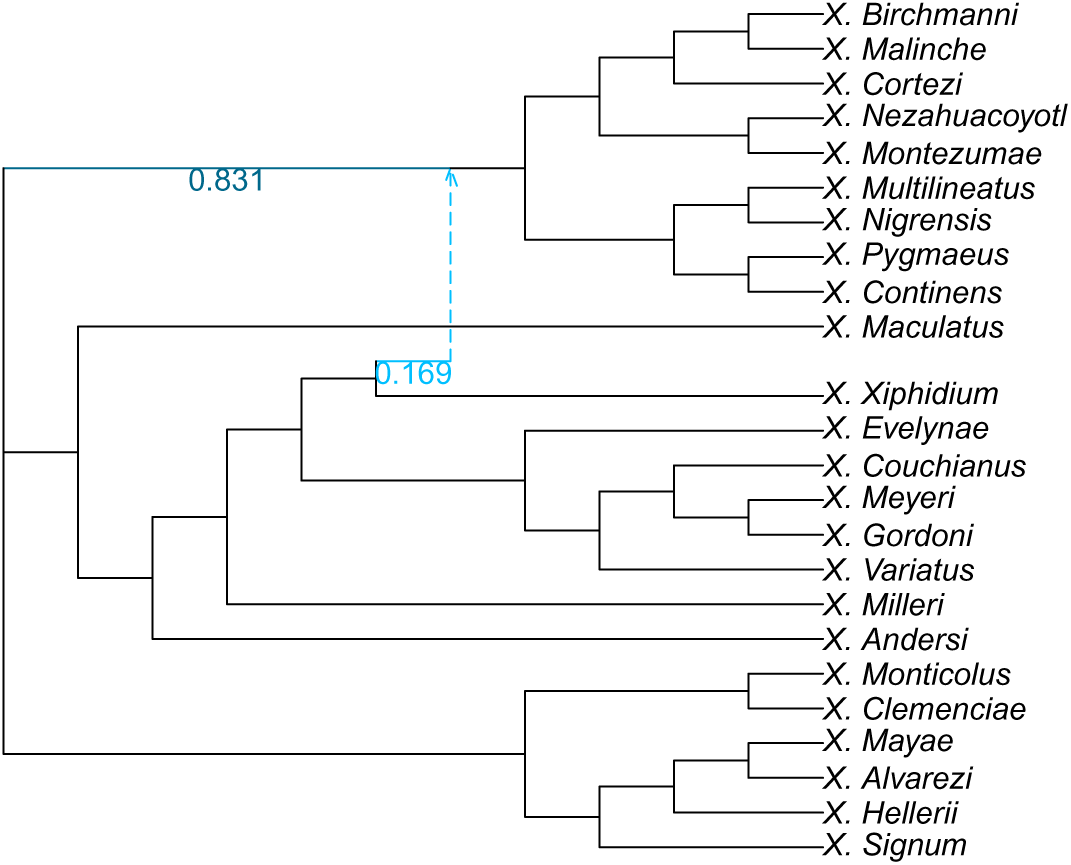
Clade-restricted (C-space) network inferred for the phylogeny of *Xiphophorus* (Poeciliidae) with 1 hybridization.

**Fig. S44:**
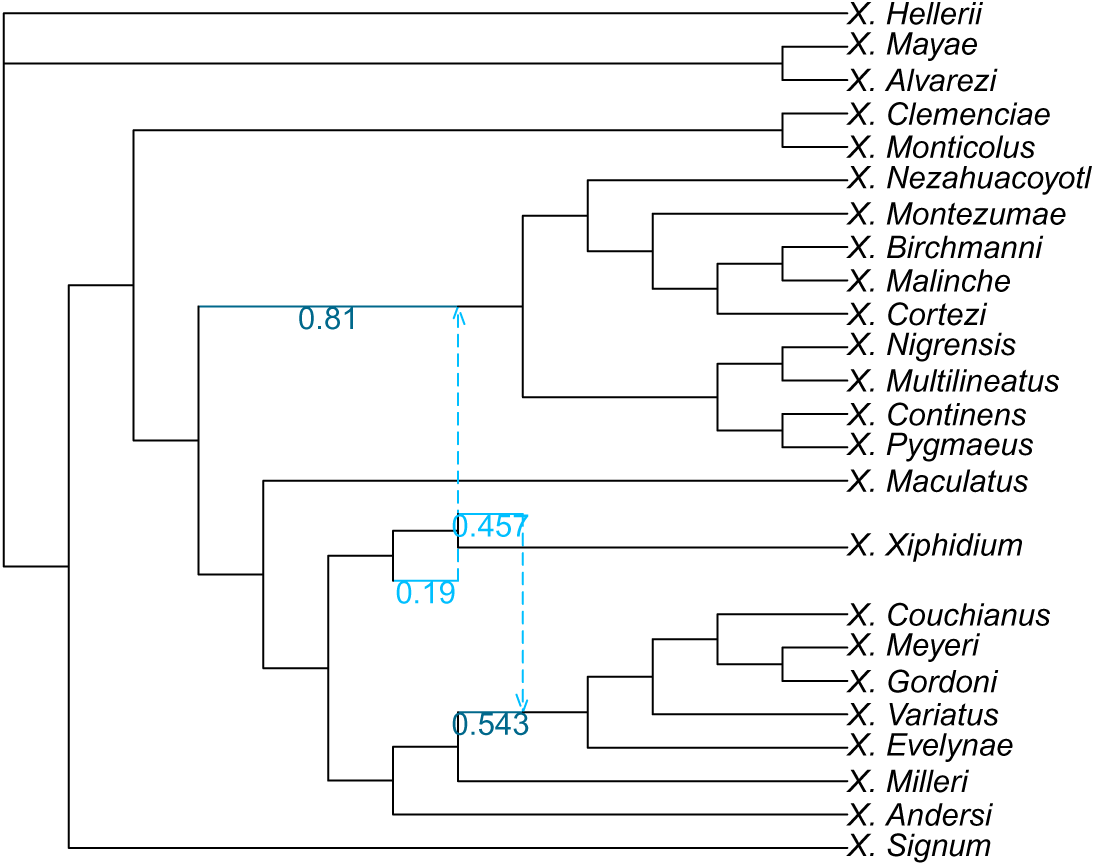
Clade-restricted (C-space) network inferred for the phylogeny of *Xiphophorus* (Poeciliidae) with 2 hybridizations.

**Fig. S45:**
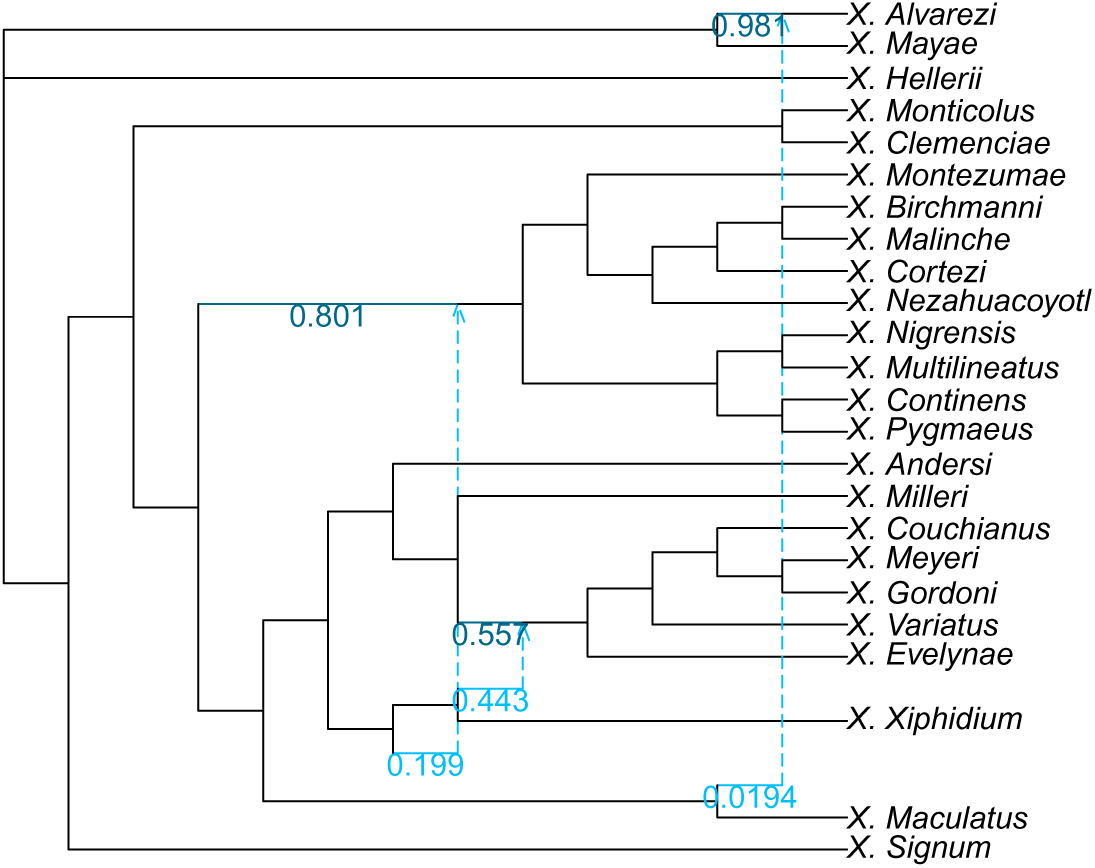
Clade-restricted (C-space) network inferred for the phylogeny of *Xiphophorus* (Poeciliidae) with 3 hybridizations.

**Fig. S46:**
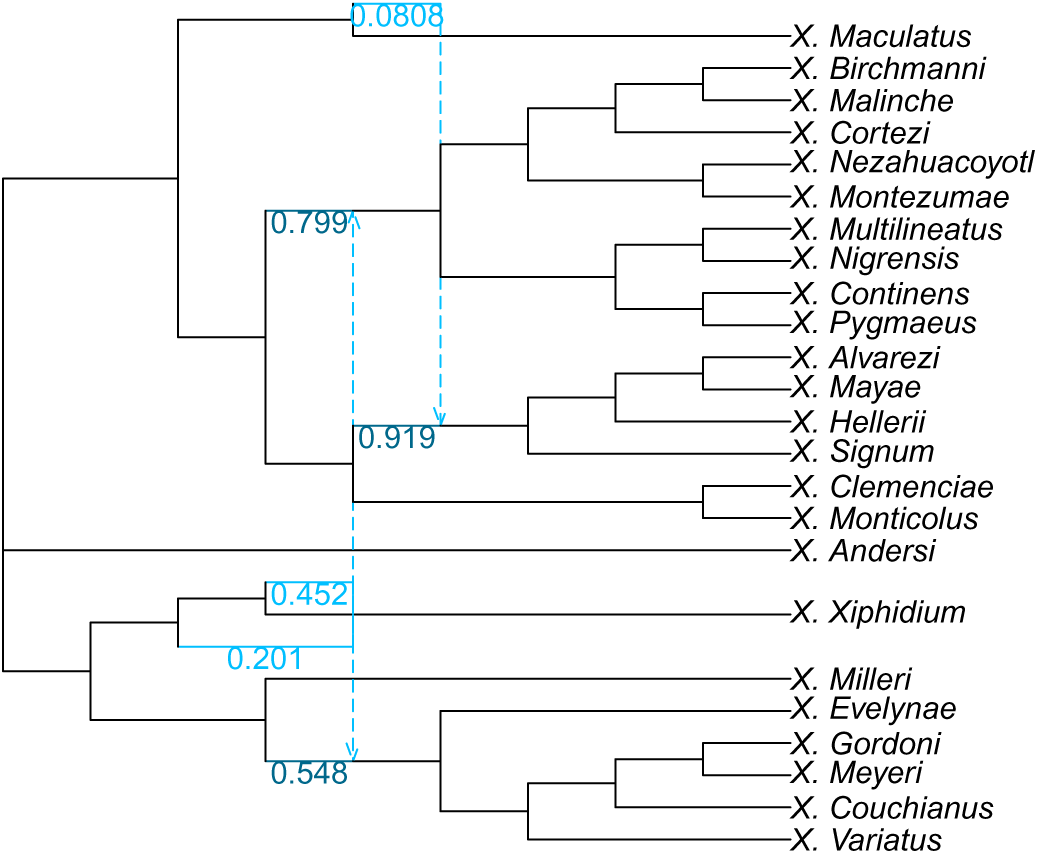
Clade-restricted (C-space) network inferred for the phylogeny of *Xiphophorus* (Poeciliidae) with 4 hybridizations.

**Fig. S47:**
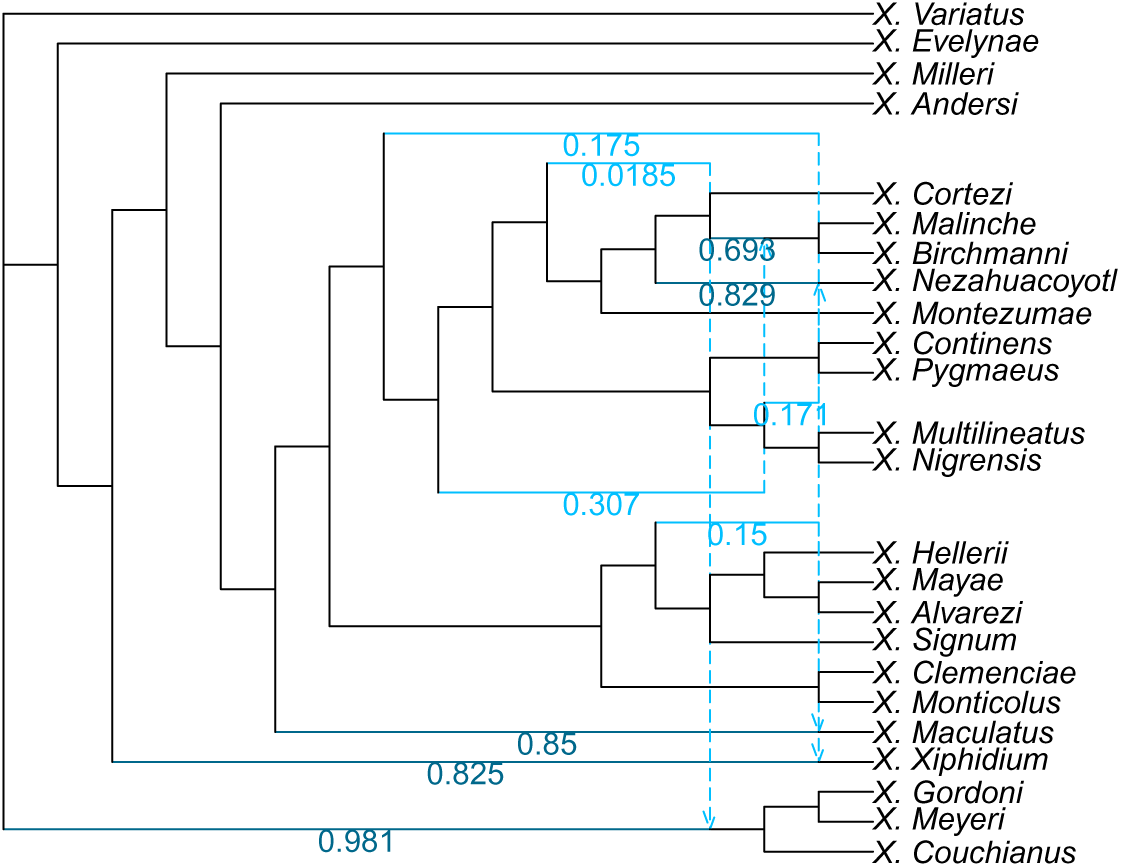
Clade-restricted (C-space) network inferred for the phylogeny of *Xiphophorus* (Poeciliidae) with 5 hybridizations.

**Fig. S48:**
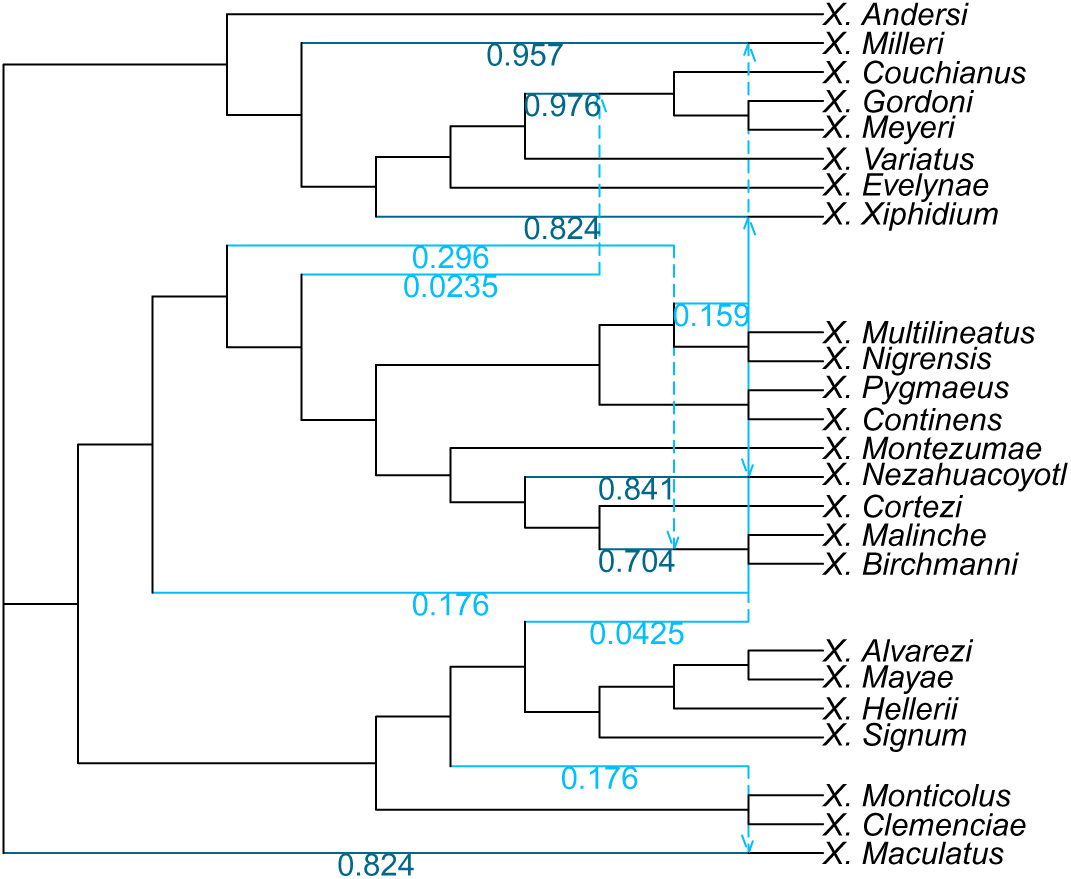
Clade-restricted (C-space) network inferred for the phylogeny of *Xiphophorus* (Poeciliidae) with 6 hybridizations.

**Fig. S49:**
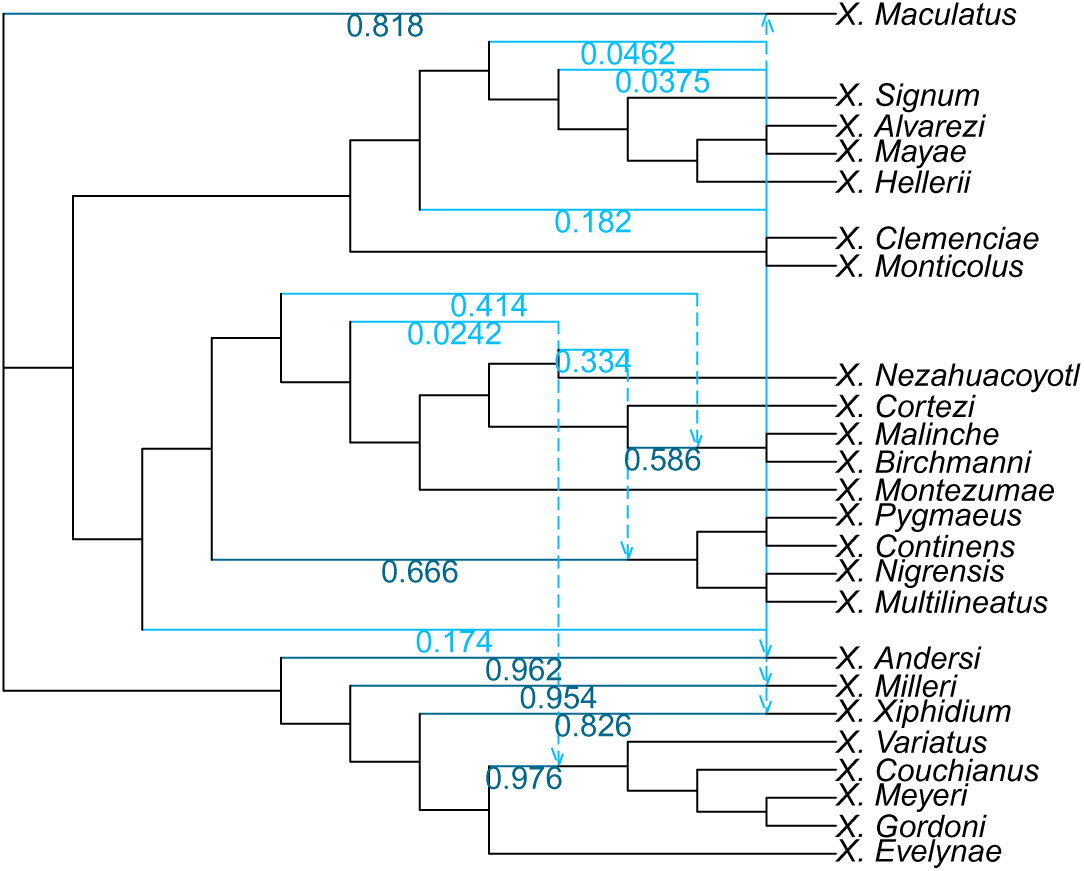
Clade-restricted (C-space) network inferred for the phylogeny of *Xiphophorus* (Poeciliidae) with 7 hybridizations.

#### C.3.6 CU-Space Networks

**Fig. S50:**
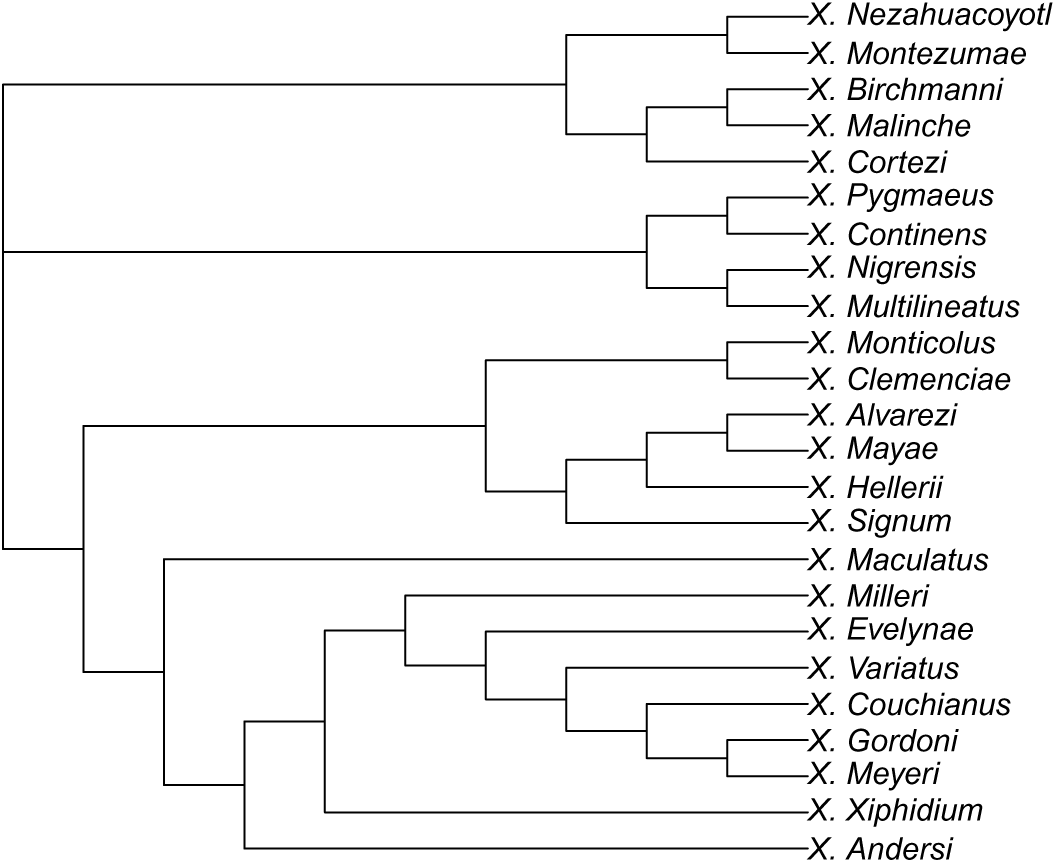
Clade-restricted (CU-space) network inferred for the phylogeny of *Xiphophorus* (Poeciliidae) with 0 hybridizations.

**Fig. S51:**
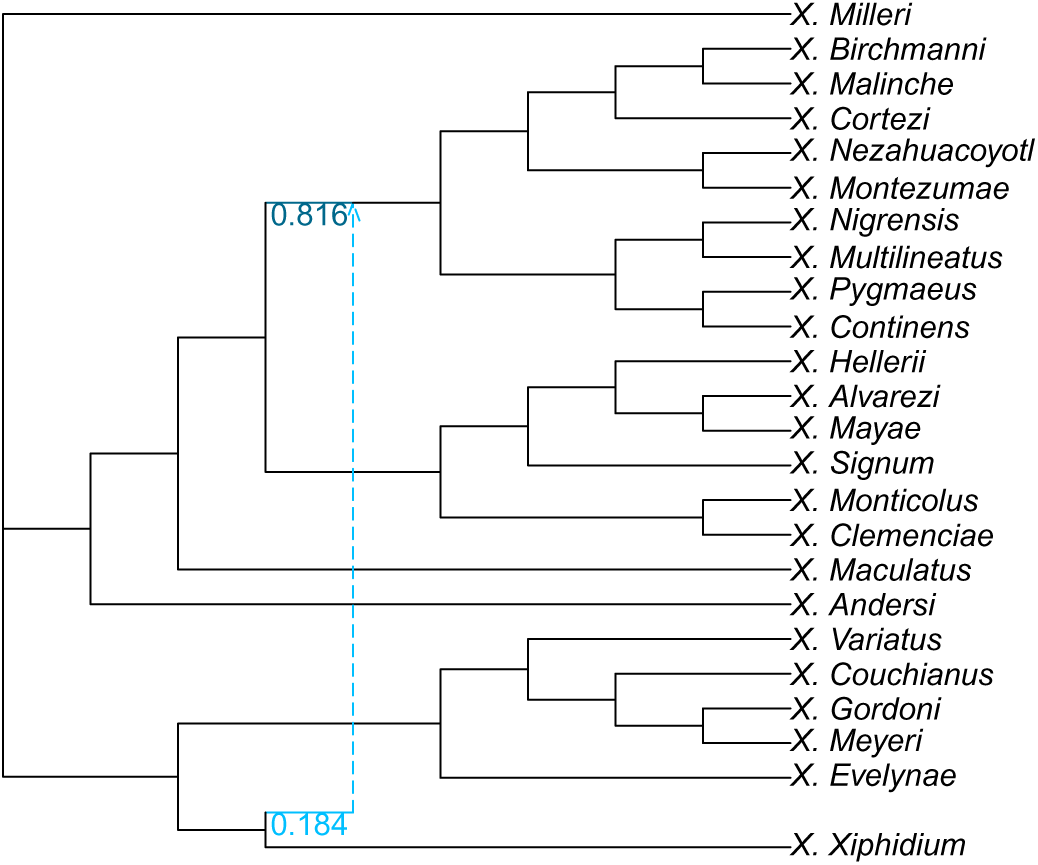
Clade-restricted (CU-space) network inferred for the phylogeny of *Xiphophorus* (Poeciliidae) with 1 hybridization.

**Fig. S52:**
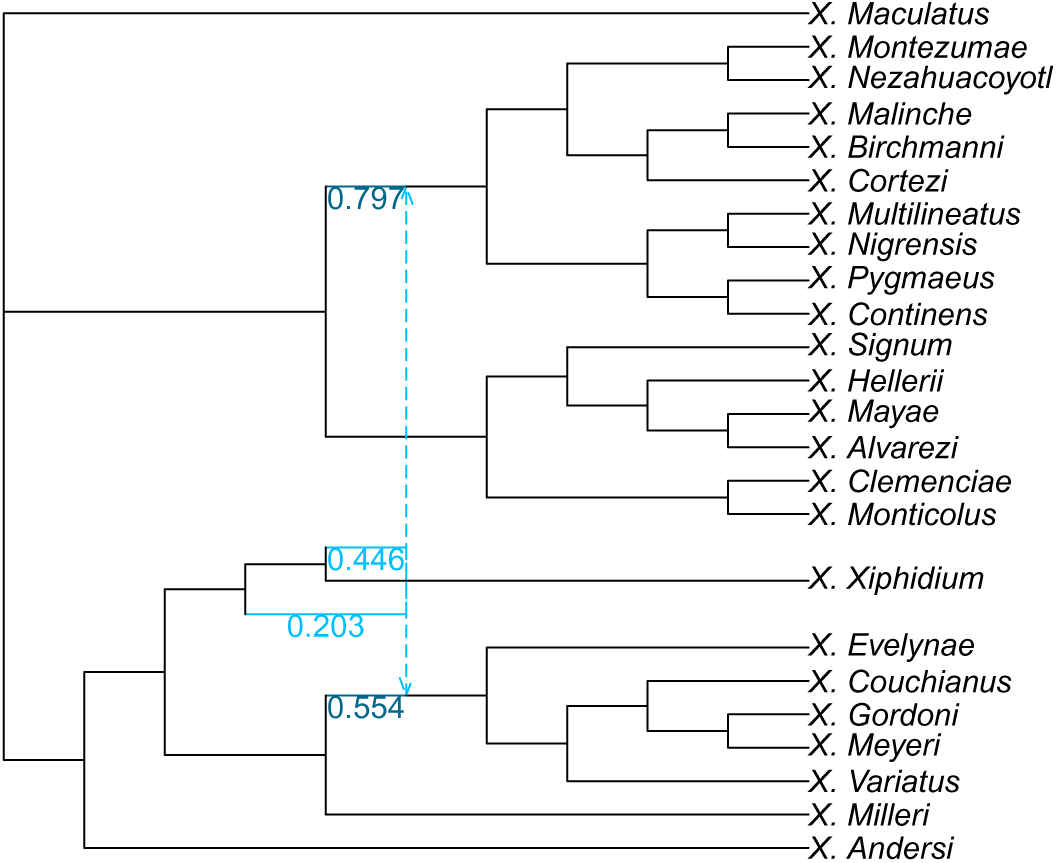
Clade-restricted (CU-space) network inferred for the phylogeny of *Xiphophorus* (Poeciliidae) with 2 hybridizations.

**Fig. S53:**
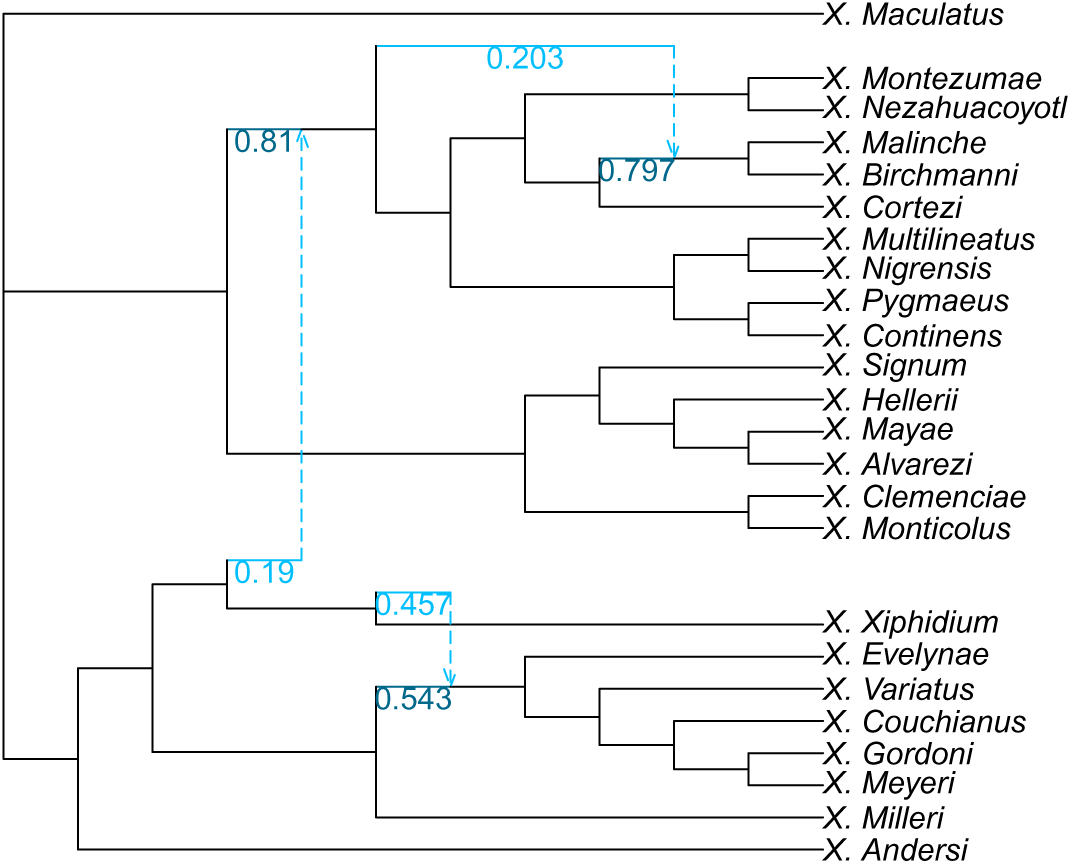
Clade-restricted (CU-space) network inferred for the phylogeny of *Xiphophorus* (Poeciliidae) with 3 hybridizations.

**Fig. S54:**
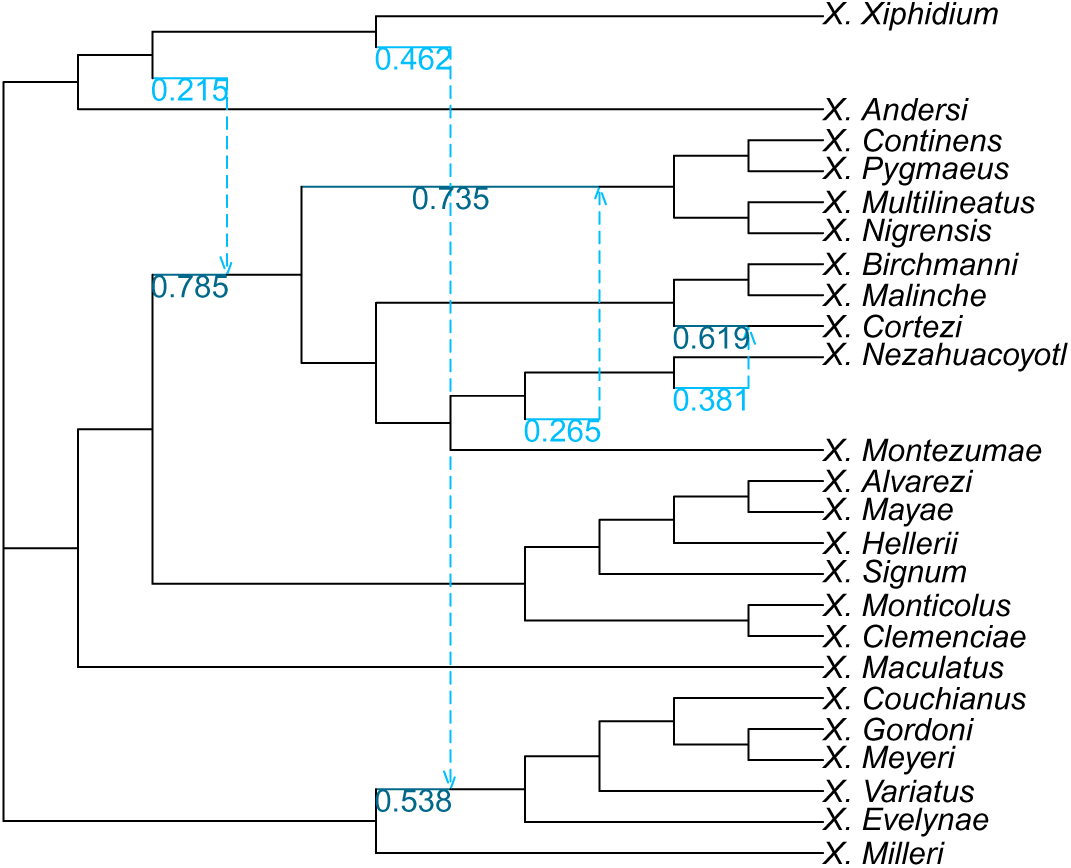
Clade-restricted (CU-space) network inferred for the phylogeny of *Xiphophorus* (Poeciliidae) with 4 hybridizations.

**Fig. S55:**
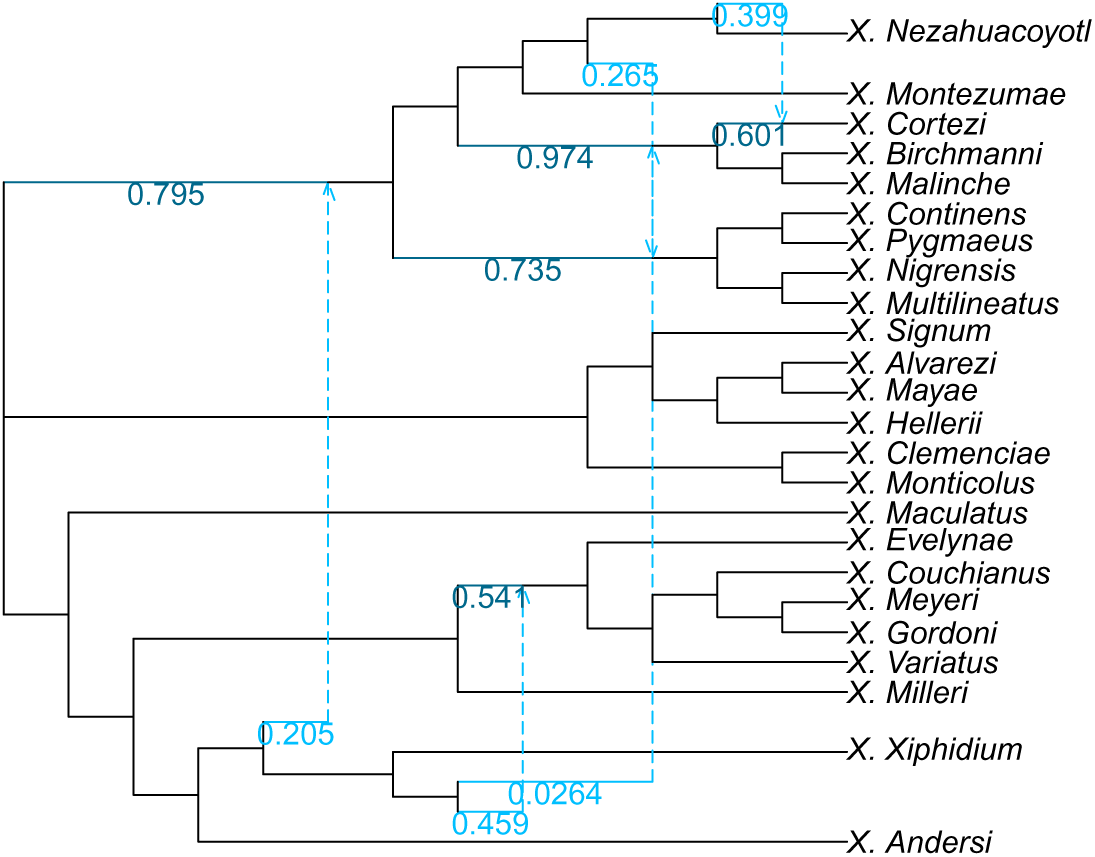
Clade-restricted (CU-space) network inferred for the phylogeny of *Xiphophorus* (Poeciliidae) with 5 hybridizations.

**Fig. S56:**
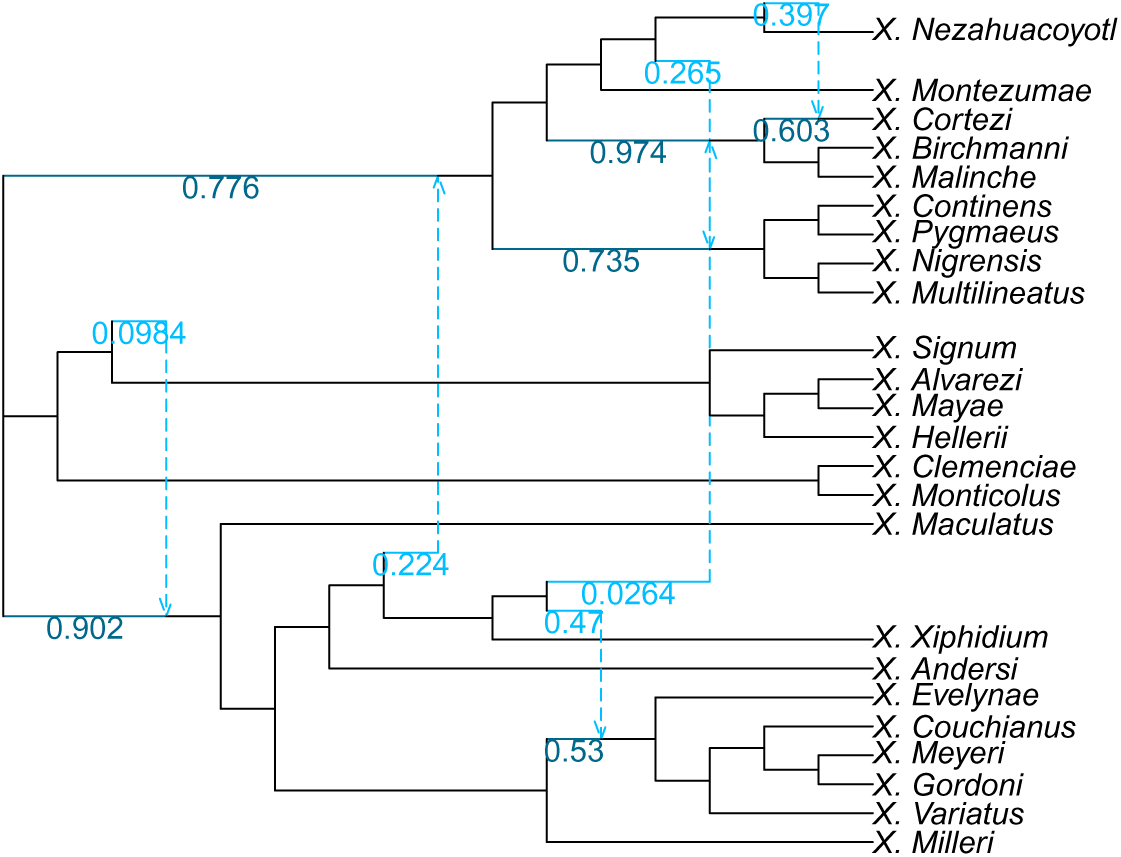
Clade-restricted (CU-space) network inferred for the phylogeny of *Xiphophorus* (Poeciliidae) with 6 hybridizations.

#### C.3.7 CUfast-Space Networks

**Fig. S57:**
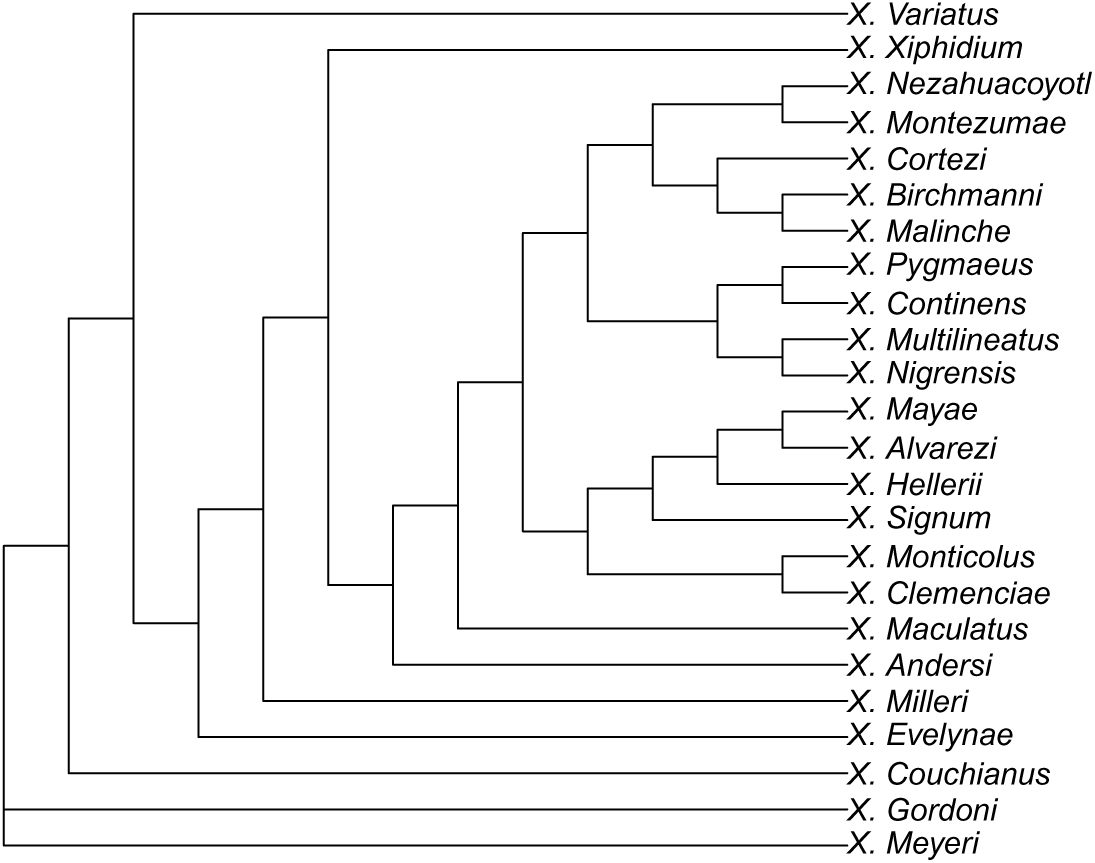
Clade-restricted (CUfast-space) network inferred for the phylogeny of *Xiphophorus* (Poeciliidae) with 0 hybridizations.

**Fig. S58:**
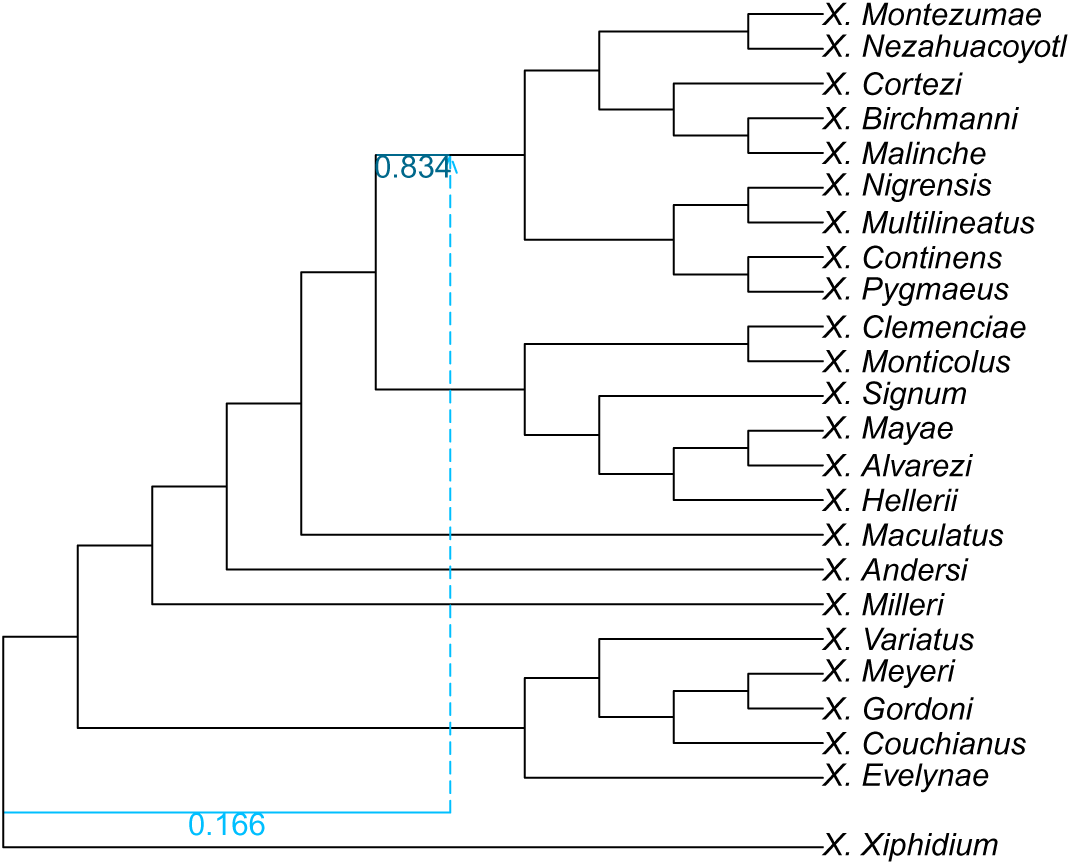
Clade-restricted (CUfast-space) network inferred for the phylogeny of *Xiphophorus* (Poeciliidae) with 1 hybridization.

**Fig. S59:**
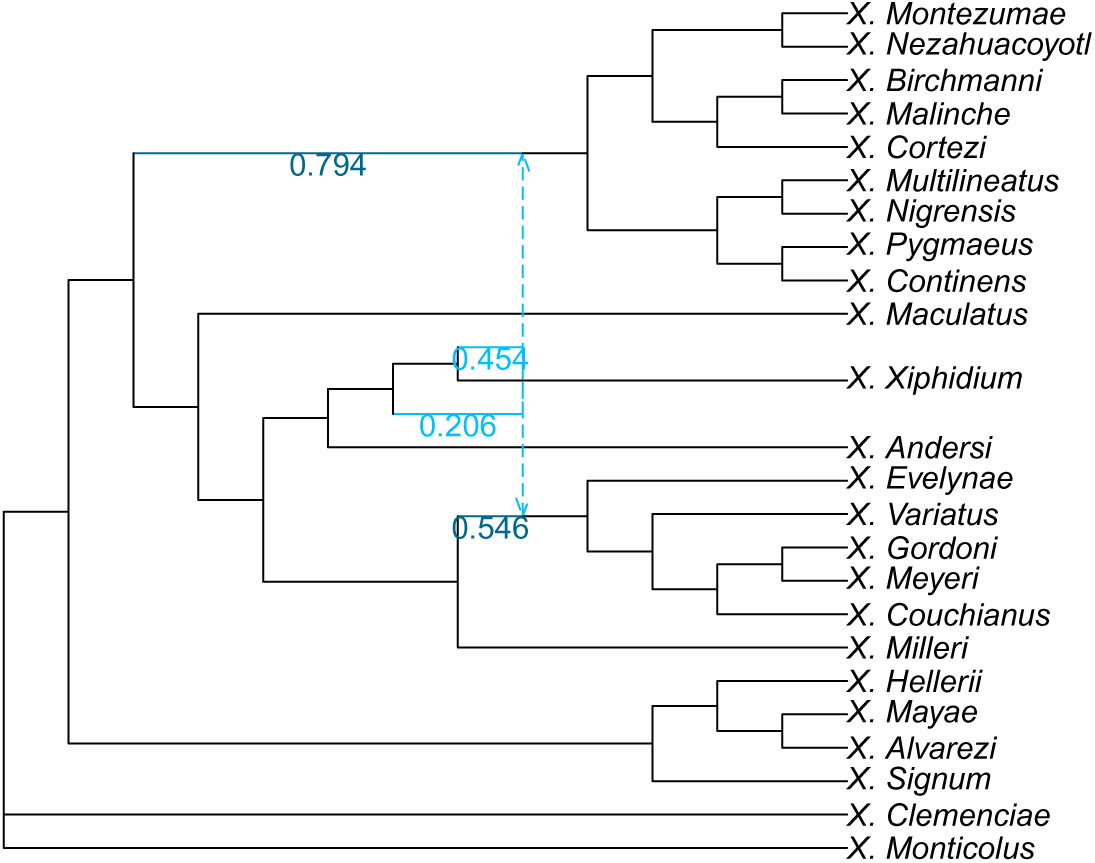
Clade-restricted (CUfast-space) network inferred for the phylogeny of *Xiphophorus* (Poeciliidae) with 2 hybridizations.

**Fig. S60:**
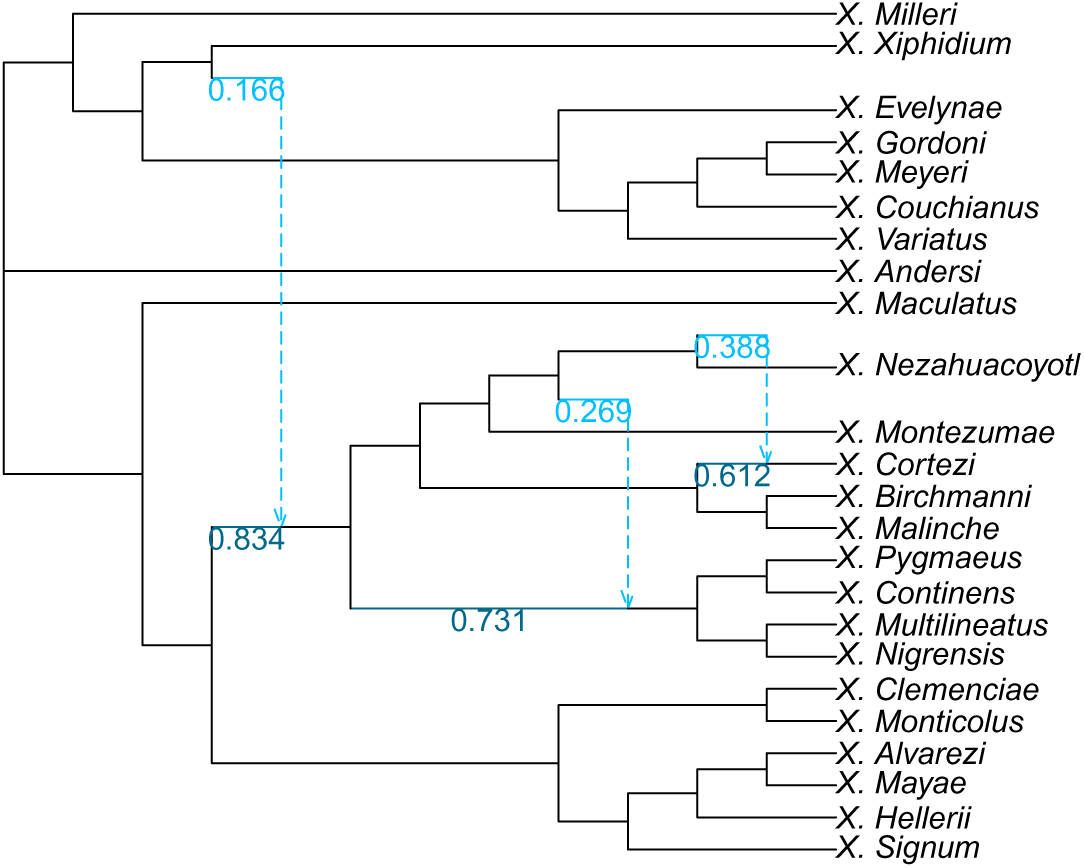
Clade-restricted (CUfast-space) network inferred for the phylogeny of *Xiphophorus* (Poeciliidae) with 3 hybridizations.

**Fig. S61:**
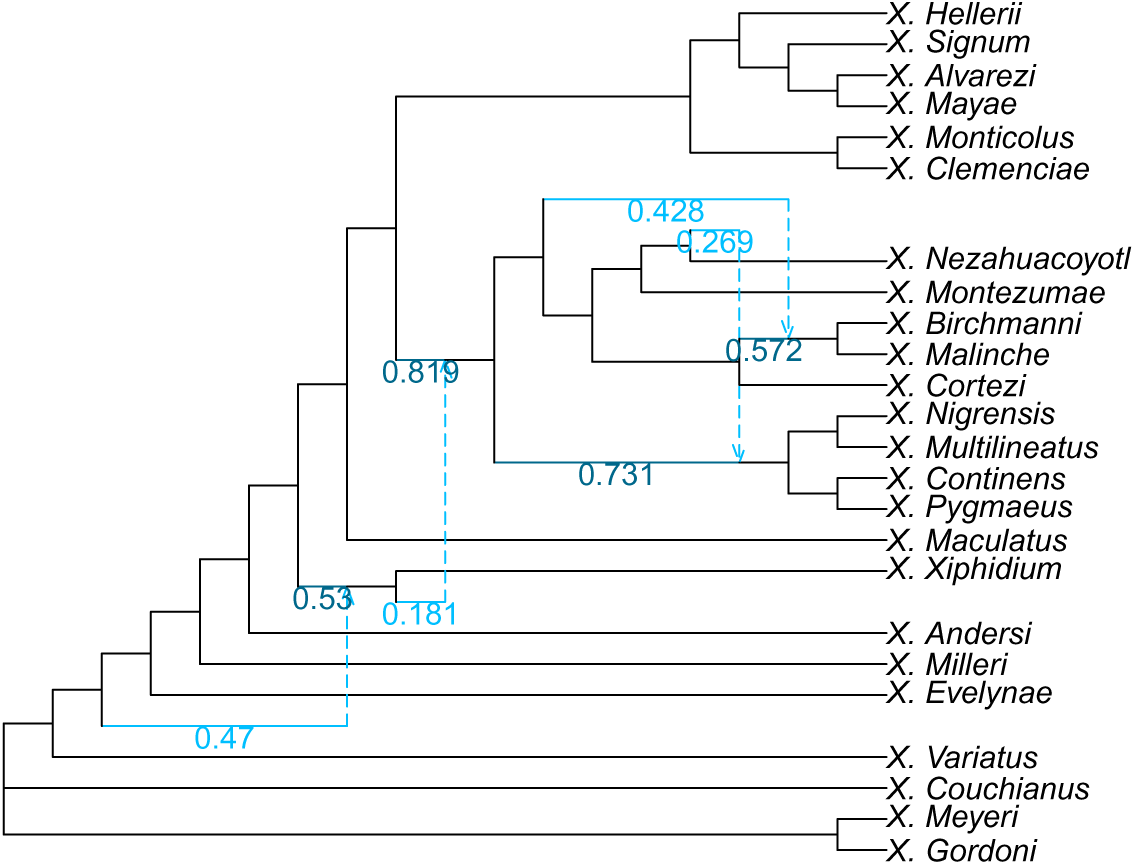
Clade-restricted (CUfast-space) network inferred for the phylogeny of *Xiphophorus* (Poeciliidae) with 4 hybridizations.

**Fig. S62:**
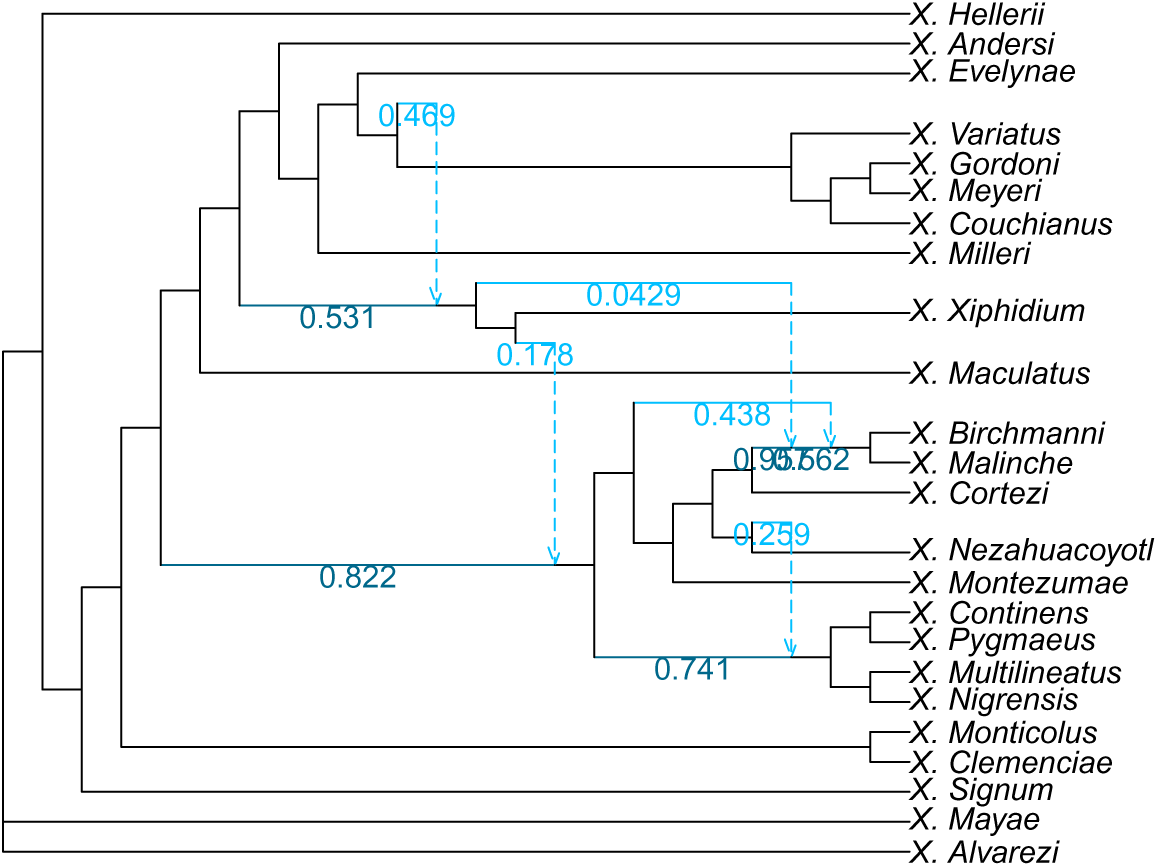
Clade-restricted (CUfast-space) network inferred for the phylogeny of *Xiphophorus* (Poeciliidae) with 5 hybridizations.

**Fig. S63:**
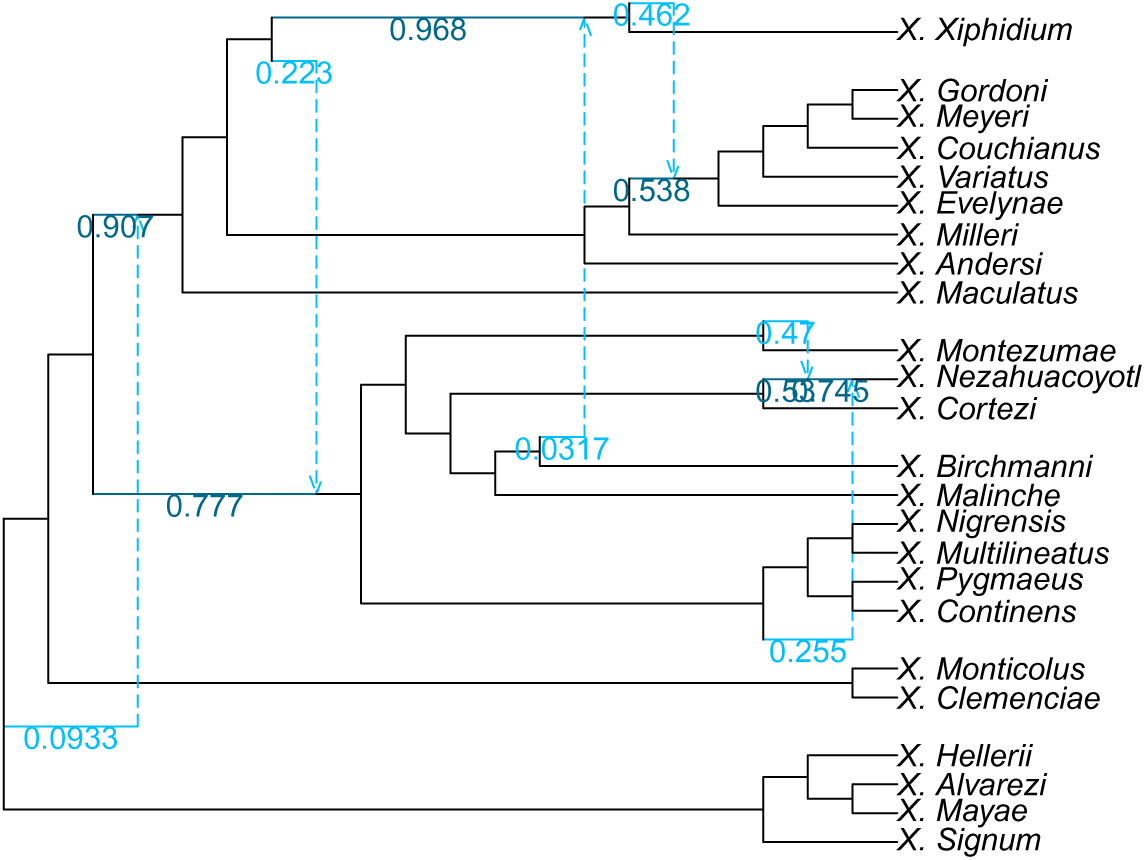
Clade-restricted (CUfast-space) network inferred for the phylogeny of *Xiphophorus* (Poeciliidae) with 6 hybridizations.

**Fig. S64:**
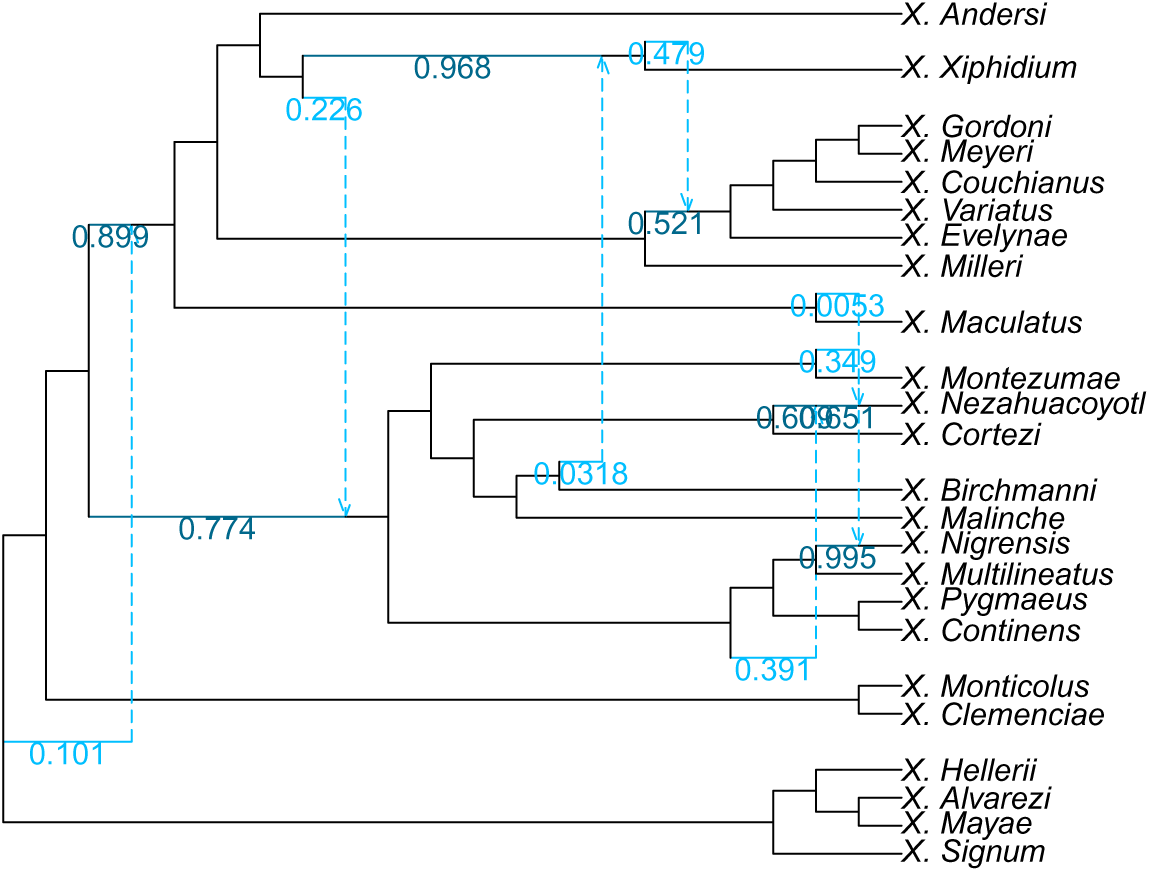
Clade-restricted (CUfast-space) network inferred for the phylogeny of *Xiphophorus* (Poeciliidae) with 7 hybridizations.

#### C.3.8 CUfast500-Space Networks

**Fig. S65:**
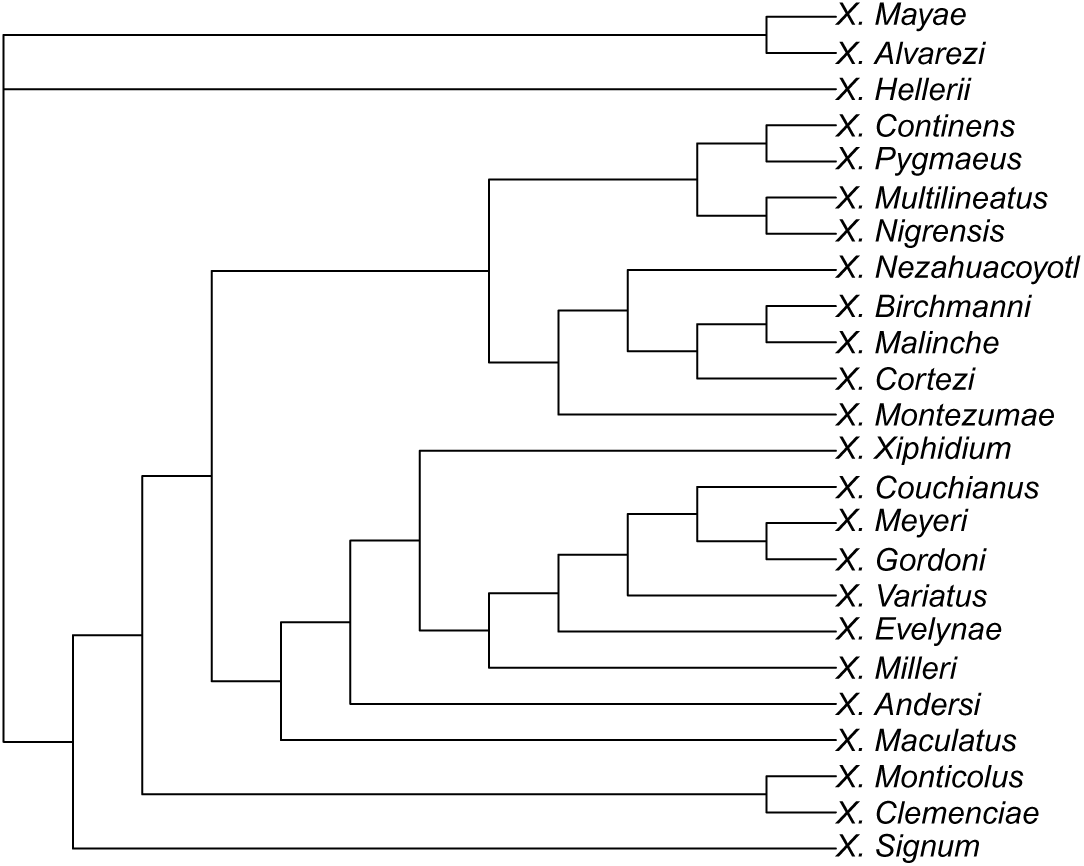
Clade-restricted (CUfast500-space) network inferred for the phylogeny of *Xiphophorus* (Poeciliidae) with 0 hybridizations.

**Fig. S66:**
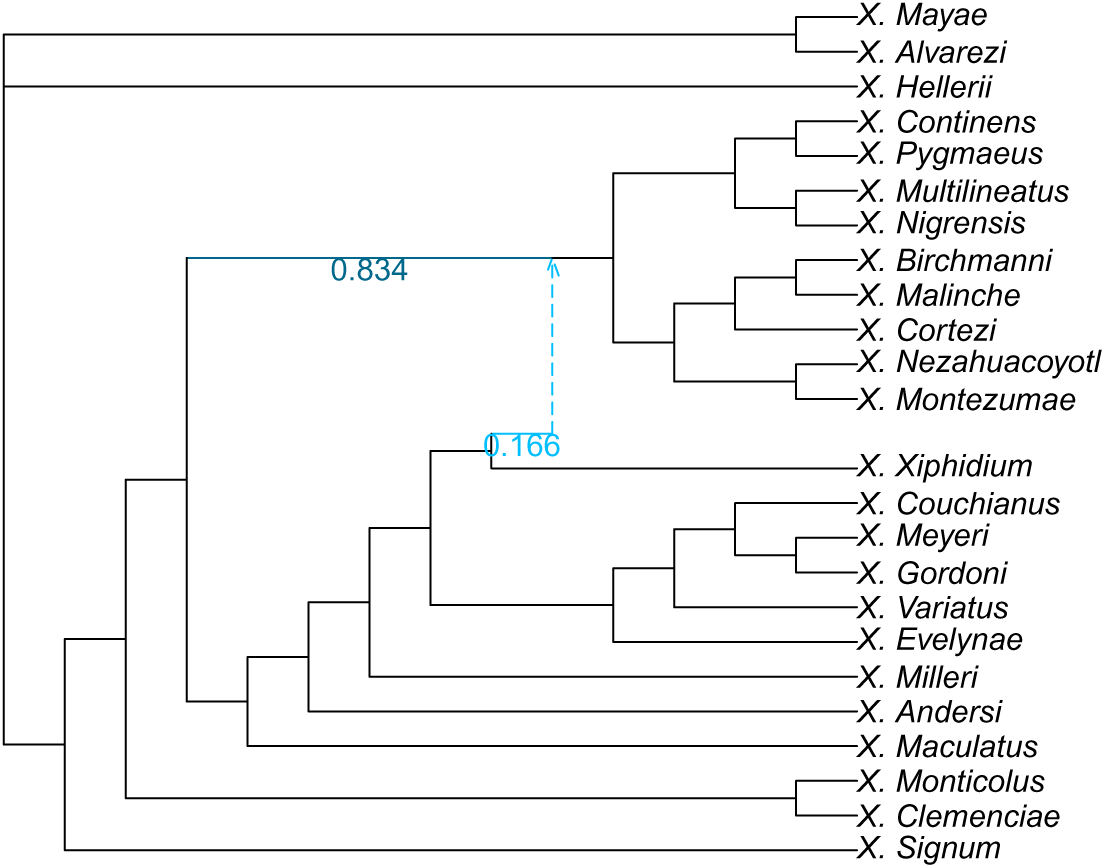
Clade-restricted (CUfast500-space) network inferred for the phylogeny of *Xiphophorus* (Poeciliidae) with 1 hybridization.

**Fig. S67:**
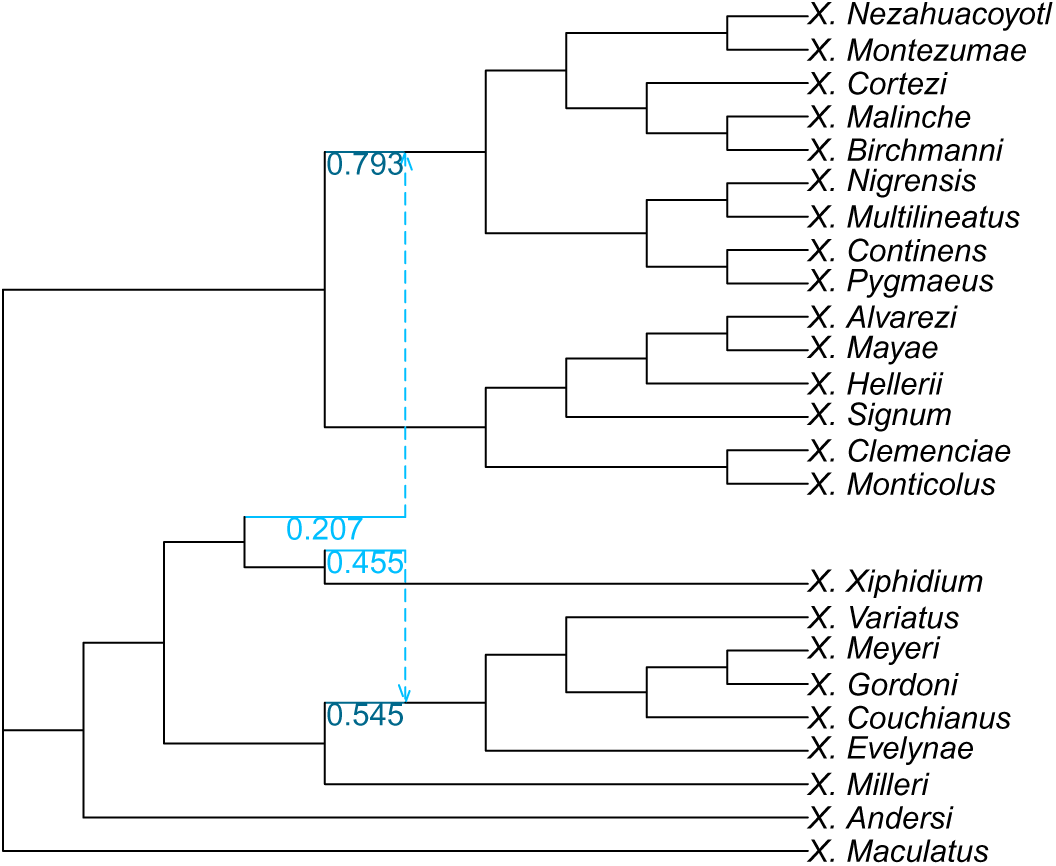
Clade-restricted (CUfast500-space) network inferred for the phylogeny of *Xiphophorus* (Poeciliidae) with 2 hybridizations.

**Fig. S68:**
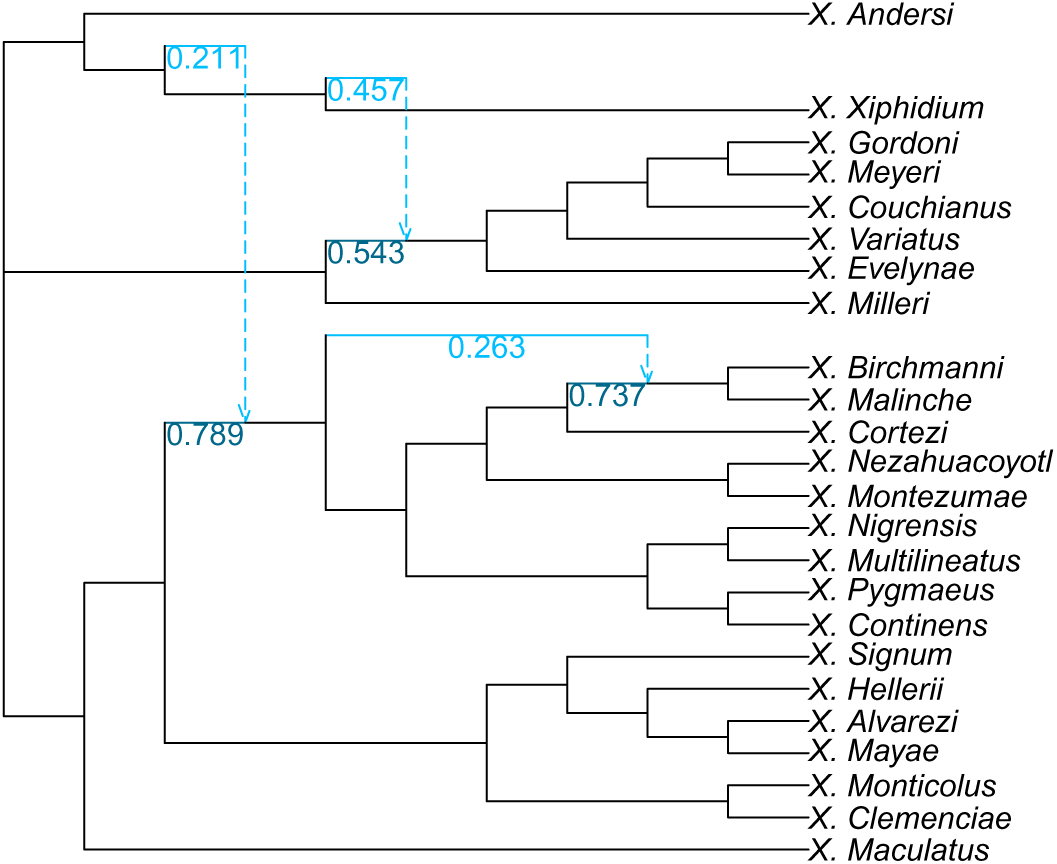
Clade-restricted (CUfast500-space) network inferred for the phylogeny of *Xiphophorus* (Poeciliidae) with 3 hybridizations.

**Fig. S69:**
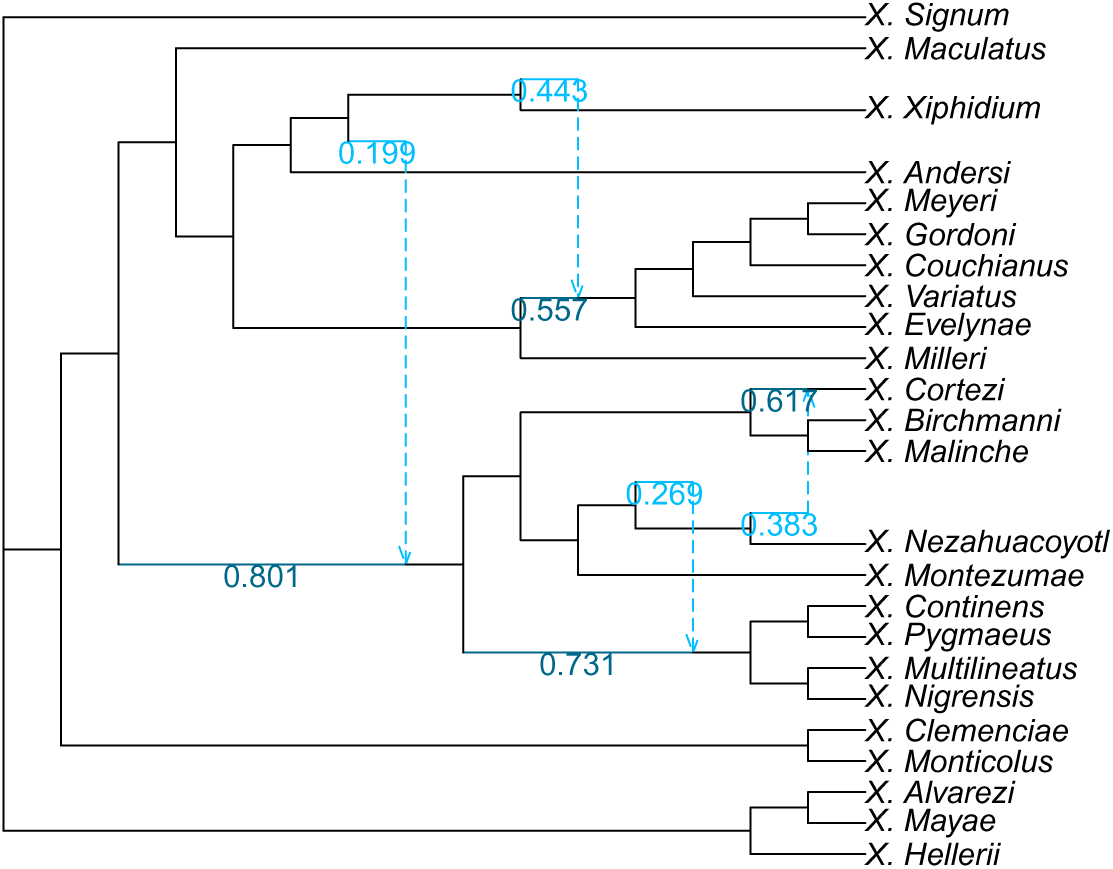
Clade-restricted (CUfast500-space) network inferred for the phylogeny of *Xiphophorus* (Poeciliidae) with 4 hybridizations.

**Fig. S70:**
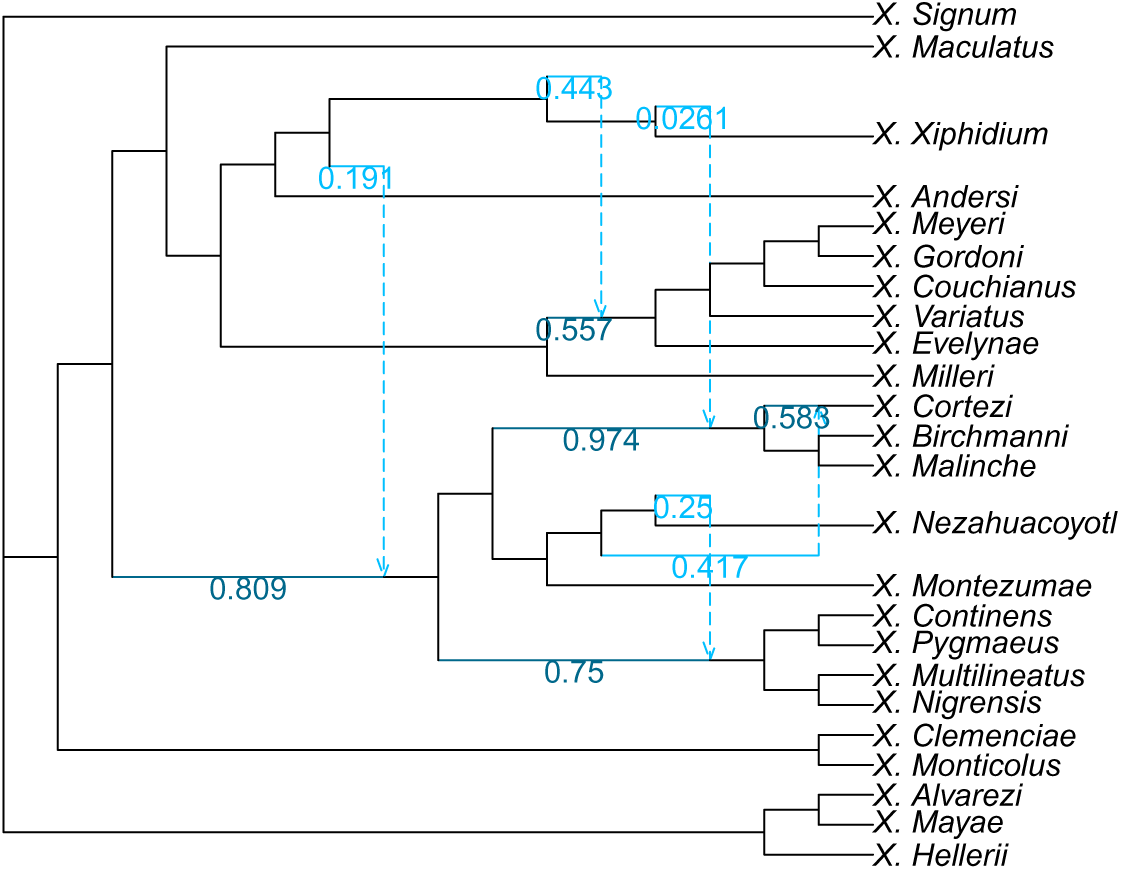
Clade-restricted (CUfast500-space) network inferred for the phylogeny of *Xiphophorus* (Poeciliidae) with 5 hybridizations.

**Fig. S71:**
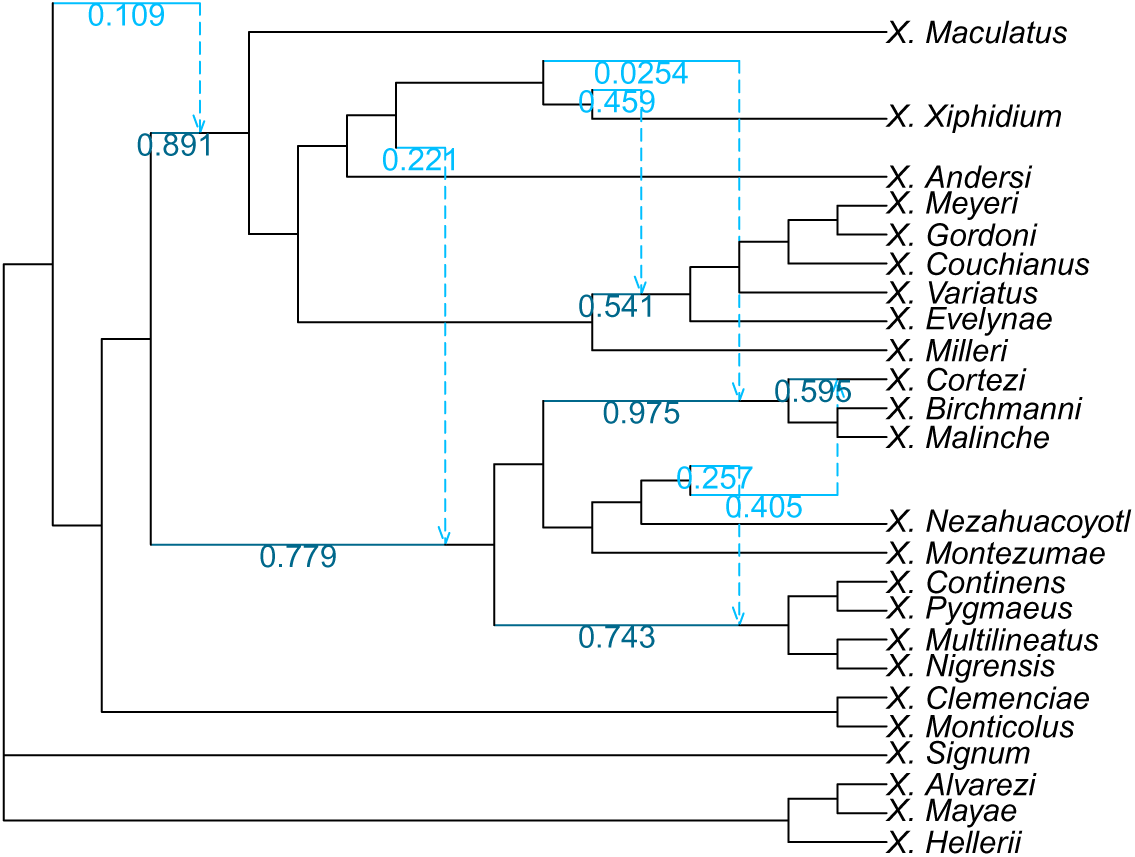
Clade-restricted (CUfast500-space) network inferred for the phylogeny of *Xiphophorus* (Poeciliidae) with 6 hybridizations.

**Fig. S72:**
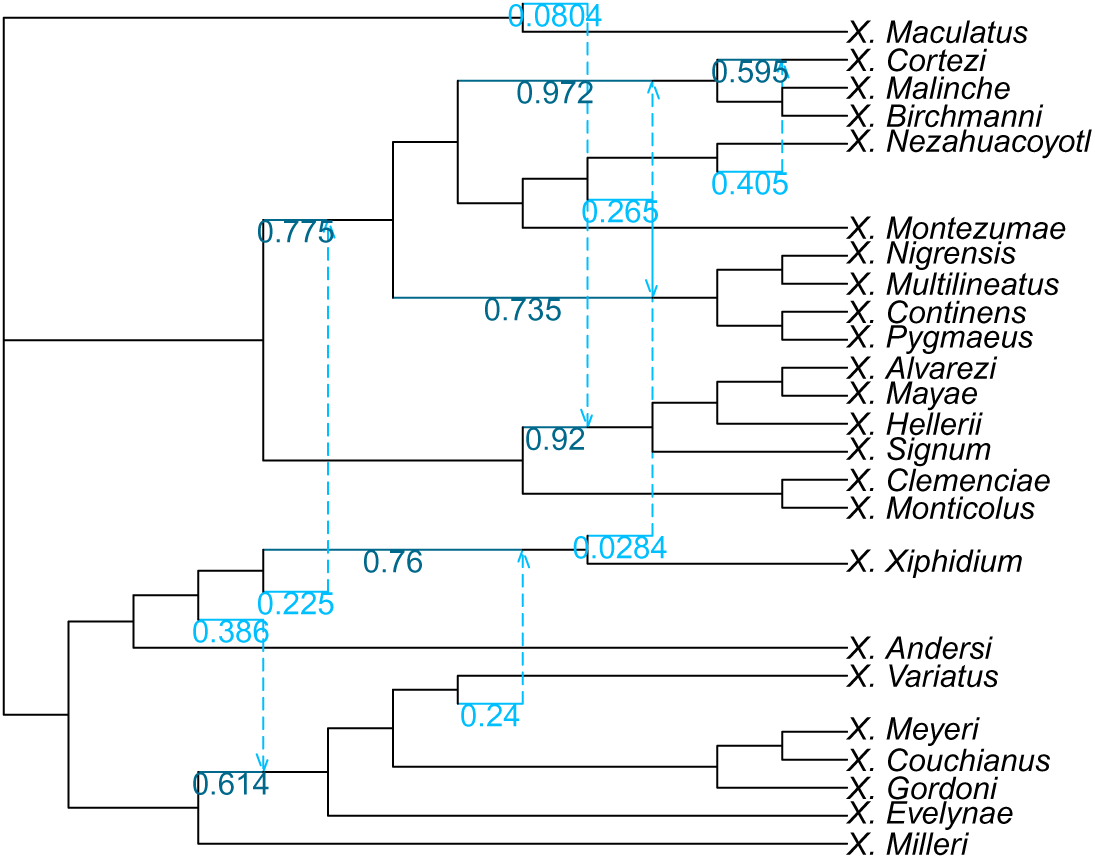
Clade-restricted (CUfast500-space) network inferred for the phylogeny of *Xiphophorus* (Poeciliidae) with 7 hybridizations.

## References

[1] Moret, B.M.E., Nakhleh, L., Warnow, T., Linder, C.R., Tholse, A., Padolina, A., Sun, J., Timme, R.: Phylogenetic networks: Modeling, reconstructibility, and accuracy 1(1), 13–23 10.1109/TCBB.2004.1017048405

[2] Kong, S., Solís-Lemus, C., Tiley, G.P.: Phylogenetic networks empower biodiversity research 122(31), 2410934122 10.1073/pnas.2410934122. Accessed 2025-11-03

[3] Steenwyk, J.L., Li, Y., Zhou, X., Shen, X.-X., Rokas, A.: Incongruence in the phylogenomics era 24(12), 834–850 (2023) 10.1038/s41576-023-00620-x. Accessed 2026-04-30

[4] Meng, C., Kubatko, L.S.: Detecting hybrid speciation in the presence of incomplete lineage sorting using gene tree incongruence: A model 75(1), 35–45 10.1016/j.tpb.2008.10.004. Accessed 2025-12-22

[5] Yu, Y., Degnan, J.H., Nakhleh, L.: The probability of a gene tree topology within a phylogenetic network with applications to hybridization detection. PLoS Genetics 8(4), 1002660 (2012) 10.1371/journal.pgen.1002660

[6] Degnan, J.H.: Modeling hybridization under the network multispecies coalescent. Systematic Biology 67(5), 786–799 (2018) 10.1093/sysbio/syy040

[7] Solís-Lemus, C., Ané, C.: Inferring Phylogenetic Networks with Maximum Pseu-dolikelihood under Incomplete Lineage Sorting. PLOS Genetics 12(3), 1005896 (2016) 10.1371/journal.pgen.1005896

[8] Hejase, H.A., Liu, K.J.: A scalability study of phylogenetic network inference methods using empirical datasets and simulations involving a single reticulation 17(1), 422 (2016) 10.1186/s12859-016-1277-1. Accessed 2026-04-29

[9] Yu, Y., Nakhleh, L.: A maximum pseudo-likelihood approach for phylogenetic networks. BMC genomics 16 Suppl 10(Suppl 10), 10 (2015) 10.1186/1471-2164-16-S10-S10

[10] Kong, S., Swofford, D.L., Kubatko, L.S.: Inference of Phylogenetic Networks From Sequence Data Using Composite Likelihood. Systematic Biology 74(1), 53–69 (2025) 10.1093/sysbio/syae054

[11] Blair, C., Ané, C.: Phylogenetic Trees and Networks Can Serve as Powerful and Complementary Approaches for Analysis of Genomic Data 69(3), 593–601 (2020) 10.1093/sysbio/syz056. Accessed 2026-04-30

[12] Kong, S., Pons, J.C., Kubatko, L., Wicke, K.: Classes of explicit phylogenetic networks and their biological and mathematical significance. Journal of Mathematical Biology 84(6), 47 (2022) 10.1007/s00285-022-01746-y

[13] Holtgrefe, N., Huber, K.T., family=Iersel, p.u. given=Leo, Jones, M., Martin, S., Moulton, V.: Squirrel: Reconstructing semi-directed phylogenetic level-1 networks from four-leaved networks or sequence alignments, 067 (2025) 10.1093/molbev/msaf067

[14] Allman, E.S., Baños, H., Rhodes, J.A., Wicke, K.: NANUQ+: A divide-and-conquer approach to network estimation. Algorithms for Molecular Biology: AMB 20, 14 (2025) 10.1186/s13015-025-00274-w

[15] Allman, E.S., Baños, H., Garrote-Lopez, M., Rhodes, J.A.: Identifiability of Level-1 Species Networks from Gene Tree Quartets 86(9), 110 (2024) 10.1007/s11538-024-01339-4. Accessed 2026-06-05

[16] Ané, C., Fogg, J., Allman, E.S., Baños, H., Rhodes, J.A.: Anomalous networks under the multispecies coalescent: Theory and prevalence 88(3), 29 10.1007/s00285-024-02050-7. Accessed 2024-08-29

[17] Allman, E.S., Ané, C., Baños, H., Rhodes, J.A.: Beyond Level-1: Identifiability of a Class of Galled Tree-Child Networks 87(11), 166 10.1007/s11538-025-01545-841123850. Accessed 2025-11-03

[18] Cardona, G., Rossello, F., Valiente, G.: Comparison of Tree-Child Phylogenetic Networks. IEEE/ACM Transactions on Computational Biology and Bioinformatics 6(4), 552–569 (2009) 10.1109/TCBB.2007.70270

[19] Huson, D.H., Klöpper, T.H.: Beyond Galled Trees - Decomposition and Computation of Galled Networks. In: Speed, T., Huang, H. (eds.) Research in Computational Molecular Biology, pp. 211–225. Springer, Berlin, Heidelberg (2007). 10.1007/978-3-540-71681-5_15

[20] Kolbow, N., Kong, S., Chafin, T., Justison, J., Ané, C., Solís-Lemus, C.: SNaQ.Jl: Improved Scalability for Phylogenetic Network Inference. 10.1101/2025.11.17.688917. https://www.biorxiv.org/content/10.1101/2025.11.17.688917v1 Accessed 2026-04-17

[21] Du, K., Ricci, J.M.B., Lu, Y., Garcia-Olazabal, M., Walter, R.B., Warren, W.C., Dodge, T.O., Schumer, M., Park, H., Meyer, A., Schartl, M.: Phylogenomic analyses of all species of swordtail fishes (genus Xiphophorus) show that hybridization preceded speciation. Nature Communications 15(1), 6609 (2024) 10.1038/s41467-024-50852-6

[22] Cui, R., Schumer, M., Kruesi, K., Walter, R., Andolfatto, P., Rosenthal, G.G.: PHYLOGENOMICS REVEALS EXTENSIVE RETICULATE EVOLUTION IN XIPHOPHORUS FISHES. Evolution 67(8), 2166–2179 (2013) 10.1111/evo.12099

[23] Rosenthal, G.G., Reyna, X.F.d.l.R., Kazianis, S., Stephens, M.J., Morizot, D.C., Ryan, M.J., León, F.J.G.: Dissolution of Sexual Signal Complexes in a Hybrid Zone between the Swordtails Xiphophorus birchmanni and Xiphophorus malinche (Poeciliidae). Copeia 2003(2), 299–307 (2003) 10.1643/0045-8511(2003)003[0299:DOSSCI]2.0.CO;2

[24] Schumer, M., Cui, R., Powell, D.L., Rosenthal, G.G., Andolfatto, P.: Ancient hybridization and genomic stabilization in a swordtail fish. Molecular Ecology 25(11), 2661–2679 (2016) 10.1111/mec.13602

[25] Schartl, M., Walter, R.B., Shen, Y., Garcia, T., Catchen, J., Amores, A., Braasch, I., Chalopin, D., Volff, J.-N., Lesch, K.-P., Bisazza, A., Minx, P., Hillier, L., Wilson, R.K., Fuerstenberg, S., Boore, J., Searle, S., Postlethwait, J.H., Warren, W.C.: The genome of the platyfish, Xiphophorus maculatus, provides insights into evolutionary adaptation and several complex traits 45(5), 567–572 (2013) 10.1038/ng.2604. Accessed 2026-03-18

[26] Meyer, A.: The evolution of sexually selected traits in male swordtail fishes (Xiphophorus: Poeciliidae) 79(3), 329–337 (1997) 10.1038/hdy.1997.161. Accessed 2026-03-18

[27] Powell, M.J.D.: The BOBYQA algorithm for bound constrained optimization without derivatives. Cambridge NA Report NA2009/06, Department of Applied Mathematics and Theoretical Physics, University of Cambridge (2009)

[28] Nocedal, J.: Updating quasi-Newton matrices with limited storage. Mathematics of Computation 35(151), 773–782 (1980) 10.1090/S0025-5718-1980-0572855-7

[29] García Fernández, J., Ahmad, N., Gerven, M.: A unified perspective on optimization in machine learning and neuroscience: From gradient descent to neural adaptation. arXiv e-prints, 2510 (2025)

[30] Fogg, J., Allman, E.S., Ané, C.: PhyloCoalSimulations: A Simulator for Network Multispecies Coalescent Models, Including a New Extension for the Inheritance of Gene Flow. Systematic Biology 72(5), 1171–1179 (2023) 10.1093/sysbio/syad030

[31] Justison, J.A., Solis-Lemus, C., Heath, T.A.: SiPhyNetwork: An R package for simulating phylogenetic networks. Methods in Ecology and Evolution 14(7), 1687–1698 (2023) 10.1111/2041-210X.14116

[32] Rambaut, A., Grass, N.C.: Seq-Gen: An application for the Monte Carlo simulation of DNA sequence evolution along phylogenetic trees. Bioinformatics 13(3), 235–238 (1997) 10.1093/bioinformatics/13.3.235

[33] Hasegawa, M., Kishino, H., Yano, T.-a.: Dating of the human-ape splitting by a molecular clock of mitochondrial DNA. Journal of Molecular Evolution 22(2), 160–174 (1985) 10.1007/BF02101694

[34] Wong, T.K.F., Ly-Trong, N., Ren, H., Baños, H., Roger, A.J., Susko, E., Bielow, C., Maio, N.D., Goldman, N., Hahn, M.W., Huttley, G., Lanfear, R., Minh, B.Q.: IQ-TREE 3: Phylogenomic Inference Software using Complex Evolutionary Models (2025)

[35] Huson, D.H., Rupp, R., Scornavacca, C.: Phylogenetic Networks: Concepts, Algorithms and Applications. Cambridge University Press, NY (2010)

[36] Zhang, C., Rabiee, M., Sayyari, E., Mirarab, S.: ASTRAL-III: Polynomial time species tree reconstruction from partially resolved gene trees. BMC Bioinformatics 19(6), 153 (2018) 10.1186/s12859-018-2129-y

[37] Ronquist, F., Huelsenbeck, J.P.: Mrbayes 3: Bayesian phylogenetic inference under mixed models. Bioinformatics 19(12), 1572–1574 (2003) 10.1093/bioinformatics/btg180

[38] Baudry, J.-P., Maugis, C., Michel, B.: Slope heuristics: Overview and implementation. Statistics and Computing 22(2), 455–470 (2012) 10.1007/s11222-011-9236-1

[39] Meyer, A., Salzburger, W., Schartl, M.: Hybrid origin of a swordtail species (Teleostei: Xiphophorus clemenciae) driven by sexual selection. Molecular Ecology 15(3), 721–730 (2006) 10.1111/j.1365-294X.2006.02810.x

[40] Oroian, C., Dăescu, A.M.: Unveiling hybridization-driven speciation: Insights from the genomic evolution of Xiphophorus 14(1) (2024)

[41] Preising, G.A., Gunn, T., Baczenas, J.J., Powell, D.L., Dodge, T.O., Sewell, S.T., Pollock, A., Machin Kairuz, J.A., Savage, M., Lu, Y., Fitschen-Brown, M., Meyer, A., Schartl, M., Cummings, M., Thakur, S., Inman, C.M., Ríos-Cardenas, O., Morris, M., Tobler, M., Schumer, M.: Recurrent evolution of small body size and loss of the sword ornament in Northern swordtail fish. Evolution 78(12), 2017–2031 (2024) 10.1093/evolut/qpae124

[42] Allman, E.S., Ane, C., Banos, H., Rhodes, J.A.: Beyond Level-1: Identifiability of a Class of Galled Tree-Child Networks. 10.48550/arXiv.2504.21116. http://arxiv.org/abs/2504.21116 Accessed 2026-06-05

[43] Englander, A.K., Frohn, M., Gross, E., Holtgrefe, N., Iersel, L.V., Jones, M., Sulli-vant, S.: Identifiability of Phylogenetic Level-2 Networks Under the Jukes-Cantor Model. 10.1101/2025.04.18.649493. https://www.biorxiv.org/content/10.1101/2025.04.18.649493v3 Accessed 2026-06-05

[44] Holtgrefe, N., Allman, E.S., Baños, H., family=Iersel, p.u. given=Leo, Moulton, V., Rhodes, J.A., Wicke, K.: Distinguishing Phylogenetic Level-2 Networks with Quartets and Inter-Taxon Quartet Distances 87(12), 168 (2025) 10.1007/s11538-025-01549-4. Accessed 2026-06-05

[45] Solís-Lemus, C., Bastide, P., Ané, C.: PhyloNetworks: A Package for Phylogenetic Networks. Molecular Biology and Evolution 34(12), 3292–3298 (2017) 10.1093/molbev/msx235

